# An Updated Polygenic Index Repository: Expanded Phenotypes, New Cohorts, and Improved Causal Inference

**DOI:** 10.1101/2025.05.14.653986

**Authors:** Robel Alemu, Anastasia Terskaya, Matthew Howell, Junming Guan, Harry Sands, Aaron Kleinman, 23andMe Research Team, David Bann, Tim Morris, George B. Ploubidis, Emla Fitzsimons, Kathleen Mullan Harris, Avshalom Caspi, David L. Corcoran, Terrie E. Moffitt, Richie Poulton, Karen Sugden, Benjamin S. Williams, Andrew Steptoe, Olesya Ajnakina, Uku Vainik, Tõnu Esko, Estonian Biobank Research Team, Archie Campbell, Caroline Hayward, William G. Iacono, Matt McGue, Robert F. Krueger, Anna R. Docherty, Andrey A. Shabalin, Ralph Hertwig, Philipp Koellinger, David Richter, Jan Goebel, Rafael Ahlskog, Sven Oskarsson, Patrik K.E. Magnusson, K. Paige Harden, Elliot M. Tucker-Drob, Charlotte K. L. Pahnke, Carlo Maj, Frank M. Spinath, Pamela Herd, Jeremy Freese, David Laibson, Michelle N. Meyer, Jonathan Jala, David Cesarini, Alexander Strudwick Young, Patrick Turley, Daniel J. Benjamin, Aysu Okbay

## Abstract

Polygenic indexes (PGIs) — DNA-based predictors of individual phenotypes — have become essential tools across biomedical and social sciences. We introduce Version 2 of the Polygenic Index Repository, which expands phenotype coverage from 47 to 61, increases the number of participating datasets from 11 to 20, and adopts a more consistent and improved methodology for PGI construction. For 16 phenotypes, we leverage summary statistics from an updated GWAS meta-analysis with greater statistical power compared to the original release, thereby improving the PGI’s predictive power. To improve power for family-based analyses, we provide imputed parental PGIs in all datasets with first-degree relatives and offer a framework for interpreting results from analyses that control for parental PGIs. We illustrate the utility of parental PGIs using two applications: (1) comparing PGI associations with and without parental PGI controls for all phenotypes in two Repository datasets with family data, and (2) for BMI and diastolic blood pressure, exploring the contribution of causal versus non-causal components of PGI associations to the imperfect portability of PGIs across subgroups within a genetic ancestry. Collectively, the updates enhance predictive performance, broaden the Repository’s scope, and introduce novel resources that reduce confounding bias and improve interpretability.

---

Polygenic indexes (PGIs, often called “polygenic scores” or “polygenic risk scores”) are DNA-based phenotype predictors. A PGI is constructed as a weighted sum of allele counts, often from millions of genome-wide single-nucleotide polymorphisms (SNPs). The weights are obtained from “genome-wide association studies” (GWAS), hypothesis-free studies that test the association between a phenotype and each of the millions of SNPs, one at a time. PGIs have become a major tool for genetics research as GWAS discovery samples have grown, enabling increasingly precise estimation of the SNP weights and thereby increasingly predictive PGIs. In medicine, the focus has been on risk stratification with potential clinical applications^1,2^. In psychology, epidemiology, sociology, and economics, PGIs have been used in a variety of creative ways, with much of the work examining the interplay between genetic and environmental factors ^3–6^.

In a previous paper^7^, we introduced the Polygenic Index Repository as a resource to facilitate research using PGIs. To maximize predictive power, each PGI was constructed using a combined GWAS meta-analysis that aggregated summary statistics from multiple sources, often including data not publicly available. By centralizing this process—both the aggregation of GWAS results and the construction of PGIs—the Repository spares individual researchers from navigating complex data access procedures and conducting the analytical work needed to construct PGIs on a per-project basis. The dissemination of high-quality PGIs built with a uniform methodology in the first version of the Repository lowered barriers to entry, improved predictive power, and enhanced comparability across PGI-based studies. The original release also introduced a theoretical framework to aid interpretation, treating each PGI as a noisy proxy for the “additive SNP factor”—that is, the PGI that would be obtained if the true SNP effects were known. The framework also motivated a measurement-error adjustment that can be applied when the PGI is used as a proxy for the additive SNP factor (which is unobserved, since weights are always estimated with some error).

The updates to the Repository described in this paper fall into two broad categories: (i) improvements that expand its scope and increase PGIs’ predictive power and (ii) new tools designed to support within-family analyses, which enable more robust causal inference.

In the first category, our new release extends the number of phenotypes covered from 47 to 61, adding eight biomarkers, eight health-related phenotypes, five psychiatric conditions, and one skill^a^ (Table 1). It also increases the number of participating datasets from 11 to 20, incorporating several large longitudinal studies and biobanks that were not included in the original release (Table 2). All PGIs are constructed using a new, improved methodology for generating SNP weights, and for 16 out of 47 of the first-release phenotypes, we were able to incorporate new data into the GWAS meta-analysis. These methodological and data upgrades substantially boost PGI predictive power—enabling them to capture more phenotypic variance and increasing their utility for research. We showcase the predictive power of the PGIs in three repository cohorts: the Health and Retirement Study, Wisconsin Longitudinal Study (WLS), and UK Biobank (UKB). With broader phenotype coverage and improved accuracy, the new release offers a more powerful and comprehensive resource for studying genetic influences across diverse outcomes. As in the original release, we distribute the PGIs as ready-to-use variables through participating data providers, thereby removing technical and administrative barriers that researchers may face.

**Table 1.**
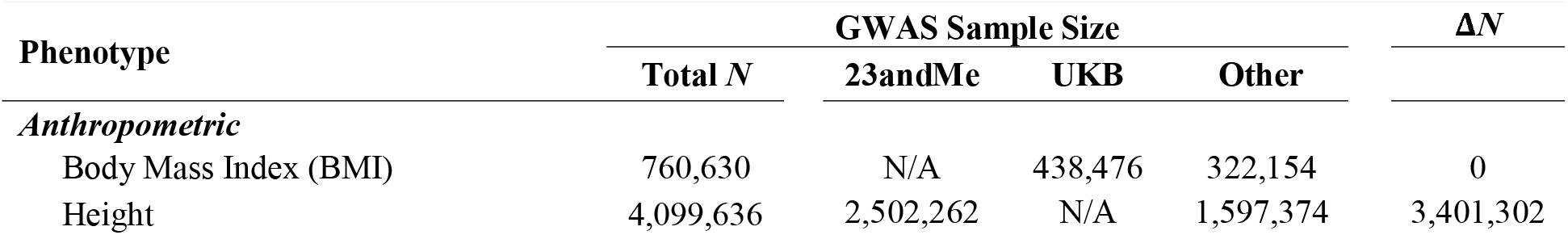
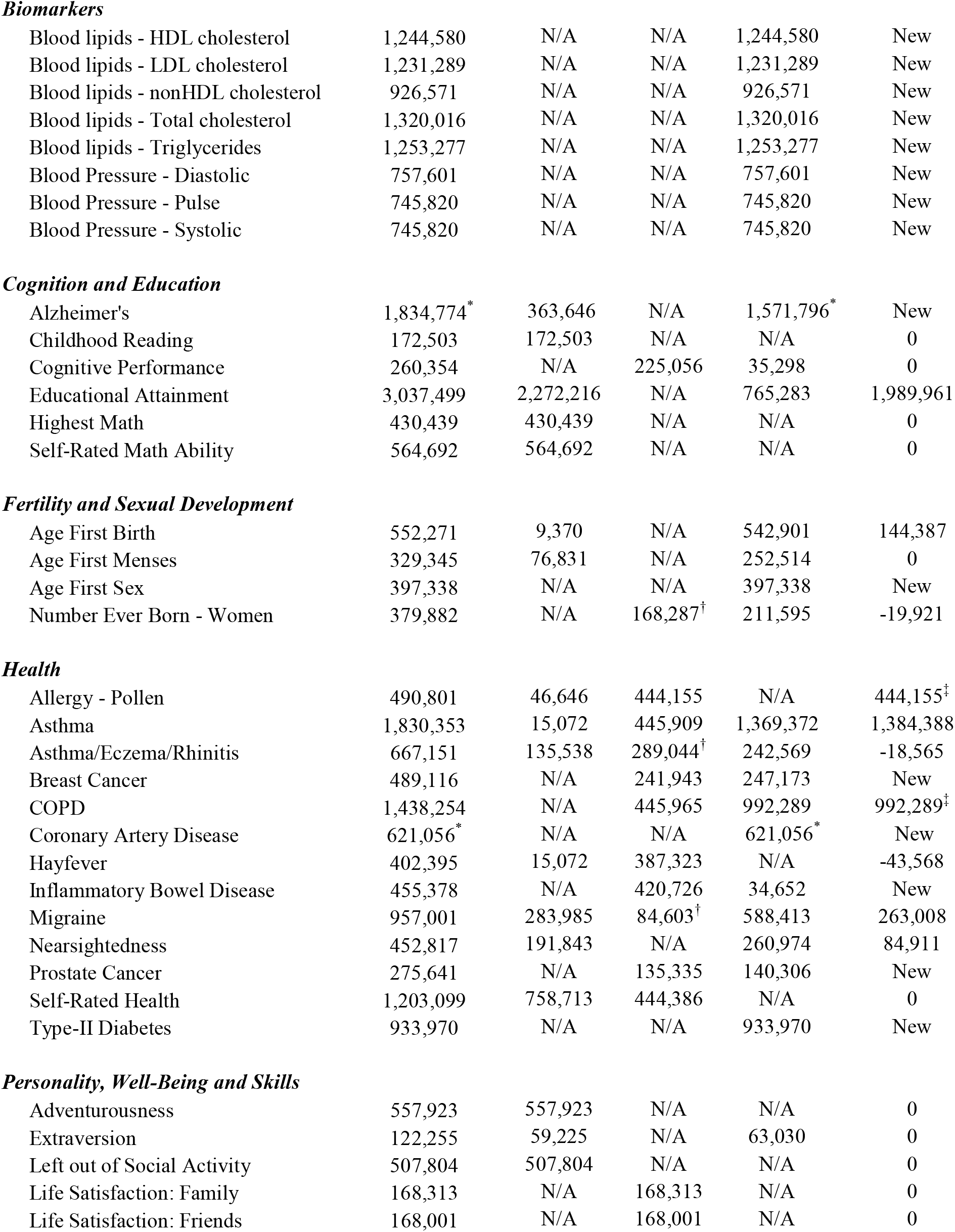
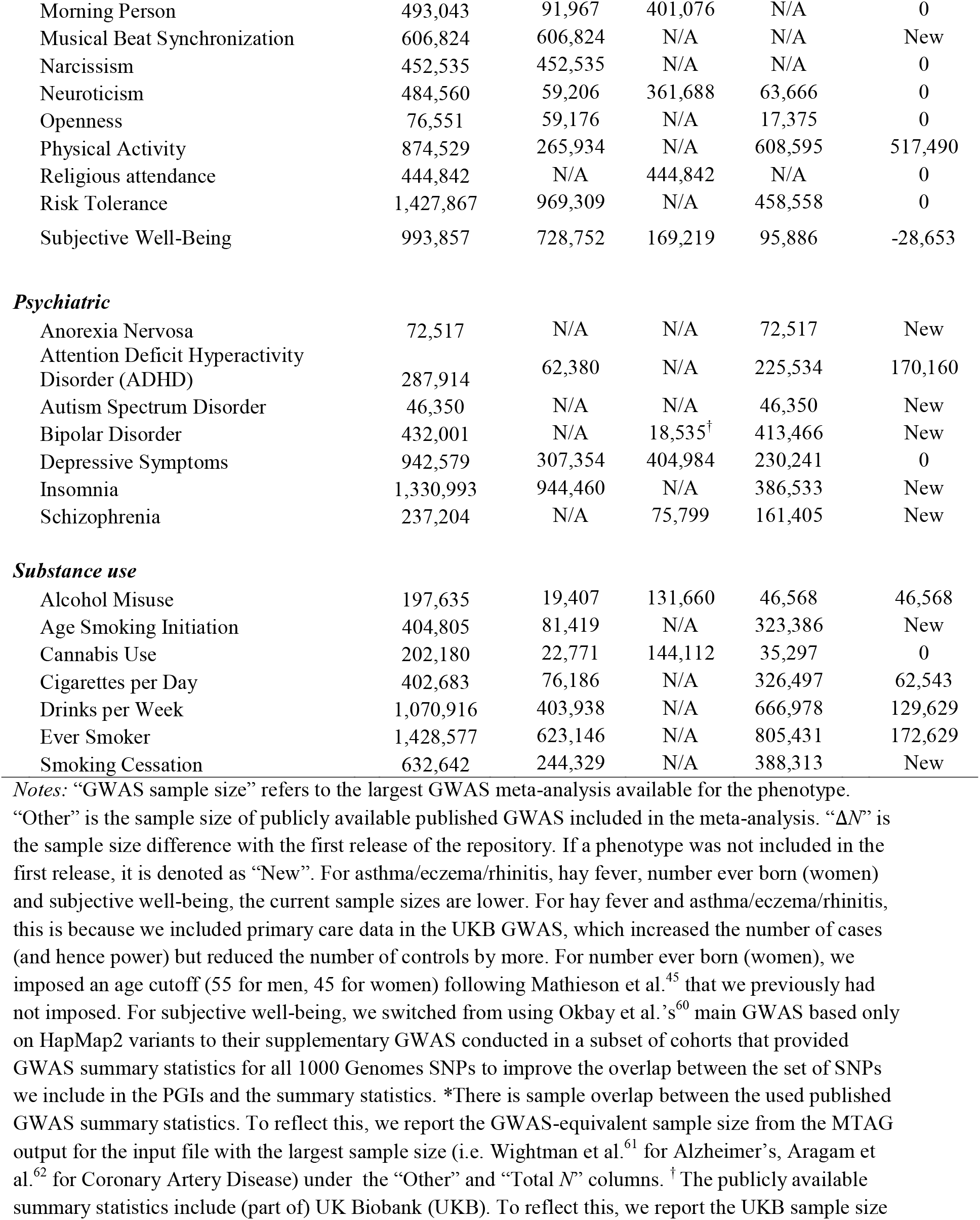
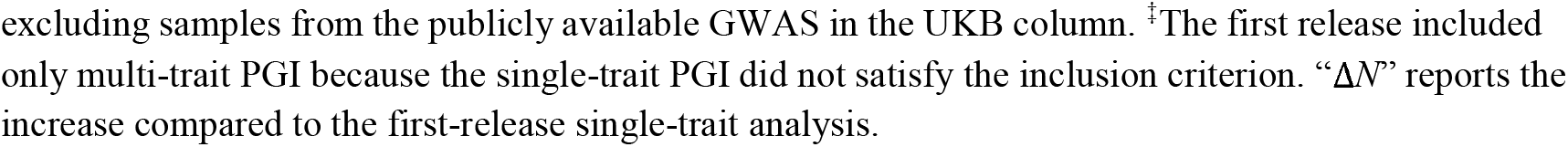
Repository phenotypes and GWAS sample sizes.

**Table 2.** Datasets participating in the repository.

| **Dataset** | **Country** | ***N* PGI** | **$\Delta N$** | ***N* with parental PGI** |
| --- | --- | --- | --- | --- |
| 1958 National Child Development Study | UK | 6,267 | New | 0 |
| 1970 British Cohort Study | UK | 5,393 | New | 0 |
| 2001 Millennium Cohort Study | UK | 17,284 | New | 6,083 |
| Avon Longitudinal Study of Parents and Children | UK | 17,450 | New | 6,624 |
| Dunedin Multidisciplinary Health and Development Study | New Zealand | 887 | 0 | 0 |
| English Longitudinal Study of Ageing | UK | 7,310 | 0 | 0 |
| Environmental Risk Longitudinal Twin Study | UK | 2,316 | 0 | 1,346 |
| Estonian Genome Center, University of Tartu | Estonia | 210,407 | 158,698 | 89,011 |
| Finnish Biobanks | Finland | 500,348 | New | 0 |
| Generation Scotland | UK | 19,966 | New | 14,135 |
| German Socioeconomic Panel (SOEP-G) | Germany | 2,495 | New | 378 |
| Health and Retirement Study | USA | 12,670 | 580 | 0 |
| Midlife in the United States | USA | 1358 | New | 0 |
| Minnesota Center for Twin and Family Research | USA | 7,634 | -20 | 2,504 |
| National Longitudinal Study of Adolescent to Adult Health | USA | 5,689 | 0 | 830 |
| Swedish Twin Registry | Sweden | 42,864 | 4,792 | 21,398 |
| Texas Twin Project | USA | 556 | 0 | 328 |
| TwinLife | Germany | 5,535 | New | 2,325 |
| UK Biobank (Full Release) | UK | 445,985 | 0 | 39,687 |
| Wisconsin Longitudinal Study | USA | 8,523 | -426 | 3,945 |

*Notes:* The “*N* PGI” column gives the number of participants in each dataset for whom the PGIs in Table 1 are supplied in the current release of the repository (i.e., those who passed the subject-level exclusion filters described in Methods). “ $\Delta N$ ” gives the increase in the number of participants in each dataset compared to the first release of the repository. If a dataset is new in the current release, it is denoted by “New”. “*N* with parental PGI” gives the number of participants for whom parental PGIs are available.

Our previous study^7^ illustrated the rise of PGI-based research in a figure that tracked the types of studies presented at the annual meeting of the Behavior Genetics Association between 2009 and 2019. At the outset, none of the studies had a PGI-based design, but by 2019, roughly 20% involved PGIs. During the same period, the prevalence of candidate-gene approaches declined sharply. There was also a modest decline in the share of studies based on twin, family, and adoption designs, although such designs remained the most common overall throughout the decade. Figure 1a updates this analysis through 2024 and shows that these trends have continued, with 40% of studies in 2024 utilizing PGI designs. Figure 1b extends the analysis by tracking the subset of PGI-based studies that employ family data (typically using sibling samples and sibling fixed-effects models). Such designs were rare prior to 2018 but have grown steadily in prominence since then. By 2024, roughly one in four PGI-based presentations used family-based designs, reflecting wider availability of genetic data on families and a growing interest in research designs that enable more credible identification of causal effects^8,9^.

**Figure 1.**
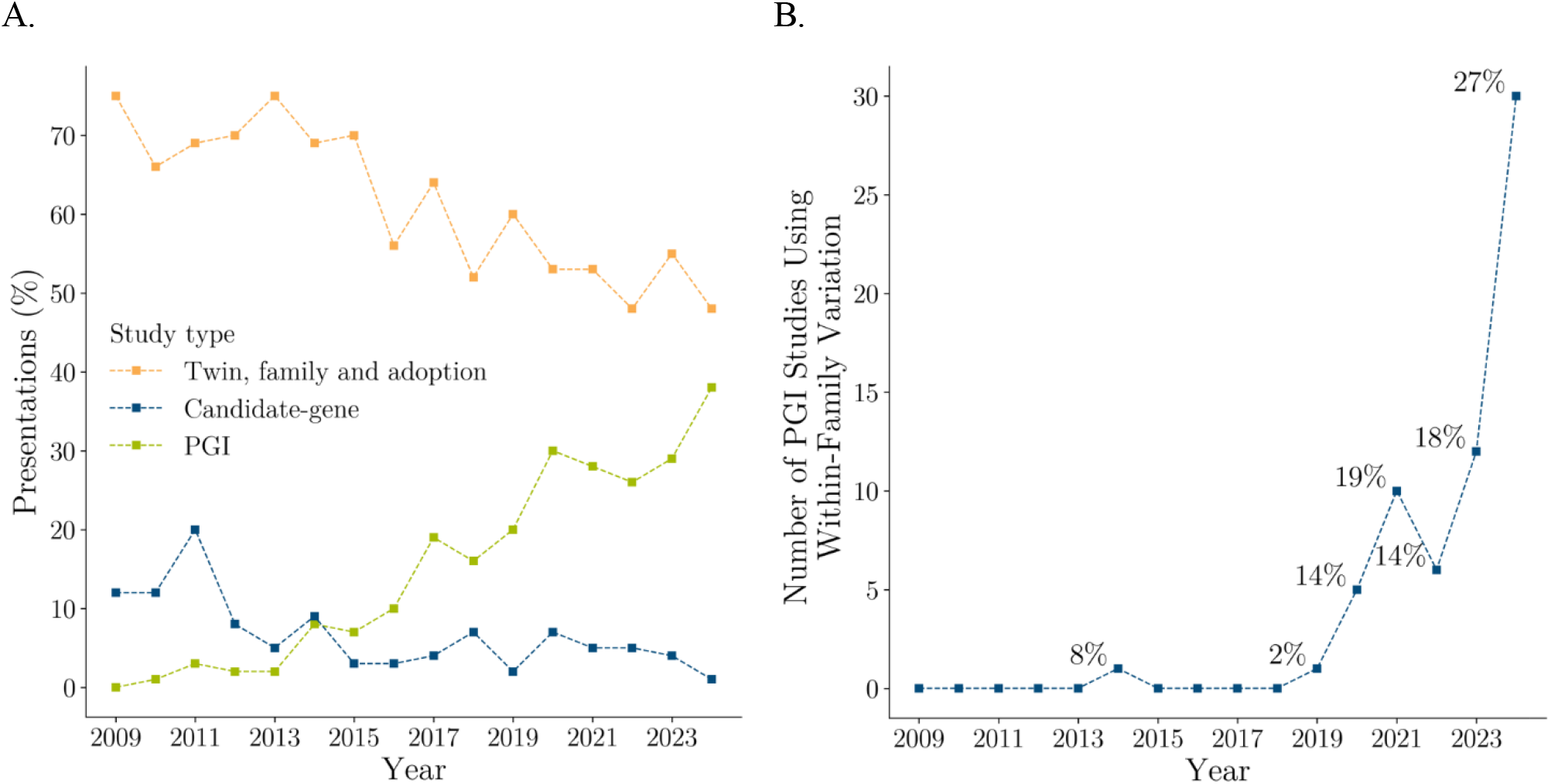
Prevalence of PGI studies in Behavior Genetics Association Annual Meeting presentations and the proportion using within-family variation. **Notes:** Panel A displays the percentages of annual Behavior Genetics Association (BGA) meeting presentations by study type. Panel B shows the number of polygenic index (PGI) studies that utilized within-family genetic variation. Using a rule-based classification algorithm, we categorized all 2,999 BGA presentations from 2009 to 2024. Each presentation was classified into one or more of four study categories (so percentages need not sum to 100%): (1) Twin, Family, and Adoption studies, (2) Candidate-Gene studies, (3) PGI (Polygenic Index) studies, and (4) PGI studies using within-family variation. An expert rater reviewed all presentations marked as “PGI studies using within-family variation” and presentations from 2023 and 2024 classified as “PGI studies,” manually recategorizing some of them. For further details and algorithm validation, see Methods.

Much of the research using PGIs in the social sciences is motivated by causal questions; for example, a researcher may want to study how the effects of genetic variants on health in middle age change when individuals stay in school for an additional year^3^. Yet, most PGI-based studies rely on observational studies of unrelated individuals. The increasing focus on family-based methods is largely driven by the realization that observational designs often fail to control for confounding factors, most notably correlation between genes and certain environmental exposures. Family-based designs help address confounding by exploiting the random segregation of alleles during meiosis, which occurs independently of environmental exposures. Some methods that were common prior to the dominance of GWAS (such as the transmission disequilibrium test, TDT) relied on this natural experiment^10,11^, but these designs have been largely abandoned because they have lower power to detect associations than standard GWAS designs.

To date, efforts to address causal research questions using family-based PGI analyses have faced several challenges. One is data availability. Even though it is widely understood that it is often desirable to control for parental PGIs in empirical analyses (an equivalent approach^12^ to that of the influential paper by Kong et al.^11^), samples with both parents genotyped are generally rare. Samples with genotyped siblings are more common, but statistical power remains a challenge. In addition, a conceptual complication arises in sibling fixed-effects analyses: because the source of variation that is leveraged is the sibling difference in PGI values, the estimator picks up any effect of a sibling’s genes on the person’s own phenotype (what are often called “sibling indirect genetic effects”)^13^. Therefore, the sibling fixed-effects estimator may be biased when, as is typical, a researcher is interested in isolating the causal effect of a person’s genes on their own phenotype.

The second category of updates to the Repository is motivated by the growing interest in family-based methods and seeks to address the associated challenges. We use Mendelian imputation^12^ to impute missing parental PGIs in the 11 participating datasets that include first-degree relatives, and we include these imputed parental PGIs in the new data release. By doing so, we increase the number of datasets in which it is feasible to control for parental PGIs in analyses. Importantly, the estimator for the coefficient on an individual’s PGI remains unbiased and consistent (for isolating the effect of a person’s genes on their own phenotype) even when parental PGIs are constructed from imputed rather than directly observed parental genotypes^12^. Moreover, because the imputation leverages the nonlinear laws of Mendelian inheritance, imputing and controlling for parental PGIs adds information and therefore is a more statistically powerful strategy than using sibling fixed effects when applied to sibling-pair data. It also enables the analysis to include additional data types such as genotyped parent-offspring pairs^12^.

We illustrate the utility of the newly available parental PGIs in two applications. First, in two Repository cohorts, UKB and WLS, we compare PGI-phenotype associations with and without parental PGI controls for all available phenotypes. Second, we revisit a recent study by Mostafavi et al.^14^, which examined how the predictive power of PGIs varies across subgroups *within* a genetic ancestry group in UKB. We focus on two specific cases—diastolic blood pressure by sex and BMI by age group—and, using parental PGIs, investigate one of the future directions proposed by the authors: disentangling the causal and non-causal components of PGI associations to better understand their portability across subgroups.

We also develop a theoretical framework that facilitates interpretation of PGI studies with controls for parental PGIs. We consider the case where the observed PGI is constructed using noisy SNP weights from a finite-sample population-based GWAS, while the true variable of interest is a latent PGI based on the (unobserved) ideal weights from a family-based GWAS conducted in an infinite sample. When researchers use the observed PGI in empirical analysis, there are two sources of error. The first is that the ideal SNP weights—the true parameters from a family-based GWAS—are not observed and must therefore be estimated, creating the familiar errors-in-variables problem analyzed in our previous paper and other work^15–17^. The second is the source of error analyzed by Trejo and Domingue^18^: even in the limit of an infinite sample, the true parameters from the standard GWAS will generally be different from those of the family-based GWAS. We find that both sources of error attenuate the coefficient on the PGI, and we provide an analytic formula that can be used to correct both the coefficient on the main effect of the PGI and the coefficients on interactions between the PGI and environmental factors. For an extended, and less technical, discussion of how the results of a family-based PGI study should be interpreted, see the Frequently Asked Questions (FAQs in Supplementary Information).

In what follows, we first describe the expanded Repository and assess the predictive power of the updated PGIs. We then report the two illustrative applications of the imputed parental PGIs. We conclude by presenting the theoretical framework for interpreting estimates from PGI studies that control for parental PGIs.

## Results

### Candidate Phenotypes

We considered a list of 26 new phenotypes for potential inclusion in the updated release of the Repository: eight biomarkers, five fertility or sexual development outcomes, five mental disorders, five other diseases, one skill, and two substance-use outcomes. As in the first release of the Repository, we combined GWAS summary statistics from all available source—previously published studies that made their data public, analyses in research-consented individuals from consumer genetics and research company 23andMe, Inc., and analyses in UKB—and used the meta-analyzed summary statistics to calculate the theoretically expected predictive power of each candidate PGI (see Methods, Supplementary Table 1). We found that 22 phenotypes had an expected predictive power above our chosen inclusion threshold of 1% and proceeded to include them in the current release.

### Updated GWAS Meta-Analysis

The first release of the Repository contained PGIs for 47 phenotypes that met the expected predictive power criterion for “single-trait” (36 phenotypes) and/or “multi-trait” PGIs (35 phenotypes). “Single-trait” PGIs were based on meta-analyses of GWAS conducted on the same phenotype in different samples, whereas “multi-trait” PGIs were based on multi-trait analyses of genetically correlated phenotypes^19^. For simplicity, we do not include multi-trait analyses in the current release and limit the list of PGIs to those based on single-trait analyses that satisfy our inclusion criterion. The 47 phenotypes in the first release were selected from an original list of 53 candidate phenotypes (separate from the 26 new candidates discussed above). For all 53 phenotypes, we systematically looked for new data that could be used to increase the combined sample size of the GWAS meta-analysis. For 14 phenotypes, we found a larger published GWAS that was used to update the meta-analysis. For 11 phenotypes, we also reran the GWAS in UKB using either an updated phenotype definition or new phenotypic data that were not available at the time of the first release (Supplementary Table 2). As a result, the expected predictive power improved for several phenotypes, tipping two phenotypes with no single-trait PGIs in the first release over the 1% cutoff. For one phenotype with updated GWAS summary statistics, Alzheimer’s disease, we waived the expected predictive power criterion because the expected predictive power is calculated using the heritability estimate from LD score regression, which is based on HapMap3 variants that do not tag the APOE region well, leading to a downward bias.

For each of the 61 phenotypes included in the second release of the Repository, Table 1 lists the total sample size of the GWAS meta-analysis, the sample size contributions from the three sources, and the increase in sample size for phenotypes included in the first release. The sample size increased for 14 phenotypes from the first release (see Table 1 caption for explanations of the small decrease in sample size reported for four of the phenotypes).

### SNP Weights

We updated our methodology for constructing SNP weights from LDpred^20^ to SBayesR^21^, and included the set of ~2.9 million SNPs from Lloyd-Jones et al.^21^ in the PGIs instead of ~1.2 million HapMap3 SNPs used in the prior release. Therefore, all PGIs in the Repository have been updated, including phenotypes whose GWAS sample sizes stayed the same.

### Parental Genotype Imputation

Thirteen of the Repository cohorts are family-based samples with data on twins, non-twin siblings, parent-offspring pairs, and/or trios (Table 2). To facilitate maximally powered family-based PGI analyses in the participating family-based cohorts (and eliminate the need for sibling-fixed-effects analyses), we include, in the current release of the Repository, PGIs for the parents of the genotyped individuals with genotyped siblings and/or at least one genotyped parent. We impute the genotypes of missing parent(s) based on Mendelian laws using the software tool *snipar*^12^ (Methods). Previous work has shown that regressions controlling for parental PGIs imputed in this way generate unbiased and consistent estimates of the causal effects of the PGIs (as defined below) and that, relative to alternative approaches, this approach more efficiently uses the genotype data of family members^12^.

### Predictive Power of PGIs

To showcase the predictive power of our PGIs, we ran analyses in three repository cohorts: the Health and Retirement Study, a representative sample of Americans over the age of 50; the Wisconsin Longitudinal Study, a sample of individuals who graduated from high school in Wisconsin in 1957; and the third partition of the UKB. (As in the first release, we split UKB into three equal-sized partitions so that we can include two of the partitions in the GWAS when making PGIs for the remaining partition (Methods).) Our measure of predictive power is incremental *R*^2^, calculated as the increase in the coefficient of determination (*R*^2^) when the PGI is added to a regression of the phenotype on covariates (Methods). The analyses shown here are from specifications that do not include controls for imputed parental PGIs. Figure 2 shows the observed incremental *R*^2^ of the PGIs in the three cohorts. Supplementary Table 3 lists the incremental *R*^2^’s alongside the results from the first release where available to demonstrate the improvement. The largest absolute increase in predictive power is observed for height (~10 percentage points), for which we also see the greatest increase in the input GWAS sample size, followed by educational attainment (~3 percentage points). For several phenotypes for which we updated the UKB phenotype definition by incorporating primary care data, the predictive power increased substantially (e.g., 253% increase for asthma/eczema/rhinitis in the third partition of UKB). For some other phenotypes with no change in the input GWAS, the predictive power improved up to 47% (self-rated health) because of the change in methodology. We also observe reductions in predictive power for some phenotypes, but these are either statistically indistinguishable from zero and/or inconsistent across the validation cohorts.

**Table 3:** Proportional Bias of PGI Effect Estimates in a Causal PGI Study.

| Phenotype | Repository<br>Effective<br>GWAS $N$ | $h_{SNP}^2$ | $R^2$ | $\rho = \sqrt{\frac{h_{SNP}^2}{R^2}}$ | $\text{Corr}(\mathbf{x}_i\boldsymbol{\gamma}, \mathbf{x}_i\boldsymbol{\mu})$ | $\frac{\text{Corr}(\mathbf{x}_i\boldsymbol{\gamma}, \mathbf{x}_i\boldsymbol{\mu})}{\rho}$ |
| --- | --- | --- | --- | --- | --- | --- |
| Educational Attainment | 2,155,939 | 0.133<br>(0.003) | 0.110 | 1.100<br>(0.012) | 0.859<br>(0.028) | 0.781<br>(0.027) |
| Ever-smoker | 1,150,055 | 0.085<br>(0.006) | 0.053 | 1.266<br>(0.045) | 0.974<br>(0.019) | 0.769<br>(0.031) |
| Height | 3,839,667 | 0.231<br>(0.011) | 0.216 | 1.034<br>(0.025) | 0.970<br>(0.005) | 0.938<br>(0.023) |
| BMI | 582,457 | 0.247<br>(0.009) | 0.175 | 1.188<br>(0.022) | 1.000<br>(0.008) | 0.842<br>(0.017) |
| Age at first birth (women) | 181,585 | 0.169<br>(0.007) | 0.057 | 1.722<br>(0.036) | 0.881<br>(0.067) | 0.512<br>(0.040) |
| Cognitive performance | 222,914 | 0.232<br>(0.008) | 0.107 | 1.472<br>(0.025) | 0.975<br>(0.045) | 0.662<br>(0.033) |
| Non-HDL cholesterol | 926,571 | 0.102<br>(0.018) | 0.062 | 1.283<br>(0.113) | 0.959<br>(0.030) | 0.747<br>(0.070) |
| Asthma | 353,336 | 0.135<br>(0.011) | 0.060 | 1.500<br>(0.061) | 0.963<br>(0.035) | 0.642<br>(0.035) |

*Notes:* Standard errors are reported in parentheses. The Repository effective GWAS sample size is pulled from the GWAS-equivalent $N$ column of Supplementary Table 1. Estimate of $h_{SNP}^2$ , $\text{Corr}(\mathbf{x}_i\boldsymbol{\gamma}, \mathbf{x}_i\boldsymbol{\mu})$ , and their standard errors are from Tan et al.’s<sup>33</sup> Supplementary Table 3. We note that the reported $\text{Corr}(\mathbf{x}_i\boldsymbol{\gamma}, \mathbf{x}_i\boldsymbol{\mu})$ estimates are obtained using LDSC and are likely to be overestimates because LDSC removes much of the contribution from population stratification to the correlation (and population stratification lowers the true $\text{Corr}(\mathbf{x}_i\boldsymbol{\gamma}, \mathbf{x}_i\boldsymbol{\mu})$ ). $R^2$ is calculated using the expected $R^2$ formula from Daetwyler et al.<sup>63</sup> using the numbers from the Repository effective GWAS sample size and $h_{SNP}^2$ columns and an effective number of SNPs set equal to 60,000. Estimates of $\rho = \sqrt{h_{SNP}^2/R^2}$ and $\text{Corr}(\mathbf{x}_i\boldsymbol{\gamma}, \mathbf{x}_i\boldsymbol{\mu})/\rho$ are calculated by plugging in estimates from earlier columns, with standard errors computed using the delta method in both cases (assuming the estimates of $\text{Corr}(\mathbf{x}_i\boldsymbol{\gamma}, \mathbf{x}_i\boldsymbol{\mu})$ and $\rho$ are independent, even though that it is unlikely to be true).

**Figure 2.**
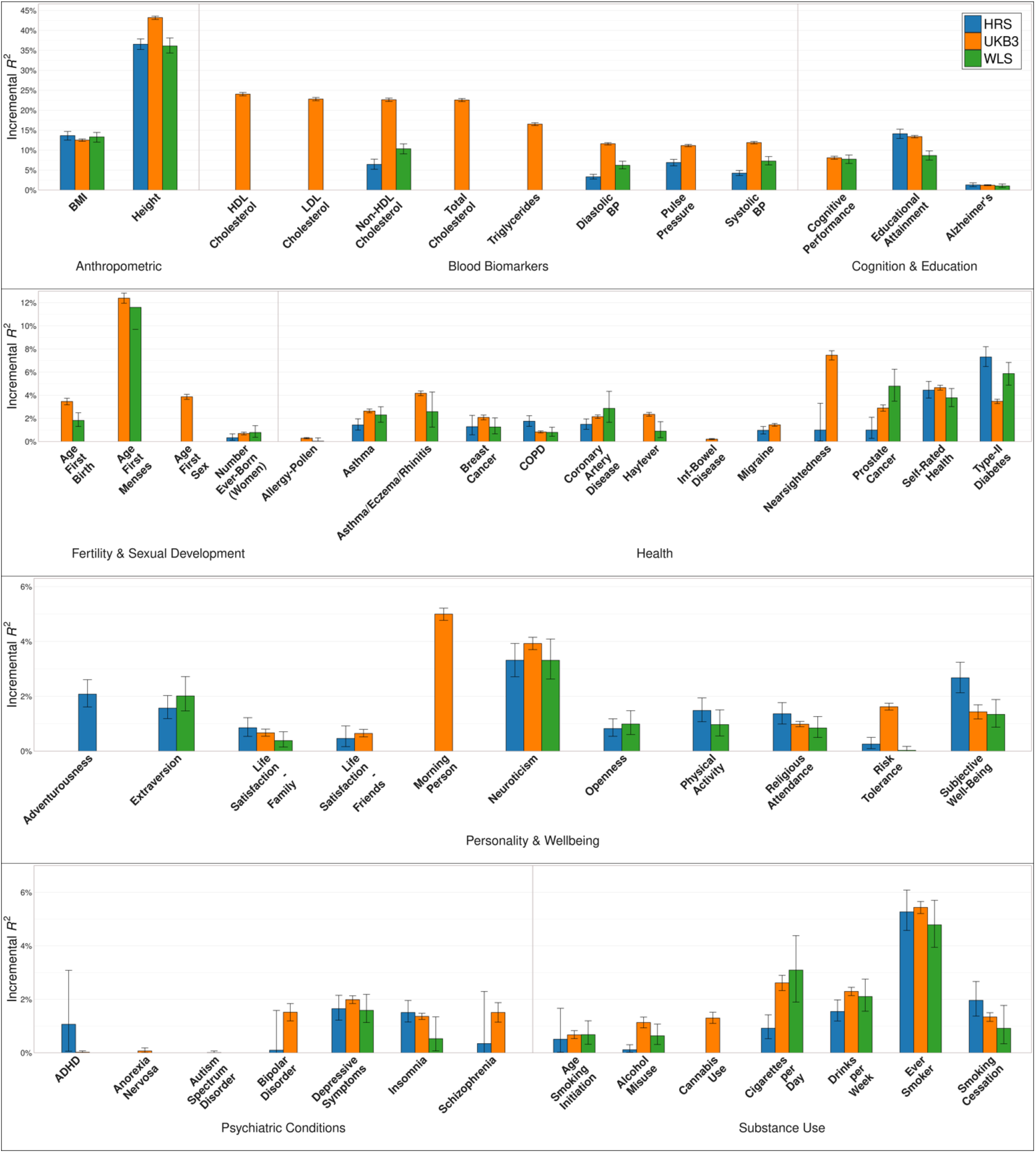
Predictive power of repository PGIs in three validation datasets. **Notes:** Incremental *R*^2^ from adding the PGI to a regression of the phenotype on 10 principal components of the genetic relatedness matrix for HRS and WLS, and on 20 principal components and 106 genotyping batch dummies for UKB (third partition). Prior to the regression, all phenotypes are residualized on a second-degree polynomial in age or birth year, sex, and their interactions (see Supplementary Tables 2 and 14). Error bars are 95% confidence intervals from bootstrapping with 1,000 repetitions. For the sample sizes of the GWAS that the PGIs are based on, see Supplementary Table 11.

### Illustrative Applications of Parental PGIs

In an observational study of unrelated individuals—the study design of most applications of PGIs to date—the association between the PGI and the phenotype may be confounded due to gene-environment correlation (including bias from population stratification), causal effects of parental (and sibling) genotypes on the environment faced by an individual (often called “indirect effects”), and assortative mating^22^. Controlling for parental PGIs is a powerful strategy for addressing concerns about these sources of confounding and strengthening the causal interpretability of estimates. Here, we provide two empirical illustrations using newly released PGIs in the Repository.

In the first and more straightforward analysis, we use two Repository cohorts with parental PGIs—UKB and WLS—to compare parameters that we term the population association (*ψ*) and the causal effect (*δ*, often referred to as the “direct effect”). The population association is defined as the coefficient on the individual’s PGI in a regression of the phenotype on that PGI, *without* controls for parental PGIs. The causal effect is defined as the coefficient on the individual’s PGI from an otherwise identical regression that also includes the parents’ PGIs as controls (Methods). We use the term “causal effect” as a convenient shorthand, without implying that *δ* is invariant across individuals or contexts. Below, we discuss additional caveats regarding the appropriate interpretation of this parameter^23–25^.

Supplementary Table 4 reports the estimates and confidence intervals of *ψ, δ*, and the ratio *δ*/*ψ* in UKB and WLS, for all phenotypes with available data (Supplementary Figures 2-5). It also includes the inverse-variance-weighted meta-analysis of the estimates. Of the 54 phenotypes analyzed, 33 are available in both cohorts, while the remaining 21 are available in only one; for the latter, the meta-analysis estimate is thus identical to the single-cohort estimate. Figure 3 depicts the meta-analyzed estimates of *ψ* and *δ* (for the meta-analyzed ratios, see Supplementary Figure 1).

**Figure 3.**
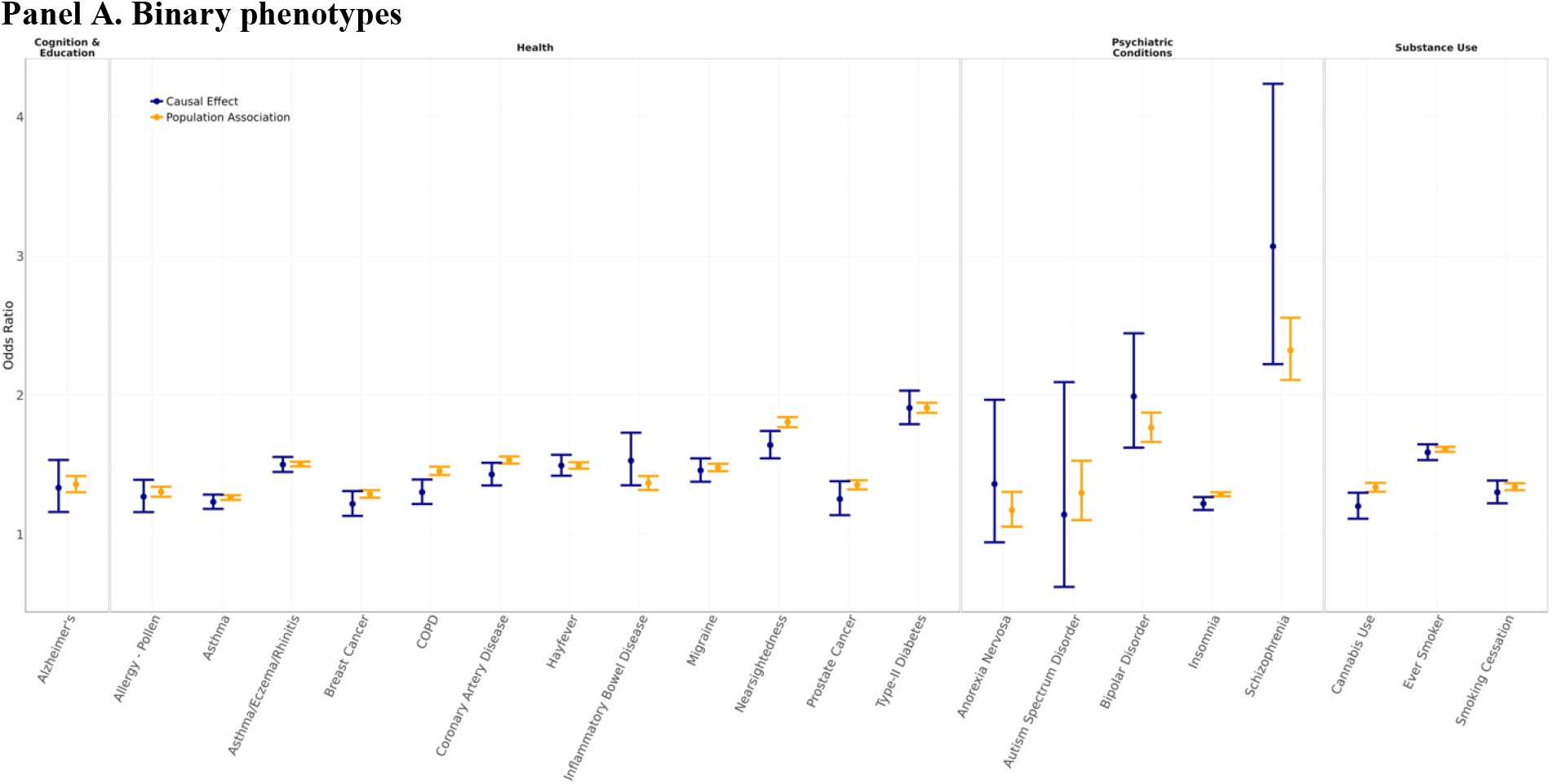
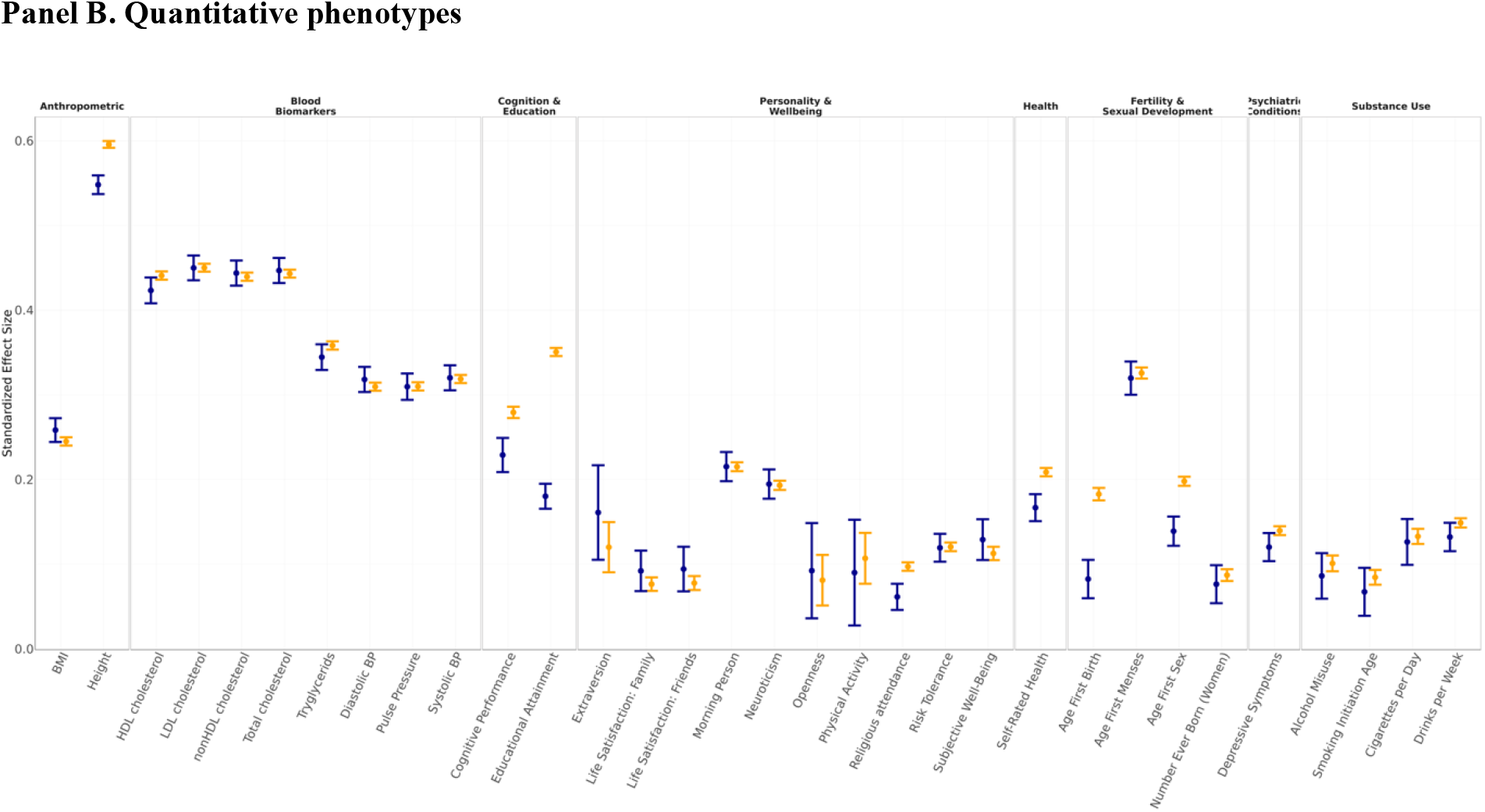
Causal effects of PGIs versus their population associations. **Notes:** Causal effects and population associations of PGIs from an inverse-variance weighted meta-analysis of UKB and WLS. Causal effects were estimated in the sample of first-degree relatives, and population associations in samples of unrelated individuals (third partition in UKB, subsample of individuals with no first-degree relatives in WLS). Panel A: Population associations were estimated using logistic regression; causal effects were estimated using a generalized linear mixed model with logistic link function, including a random intercept for family ID. Panel B: Population associations were estimated using linear regression; causal effects were estimated using from a linear mixed model, including a random intercept for family ID. Error bars are 95% confidence intervals. Full details of the models are provided in Methods.

Because the sampling distribution of the ratio *δ* / *ψ* is not well-behaved for *ψ* close to 0, we conduct hypothesis tests using the null hypothesis that the difference *δ* - *ψ* equals zero. However, because the *δ* - *ψ* difference lacks intuitive interpretation (particularly for binary phenotypes, where it is a difference in log-odds), we instead report *δ* / *ψ* estimates for phenotypes where 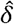 and 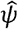 are statistically distinguishable. At a Benjamini-Hochberg^26^ 5% false discovery rate, our meta-analysis identified 11 phenotypes with a difference statistically distinguishable from zero. In all cases, the causal effect is estimated to be smaller than the population association. It is notable that *none* of the 11 phenotypes are biomarkers. We estimate substantial differences between the meta-analyzed causal effect and population association for several phenotypes related to cognition, fertility and substance use: 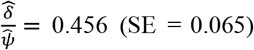 for age at first birth, 0.513 (0.022) for educational attainment, 0.634 (0.138) for cannabis use, 0.702 (0.046) for age at first sex, 0.714 (0.096) for COPD, the leading cause of which is cigarette smoking^27^, and 0.821 (0.038) for cognitive performance. We also reject a difference of zero for religious attendance: 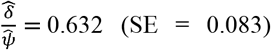,insomnia: 0.780 (0.079), self-rated health: 0.801 (0.041), nearsightedness: 0.839 (0.054), and height: 0.921 (0.010). Overall, we find systematically stronger evidence for PGIs for behavioral and social phenotypes—those for which genes are likely to exert their influence through complex, environmentally mediated pathways—to have causal effects that are smaller than their population associations. A possible exception to this pattern is height, but the estimated ratio is close to one, and the small standard error reflects the exceptionally high predictive accuracy of the height PGI.

Our second analysis revisits a recent study by Mostafavi et al.^14^ that examined the predictive accuracy of PGIs across subgroups *within* a genetic ancestry group: the “White British” UKB sample, as defined by UKB based on self-reported ethnicity and genetic principal components^28^. The original paper focused on three case studies: predicting diastolic blood pressure by sex, BMI by age, and educational attainment across adult socioeconomic status (SES). The paper found variation in predictive accuracy across all three and explored several possible explanations. The paper concludes by proposing to study PGIs in a family-based design in order to make further progress distinguishing among explanations, in particular to shed light on the portability across subgroups of causal versus non-causal components of PGI associations. Here, we take some first steps in this direction.

Our approach differs from that of the original paper in three main ways. First, whereas Mostafavi et al. used (non-overlapping) samples from UKB for both training and prediction, we leverage the considerably more powerful Repository PGIs in our analyses. Second, we focus on two of the three case studies, omitting the analysis of heterogeneity in the predictive power of educational attainment by adult SES because adult SES is clearly endogenous to the PGI for educational attainment. Third, the original paper focused on comparing the incremental *R*^2^ of the PGI across subgroups, whereas we emphasize comparisons of standardized regression coefficients, relegating incremental *R*^2^ results to the Supplementary Information (see Supplementary Table 5 and Supplementary Figure 6). We focus on the standardized coefficients because: (a) they are more interpretable in regressions that control for parental PGIs; (b) they are not confounded by differences in environmental variance across subgroups; and (c) Mostafavi et al. found that the observed differences in prediction accuracy could be explained by a model in which SNP effect sizes are uniformly scaled up in the higher-accuracy group.

Figure 4 depicts the results of our analysis (see also Supplementary Table 5). We replicate Mostafavi et al.’s finding that the PGI for diastolic blood pressure is more predictive in women than men (Panel A). When we control for parental PGIs, the point estimates for both women and men remain virtually unchanged but are less precise. The female/male ratio of population associations is 1.135 (95% CI = [1.065,1.205]), compared to a female/male ratio of 1.122 for the causal effects (95% CI = [1.004, 1.240]). (Unlike in our first illustrative application, the point estimates are never close to zero and we can bootstrap confidence intervals, so we conduct all analyses on ratios.) Directionally, we also replicate the finding that the predictive power of a PGI for BMI is greater among young subjects, although the young/old ratio of population associations is not statistically distinguishable from one: 1.028 (95% CI = [0.942, 1.115]) (Panel B). The estimated young/old ratio of causal effects is similar: 1.053 (95% CI = [0.911, 1.195]). In both case studies, our finding that controlling for parental PGIs does not have a statistically detectable effect on subgroup differences helps cast doubt on several possible explanations that Mostafavi et al. could not address, including that the differences in PGI predictive power across subgroups are driven by gene-environment correlation, effects of parental (or sibling) genotypes that differ across subgroups, and, for the age results, changes in assortative mating over time.

**Figure 4.**
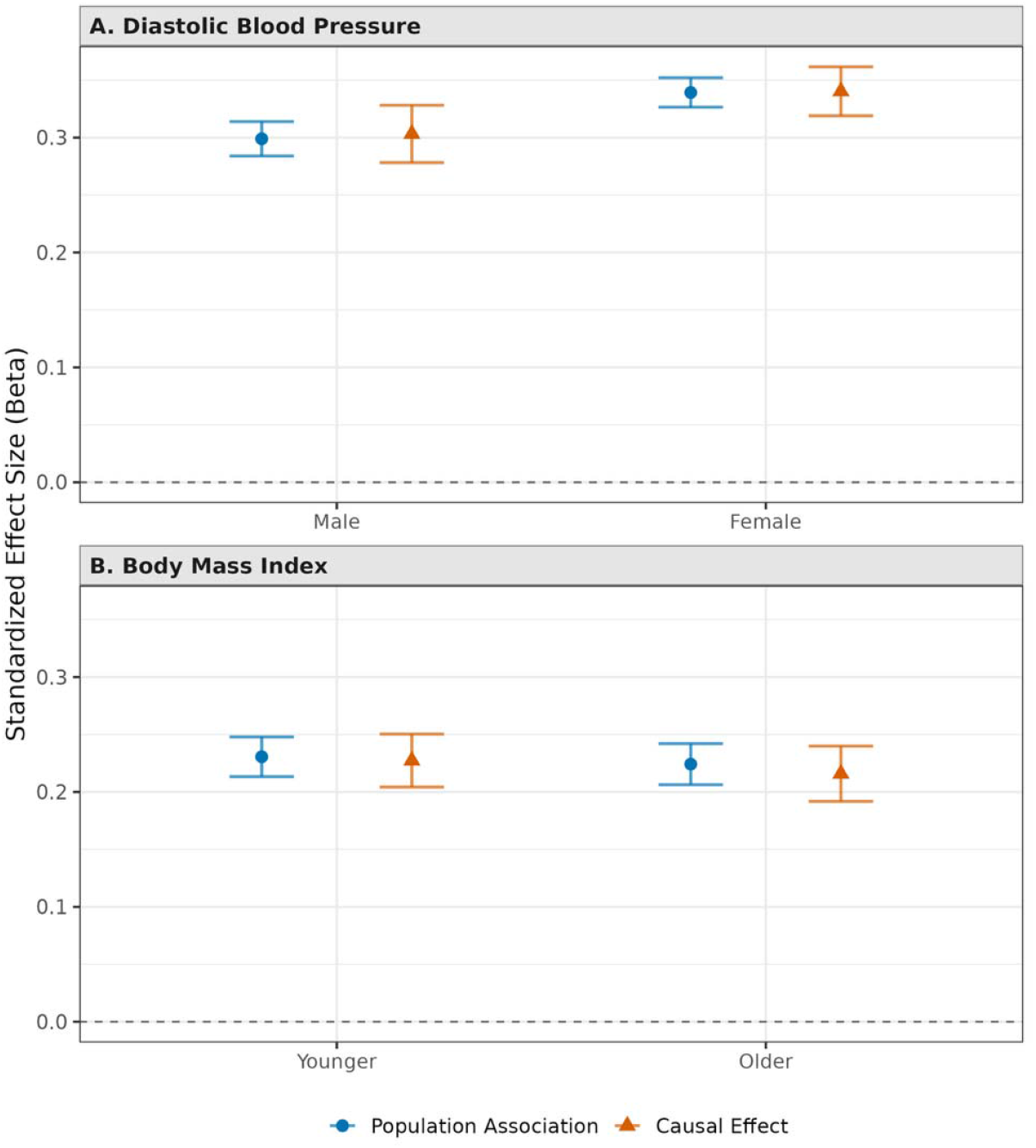
Causal effects and population associations of PGIs by demographic group. **Notes:** Standardized regression coefficients of PGIs for diastolic blood pressure and BMI from models that do (triangle) and do not (circle) control for parental PGIs. All analyses were conducted in the first-degree-relatives subsample of the UKB. Panel A: PGI effect on diastolic blood pressure by sex. Panel B: PGI effect on body mass index by age group (below vs. above median age). Error bars are 95% confidence intervals. Full details of the models are provided in Methods.

### Theoretical Framework

We develop an analytic framework to facilitate the interpretation of PGI analyses that control for parental PGIs. We begin by establishing some notation. Let 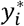 denote individual *i’*s phenotype value, 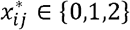 denote *i*’s allele count at SNP *j* and 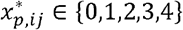 denote the sum of parental allele counts (sum of paternal and maternal counts) at SNP *j*. We define mean-centered variables as:

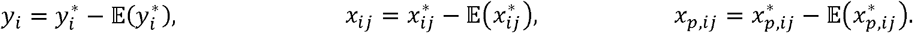

Finally, let ***x***_*i*_ =(*x*_*i*1_, *x*_*i*2,…,_ *x*_*iJ*_) and ***x***_*i*_ =(*x*_*p,i*1_, *x*_*p,i*2,…,_ *x*_*p,iJ*_) denote the length-*J* vectors of mean-centered allele counts at the SNPs included in individual *i*’s PGI and the PGI of *i*’s parents, respectively. For simplicity, here we assume that the causal effects of the genetic variants are homogeneous across individuals.

In a more realistic scenario where causal effects vary across individuals, the causal parameters below should be interpreted as weighted averages. Individuals with heterozygous parents receive more weight because the variance in their genotype is greater (see refs. ^23^ and ^25^). In this model, “causal effect of a genetic variant” is a convenient shorthand for such a weighted average.

Due to meiosis, ***x***_*i*_ is randomly assigned conditional on the maternal and paternal alleles^29^. The coefficient vector 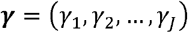 from the population regression

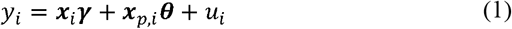

is therefore free from confounding due to gene-environment correlation and has a causal interpretation (Supplementary Note). Specifically, ***x***_***i***_ ***γ*** can be interpreted as the best linear approximation to the causal genetic component, given the set of SNPs included in the analysis^25^. It captures causal effects of SNPs included in ***x***_***i***_ and also of genetic variants not included in ***x***_*i*_ to the extent that those genetic variants are correlated with (i.e., “tag”) the SNPs in ***x***_*i*_ due to linkage (i.e., physical proximity). Note that ***γ*** would remain unchanged if controls were included in Equation (1), as long as those controls are causally prior to ***x***_*i*_ (i.e., the controls are not themselves causally affected by ***x***_*i*_). The coefficient on the parental genotype vector, ***θ***, captures (a linear approximation to) causal effects of parental genotypes on *y*_*i*_ that operate through parental phenotypes that affect individual *i*’s environment—which we call *parental genetic effects* (the more common term is “parental indirect genetic effects”)—but it also captures confounding from gene-environment correlation, population stratification, and effects of genetic variants not included in ***x***_***i***_ but that are correlated with ***x***_***i***_ due to non-random mating (assortative mating and population structure)^30^. Adding controls could generally change the coefficients ***θ***, in even when ***γ*** remains unchanged.

Define the *causal additive SNP factor* as 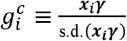,The corresponding *parental additive SNP factor* is 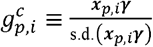.We refer to the variance explained by the causal additive SNP factor as the *causal SNP heritability*. (If ***x***_*i*_ and ***x***_*p,i*_, contained all genetic variants, then 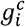 would be the *causal additive genetic factor*, and the phenotypic variance explained by ***x***_***i***_ ***γ*** would be the *narrow-sense heritability*.)

We denote individual *i*’s own PGI as

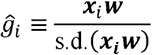

and the parental PGI of individual *i* as

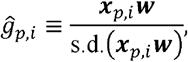

where ***w*** is some nonzero vector of SNP weights and s.d. (·) is the standard deviation. Note that when ***w = γ***, the PGIs correspond to the causal and parental additive SNP factors defined above, respectively.

In many (but not all) empirical applications of PGIs, the parameters of interest are defined by an ideal population regression that takes the general form:

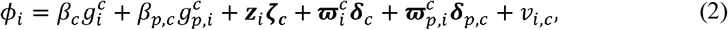

where *ϕ*_*i*_ is some de-meaned phenotype, which need not coincide with the phenotype that the PGIs were trained to predict; is a vector of controls that are causally prior to the genotype vector ***z***_*i*_; is a vector is a vector of controls that are causally prior to the genotype vector 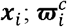 is a vector of interactions between 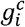 and a subset of the elements of ***z***_*i*_ (possibly all of them), denoted ***z***_*int, i*,_; and 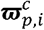 is a vector of interactions between and 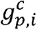, and ***z***_*int, i*,_. We assume in what follows that the interacted controls ***z***_*int, i*_, are independent of ***x***_*i*_ and ***x***_*p,i*_, as would be the case for a causal gene-environment interaction study, where the interacted controls are randomly assigned experimentally or quasi-experimentally. From these assumptions and the random transmission of alleles from parents to offspring, it follows that *β*_*c*_ and ***δ***_*c*_ have a causal interpretation (though, *β*_*p,c*_, ***δ***_*p,c*_, and ***ζ***_*c*_ in general do not; for example, the coefficients on the parental PGI variables may be confounded by gene-environment correlation, including population stratification, and assortative mating^30,31^).

In practice, it is not possible to estimate the parameters of Equation (2) directly because ***γ*** is unknown and thus 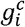 and 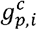 are unavailable. Although ***γ*** can be estimated from the summary statistics of a family-based GWAS^32,33^, the estimates are noisy, mostly because sample sizes are relatively small. Indeed, even though standard population-based GWAS is a biased estimator of ***γ***, at present, the mean squared error of the estimates from standard GWAS tend to be far smaller than the unbiased estimates from family-based GWAS due to the smaller estimation error^8^. Consequently, PGIs derived from population-based GWAS have much greater predictive power. For that reason, most PGIs used in practice—including those in the Repository—have been constructed from standard (non-family-based) GWAS.

To understand these PGIs constructed from standard GWAS, consider the population regression of *y*_*i*_ on ***x***_*i*_ :

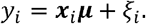

This is the same population regression as Equation (1) but without controlling for the parental genotypes. The resulting coefficient is ***μ* = *γ* + *θ***, with the ***θ*** component capturing confounding, including gene-environment correlation (see Trejo and Domingue’s equation (4a.v) and Supplementary Methods). More generally, if Equation (1) substitutes parental genotypes with imperfect controls (i.e., controls that mitigate but do not eliminate confounding), then the difference between ***μ*** and ***γ*** depends on how effectively the controls mitigate confounding. If confounding remains, then ***μ*** does not have a causal interpretation. Standard methods for constructing SNP weights from GWAS summary statistics, such as SBayesR^21^ (which we use) and LDpred^20,34^, generate a SNP-weight vector 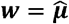 that is a consistent estimate of ***μ***. We assume that 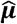 is an unbiased estimate of ***μ***, which we showed in our earlier paper^7^ is a good approximation when the GWAS sample size is large. Below, we refer to ***x***_*i*_***μ*** as the *associational additive SNP factor*, and we refer to the proportion of phenotypic variance explained by it as the *associational SNP heritability*, denoted 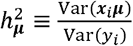. This parameter can be estimated using genomic-relatedness-matrix restricted maximum likelihood (GREML)^35^ in a sample of unrelated individuals or using LD score regression^36^, but these estimators can be biased under assortative mating.

In a family-based PGI study, since researchers cannot estimate the ideal regression Equation (1), they instead estimate the feasible regression:

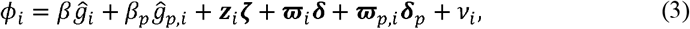

which differs from Equation (1) by replacing 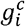 by 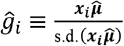 and 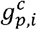 by 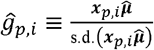.

How do the estimands *β* and ***δ*** from the feasible regression Equation (3) relate to the estimands *β*_*c*_ and ***δ***_*c*_ from the ideal regression Equation (1)? In the Supplementary Methods, we show that under random mating, the proportional bias in *β* and in every element *δ*_*k*_ of the vector ***δ*** is:

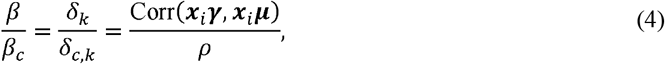

where 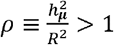 and *R*^2^ is the fraction of variance explained in a regression of *y*_*i*_ on 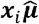. We also show that the standard error of the Ordinary Least Squares estimate of *β* (obtained by estimating Equation (3)) is attenuated relative to the standard error of the Ordinary Least Squares estimate of *β* _*c*_ (obtained by estimating Equation (1)) by the same proportional bias, Corr (***x***_*i*_ ***γ, x***_*i*_***μ***) / *ρ*. (See Supplementary Methods for simulation evidence that suggests that theoretical predictions based on Equation (4) remain accurate under realistic amounts of assortative mating, with the true attenuation slightly greater than the theoretical prediction derived under a random-mating benchmark.)

To understand this proportional bias, we discuss the numerator and denominators separately. Intuitively, *ρ* captures attenuation from sampling error in the underlying GWAS. The sampling variation creates a standard errors-in-variables bias that attenuates the coefficients, as discussed in several previous papers^7,15,16,37^. The attenuation goes to zero as the GWAS sample becomes large and *ρ* asymptotes to 1. The farther *ρ* is from 1, the more *β* and ***δ*** are attenuated toward zero relative to *β*_*c*_ and ***δ*** *c*. The term Corr (***x***_*i*_ ***γ, x***_*i*_ ***μ***) captures a distinct source of measurement error that is present whenever the PGI is constructed from a GWAS with imperfect controls. Even though this type of measurement error is not classical, *β* and ***δ*** are further attenuated toward zero in proportion to Corr (***x***_*i*_ ***γ, x***_*i*_ ***μ***).

Equation (4) generalizes and unifies earlier analyses by Trejo and Domingue^18^ and Becker et al.^7^. Under the assumption Corr (***x***_*i*_ ***γ, x***_*i*_ ***μ***) =1 which corresponds to the case where the only source of bias is measurement error in the PGI, Equation (4) specializes to the formula derived in Becker et al.^7^. Under the assumption *ρ* =1 which corresponds to the case where the feasible regression uses a PGI with weights ***w = μ*** rather than the family-based weights ***w = γ*** that are used for the PGI in the ideal regression, Equation (4) coincides with a formula derived by Trejo and Domingue. In many empirical settings, it is plausible that both (i) estimation error in the SNP weights (*ρ* < 1) and (ii) differences between and (ii) differences between the true associational and family-based parameters (***μ* ≠ *γ*)** contribute meaningfully to the bias. Equation (4) provides a unified correction for both error sources in such cases.

A recent paper^33^ estimated the correlation between the causal and associational additive SNP factors, Corr (***x***_*i*_ ***γ, x***_*i*_ ***μ*)**, for a range of phenotypes. The correlation takes the value 1 if there is no confounding from gene-environment correlation that biases ***μ*** away from ***γ***^b^. It can also take the value 1 when ***θ*** is proportional to ***γ***. In general, however, we anticipate the correlation to be between 0 and 1^12,33^. For a range of Repository PGIs, using estimates from Tan et al.^38^, Table 3 calculates an estimate of the proportional bias term, Corr (***x***_*i*_ ***γ, x***_*i*_ ***μ***) / *ρ* We estimate the smallest attenuation for height, roughly 5%, for which both *ρ* and Corr (***x***_*i*_ ***γ, x***_*i*_ ***μ***) / *ρ* are close to 1. We estimate the largest attenuation for age at which a woman had her first child, almost 60%, for which there is substantial estimation error in the PGI (*ρ* ≈ 2) and non-trivial confounding in ***μ*** (Corr(***x***_*i*_ ***γ, x***_*i*_ ***μ***) ≈ 0.88, although with a large standard error).

In summary, our derivations imply that, in many applications, the coefficients from a family-based PGI study will be attenuated toward zero by a factor of Corr (***x***_*i*_ ***γ, x***_*i*_ ***μ***) / *ρ* The additional bias beyond measurement error arises because for many phenotypes, PGIs that are sufficiently predictive for use in applications at present—such as those in the Repository—are only available from standard GWAS designs that do not fully control for confounders. Consequently, relative to the coefficients that would be obtained using a noiseless PGI constructed from causal genetic effects, the coefficients from a family-based PGI study suffer from two sources of measurement error in the PGI weights: they are based on associations rather than causal effects, and they are estimated in a finite sample.

## Discussion

This paper introduces version 2.0 of the Polygenic Index Repository, a substantially expanded resource of PGIs designed to support research across the behavioral and social sciences. The new release is based on an improved methodology for estimating SNP weights, increases the number of covered phenotypes from 47 to 61, and expands the number of participating cohorts from 11 to 20. Where possible, PGIs are constructed from updated GWASs that incorporate previously unavailable data, further boosting prediction accuracy. To minimize technical and administrative barriers, we continue to distribute the PGIs as ready-to-use variables via participating datasets.

Another major innovation is that in cohorts with first-degree relatives, we supply parental PGIs based on imputed parental genotype data. These allow researchers to estimate PGI associations conditional on parental PGIs—a strategy that leverages random genetic variation from meiosis and is widely regarded as a credible approach to strengthening causal inference. We use the parental PGIs in two illustrative applications.

Alongside these empirical advances, we develop a theoretical framework to clarify the interpretation of coefficients from family-based PGI regressions that control for parental PGIs. Our analysis unifies and extends previous work, showing that attenuation in main-effect and PGI-by-environment interactions arises from two distinct sources of measurement error: sampling error in the GWAS-based SNP weights, and bias due to divergence between SNP weights from a population-based GWAS and SNP weights from a family-based GWAS. We derive an analytic correction formula that is straightforward to apply and that can be used to facilitate cross-study comparability, e.g., of gene-by-environment interaction estimates. We caution, however, that our analytic correction formula does not account for assortative mating.

As researchers pursue applications, it is essential to recall that “causal” does not mean “immutable.” Genetic influences often operate through modifiable environmental pathways. To avoid overly deterministic interpretations, PGI associations should be analyzed and presented with appropriate nuance. We have updated the repository’s User Guide to highlight interpretational issues we view as especially important (Supplementary Methods).

A major limitation of the current release is that it remains restricted to individuals of European genetic ancestries. To date, except for a few phenotypes, sufficiently large GWAS from other ancestry groups have not been available to generate PGIs that meet our inclusion criteria for predictive power. Addressing this imbalance is a priority for future releases. Another priority for future releases is to supply PGIs based on weights from family-based GWASs. While analysis of such PGIs will still suffer from errors-in-variables bias due to sampling error in the SNP weights—indeed much more so than the PGIs currently available because they will have much greater estimation error—they will not suffer from the other bias described here because their SNP weights will be (noisy) estimates of the ideal SNP weights.

## Methods

### I. Repository Cohorts

There are 20 cohorts participating in the current release of the Repository. Eleven of these participated in the initial release, and nine have joined subsequently. For two of the cohorts that participated in the first release, Estonian Biobank and Swedish Twin Registry, we received additional samples. Table 2 lists all cohorts participating in the current release, their sample sizes and some additional details.

#### Genotyping and Imputation

Genotyping was performed using a range of commercially available arrays. Cohorts were encouraged to upload genotypes imputed against HRC^39^ or TOPmed^40^ imputation panels. From some cohorts, we received only genotyped SNPs. We imputed these data against either the HRC (version 1.1) or the TOPmed reference panels using the Michigan/TOPmed Imputation Servers. Supplementary Table 6 provides study-specific details on the genotyping arrays, pre-imputation quality control filters, imputation software used, and reference panels.

#### Subject-level QC

We restricted the samples to individuals that cluster together with the EUR subsample of the third phase of 1000 Genomes Project^41^ along the first four principal component axes. To obtain the first four principal components (PCs), we largely followed the same procedure as Becker et al.^7^. We first converted the imputed genotype dosages for HapMap3 SNPs into hard calls. Then, we merged the data with all samples from the third phase of the 1000 Genomes Project. We dropped SNPs that had a call rate less than 99% or minor allele frequency less than 1% in the merged sample. We computed the PCs using the 1000 Genomes sample and projected the remaining individuals onto this PC space. The next step we applied differs from Becker et al.^7^ in that we did not rely on visual identification of individuals that cluster together with the 1000 Genomes EUR sample in PC plots. Instead, we identified an individual as having a European genetic ancestry if for each of the first four PCs, they are within 5 standard deviations away from the mean of that PC in the 1000 Genomes EUR sample. We applied this new criterion to all new cohorts, as well as to the cohorts from the initial release that we had to re-analyze because we lost and re-gained access due to changes in access procedures or similar issues. Therefore, some of the cohorts from the initial release have slightly different (smaller) sample sizes in the current release. Supplementary Table 6 lists, for each cohort, the size of the sample for which we made PGIs (Column E).

#### Creation of PCs in Repository Cohorts

To obtain PCs in the Repository cohorts, we first applied some SNP-level filters to the genotype data. We excluded SNPs with imputation accuracy less than 70% or minor allele frequency less than 1%, as well as SNPs in long-range LD blocks. We pruned the remaining SNPs using a 1Mb rolling window (incremented in steps of 5 variants) and an *r*^2^ threshold of 0.1. Using this set of approximately independent SNPs, we estimated a genetic relatedness matrix and identified all pairs of individuals with a relatedness coefficient greater than 0.05 as calculated by in Plink1.9^42^. We excluded one individual from each pair, calculated the first 20 PC loadings in the sample of unrelated individuals, and projected the sample onto the PC space using Plink1.9. This procedure was the same as the initial release of the Repository, with the exception of the list of long-range LD blocks that we excluded in the first step. In the current release, we excluded a larger set of genomic regions provided by the plinkQC^43^ R package.

### II. Summary Statistics UKB GWAS

We conducted GWAS in UKB for 22 new phenotypes that we intended to include in the second release of the repository (Supplementary Table 2, column C), one phenotype that was included in the first release but for which we had not run a GWAS in UKB (column D), and updated the GWAS for 10 of the first release phenotypes (column E) with an improved phenotype definition and/or additional data. For some additional phenotypes^c^, we ran GWAS in UKB but it was not possible to meta-analyze these with the 23andMe, Inc., and/or published GWAS because of the mean *χ*^2^-statistic being too low (< 1.02) for MTAG to produce stable estimates. Therefore, these GWAS were not used in further analyses.

For each phenotype, we conducted three GWAS in three equal-sized partitions of UKB so that we could use summary statistics from two partitions when making PGIs for the remaining partition^d^. The partitioning rule is described in detail in Becker et al., but briefly, the first partition (UKB1) included all individuals with brain scan data, all pairs of relatives up to second degree and some pairs of third degree; the second partition (UKB2) included all remaining third-degree relatives and randomly selected individuals with no third degree or closer relatives; and the third partition (UKB3) was composed only of participants with no third degree or closer relatives.

Of the 22 new phenotypes, eight were biomarkers, three were fertility or sexual development outcomes, five were diseases, four were mental disorders, and two were substance use phenotypes. For the health phenotypes among the ones with updated GWAS, the main change in phenotype definition was the incorporation of primary care data, which in most cases resulted in a higher number of cases but lower number of controls. Therefore, for most of these phenotypes, the sample size decreased. One exception was migraine, for which we included additional variables in the phenotype construction, leading to a higher number of cases and controls. For the three non-health phenotypes, i.e., educational attainment (EA), number ever born (men) and number ever born (women), the sample sizes decreased slightly. For EA, we used the updated phenotype definition from Okbay et al.^44^. For the number ever born phenotypes, we imposed an age cutoff (55 for men, 45 for women) following Mathieson et al.^45^ that we previously had not imposed. The phenotype definitions and sample sizes for all 33 phenotypes for which we ran new or updated GWAS in UKB, as well as the definitions from the first release of the Repository for the 11 phenotypes with updated GWAS, can be found in Supplementary Table 2.

We applied the same sample-level quality control filters described in Becker et al. prior to running the GWAS and restricted the sample to individuals of European genetic ancestries, defined as the first genetic PC provided by UKB being greater than 0 and individual self-reporting to be of “British”, “Irish”, or “Any other white background”. After obtaining the phenotypes as described in Supplementary Table 2, within each partition, we residualized the phenotypes on sex, a third-degree polynomial in birth year (defined as (*birthyear* - 1900)/10), their interactions, 106 genotyping batch dummies, and the first 40 of the PCs released by the UK Biobank. We ran the GWAS on these residuals using mixed-linear models implemented by the software BOLT-LMM^46^ with the settings used by Becker et al.

#### 23andMe, Inc. GWAS

The first release of the repository used GWAS summary statistics from 23andMe for 37 phenotypes. In the current release, we use an additional nine GWAS conducted by 23andMe in samples of research-consented participants with European genetic ancestries. All of these GWAS have been previously published. For height, the GWAS was run separately in samples of men and women. The remaining phenotypes were analyzed in pooled-sex samples, using sex as a covariate. Supplementary Table 7 provides an overview of the nine GWAS, including phenotype definitions, association models, sample sizes, case prevalence for binary phenotypes, and the original publications. For details on 23andMe’s genotyping and imputation, see Supplementary Tables 17 and 18 in Lee et al.^47^

#### Quality control of summary statistics

We applied a uniform set of quality-control filters to all UKB and 23andMe GWAS described above, as well as to the published GWAS. The quality control pipeline is described in Becker et al. and includes checks for strand misalignment, allele mismatch, chromosome and base pair position concordance, and allele frequency discrepancies with the Haplotype Reference Consortium reference panel (r1.1)^39^; filters for minor allele frequency, imputation accuracy, and standard-error outliers; and visual inspection of diagnostic plots. Supplementary Table 8 summarizes the number of SNPs dropped in each group of filters in the files that passed all diagnostic checks. After applying the quality control pipeline, for each phenotype, we estimated the heritability of the available GWAS as well as the pairwise genetic correlations between them using the LDSC software package^48^ (Supplementary Tables 9 and 10). This final check allowed us to confirm that the phenotype coding was in the same direction across 23andMe, UKB, and published studies, as well as informed us about the amount of signal in each GWAS and about the homogeneity of phenotype definitions across them.

#### Input GWAS

The input GWAS that the weights are based on are obtained by meta-analyzing GWAS summary statistics from up to three sources: UKB, 23andMe and previously published GWAS on the phenotype. For most phenotypes, input GWAS varies across repository cohorts due to sample overlap. When the cohort is included in the largest previously published GWAS for that phenotype, instead of that GWAS, we use the largest published GWAS that does not include the cohort. If the cohort is included in all previously published GWAS on the phenotype, we use summary statistics from the remaining two sources (UKB and 23andMe) for that cohort.

For the three UKB partitions, we use the same strategy but with a nuance. The input GWAS for each partition is based on up to three sources (fewer if one or more is unavailable): summary statistics from the remaining two partitions of UKB, 23andMe, and the largest previously published GWAS that does not include UKB. For several phenotypes (e.g., cigarettes per day), the authors of the published GWAS have made publicly available a version of the GWAS summary statistics that excludes UKB; for those phenotypes, we use that version of the summary statistics. For other phenotypes, we use an earlier publicly available GWAS that does not include UKB in the meta-analysis. We conduct the meta-analyses in MTAG^19^, allowing for sample overlap and fixing genetic correlations at unity. We allow SNP heritability to vary across input files. Additionally, we make the following changes to the default settings of MTAG:

- We turn off the filter that drops SNPs with *P*-values equal to 0. This change was prompted by the fact that for several phenotypes, we had SNPs with extremely small *P*-values in the input files which were perceived as 0 by MTAG due to a numerical precision issue. Therefore, these SNPs, which are the most strongly associated SNPs with the respective phenotypes, were being excluded from the analyses. Turning off the *P*-value filter allowed us to keep those SNPs.
- We use the --use_beta_se option, which tells MTAG to obtain the *Z*-statistics using beta and standard error instead of the *P*-value. This was required due to the previous change that kept SNPs with *P*-values perceived as 0.
- For two phenotypes, Alzheimer’s disease and prostate cancer, we ran MTAG using SNP identifiers of the format “Chromosome:Base Pair” rather than rs-numbers since the rs-number overlap across the input files for these phenotypes was poor.
- MTAG uses LD score regression to estimate various parameters. By default, it runs LD score regression without any filter for maximum association *χ*^2^-statistic (the LDSC default filter is 30). For the blood lipid phenotypes other than triglycerides, LD score regression failed to produce reasonable results with no *χ*^2^ filter. Therefore, for these phenotypes, we set the filter to 1000.

For some phenotypes that had input files with very large sample sizes, although there was no sample overlap across the input files, MTAG reported bivariate LD score regression intercepts that were considerably larger than 0 and corrected for the supposed overlap by deflating the *χ*^2^-statistics. Yengo et al.^49^ show that this behaviour from LD score regression is expected when GWAS sample sizes are large and the traits are highly heritable. In fact, in the absence of sample overlap, the bivariate LD score regression intercept is proportional to the geometric mean of the sample sizes of both GWAS and the genetic covariance between the traits. To deal with this overcorrection of the *χ*^2^-statistics, we fixed the intercepts at 0 by using the –-no-overlap option for these phenotypes. This adjustment was required for the input GWAS for all blood lipids except for triglycerides, BMI, height and educational attainment^e^.

Supplementary Table 11 shows, for all phenotypes, the different versions of the input GWAS along with sample sizes, the cohorts that have sample overlap with the input GWAS (column I), and which cohorts the input GWAS is used for (column J). For example, there are five versions of the input GWAS for BMI, coded BMI1 - BMI5. BMI1 is the largest GWAS and includes UKB, and Locke et al.^50^. BMI1 is used for all cohorts except for Estonian Biobank, Finnish Biobanks, 1958BC, STR and the UKB partitions, which have sample overlap with BMI1. For Estonian Biobank, Finnish Biobanks, 1958BC and STR, the overlap is with Locke et al. For these cohorts, we use a version of the GWAS, BMI2, that is only based on UKB. The UKB partitions have no overlap with Locke et al., but for each partition, we need to remove that partition from the input GWAS and include the remaining two. Therefore, for UKB1, we use BMI3, which is based on UKB2, UKB3 and Locke et al. BMI4 and BMI5 are obtained similarly and used for UKB2 and UKB3, respectively.

As a final check to ensure the phenotypes are coded in the right direction, we estimated the genetic correlation between the largest input GWAS for each pair of phenotypes. These results can be found in Supplementary Table 12.

### III. Constructing Repository PGIs Criterion for Inclusion in Repository

We use the same criterion as the initial release of the Repository to include a PGI in the second release: the theoretically expected out-of-sample predictive power has to be greater than 1%. (The actual predictive power may differ due to sampling error in the predictive power or estimation error in the parameter estimates we use to calculate our theoretical expectation.) We calculate the theoretically expected out-of-sample predictive power using the following formula^51^:

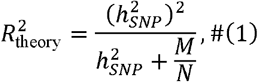

where 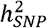 is the phenotype’s SNP heritability, *M* is the effective number of independent SNPs which we assume to be equal to 60,000^52^, and *N* is the GWAS sample size for the phenotype.

For each phenotype, we estimate 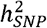 with LD score regression^53^ using the GWAS with the largest sample size (i.e., the GWAS numbered 1 in Supplementary Table 11, e.g., BMI1, HEIGHT1). In LD score regression, we use the “--two-step inf” option to avoid excluding SNPs with large association *χ*^2^-statistics for all phenotypes except for the blood lipids other than triglycerides. For those, we use a *χ*^2^ cutoff of 1000 (see the previous section for a discussion). Prior to estimating heritability, we calculate the effective sample size as

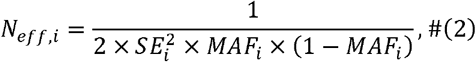

where *SE*_*i*_ is the standard error of the estimated effect size for the SNP *i* and *MAF*_*i*_ is the minor allele frequency. We subsequently filter out SNPs with *N*_*eff*_<0.8 × median(*N*_*eff*_) We set the sample size for all remaining SNPs to the GWAS-equivalent-*N* reported in the MTAG output. We also use this sample size as *N* in the 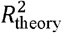 formula. Supplementary Table 13 lists the number of SNPs excluded by the effective sample size filter as well as the GWAS-equivalent-*N* for each input GWAS.

Supplementary Table 1 shows the heritability estimates and expected predictive power for all phenotypes.

With the exception of Alzheimer’s disease, childlessness (men, women, pooled) and number ever born (pooled), all new phenotypes satisfy the 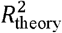 criterion. For Alzheimer’s disease we waive this criterion because heritability is estimated using HapMap3 SNPs, which insufficiently capture the effect of the APOE gene. In addition, two phenotypes for which single-trait PGIs did not meet the inclusion criterion in the first release, Allergy-Pollen and COPD, pass the cutoff in the current release with their updated GWAS. In total, there are 60 phenotypes (not including Alzheimer’s disease) in the second release that satisfy the criterion. Of these, 22 are new, 19 have updated GWAS, and 20 are based on the same GWAS as the initial release.

#### Constructing PGIs

In the initial release of the Repository, we used LDpred^20^ to adjust the SNP weights for LD. Here, we used the SBayesR methodology implemented in the GCTB software^54^. Instead of imposing a point-normal mixture distribution as the prior on SNP effects like LDpred, SBayesR imposes a flexible finite mixture of normal distributions. To estimate the posterior mean SNP effects adjusted for LD, SBayesR requires a reference data set containing LD estimates between SNPs. For this purpose, we used the LD matrix for the 2,865,810 pruned common variants from the full UKB European-genetic-ancestry (*N*□≈□450,000) data set from Lloyd-Jones et al.^21^. Following the recommendation by Lloyd-Jones et al.^21^, we excluded 3,638 SNPs in the MHC region (Chr6 : 28-34Mb) to improve model convergence. Because SBayesR is prone to convergence problems when sample size varies greatly across SNPs, we also excluded SNPs that have an effective sample size, calculated as in equation (2), smaller than 80% of the median effective sample size. After imposing the filter, we set the sample size of all SNPs to the GWAS-equivalent *N* reported in the MTAG log file. Supplementary Table 13 shows the number of SNPs dropped by the effective sample size filter from each input GWAS, as well as the minimum, maximum and median sample sizes and the GWAS-equivalent *N*.

We ran SBayesR assuming a four-component normal mixture model, with initial mixture probabilities π = (0.95,0.02,0.02,0.01), and γ = (0.0,0.01,0.1,1), where γ contains the scaling factors for the variance of the distribution of genetic effects for each of the four components. The MCMC was run for 10,000 iterations with 2,000 taken as burn-in. We experienced no convergence issues with these settings. However, for a few input GWAS^f^, we observed that although the SBayesR algorithm converged and the software reported no error messages, the reported heritability estimate seemed unreasonable. For these GWAS, we ran SBayesR with the “—impute-n” and “--robust” flags, which solved the issue.

We obtained the adjusted weights for the overlapping SNPs between the ~2.9 million SNPs available in the LD matrix and the input GWAS after the filters described above. This, too, is a deviation from the initial release of the Repository where the set of variants in the PGIs were restricted to HapMap3 SNPs. This decision was partly based on the finding by Lloyd-Jones et al.^21^ that SBayesR achieved the highest prediction accuracy for height in their dataset when using the ~2.9 million SNPs. Additionally, for the phenotypes available in the initial release of the Repository, we ran SBayesR using both sets of SNPs (HapMap3 and ~2.9 million) and compared the prediction accuracy in our three validation cohorts. We found that PGIs based on the set of ~2.9 million SNPs outperformed those based on HapMap3 SNPs on average.

The PGIs were calculated in Plink2^42^ by multiplying the genotype dosages at each SNP by the corresponding estimated posterior mean calculated by SBayesR, and then summing over all included SNPs.

#### Between-Family Prediction Analyses

We assessed the predictive power of the PGIs in three Repository cohorts: HRS (*N* = 12,670; 57% female; median birth year = 1941), WLS (*N* = 8,478; 52% female; median birth year = 1939), and UKB (third partition; *N* = 148,662; 54% female; median birth year = 1950). Supplementary Table 14 describes the phenotype measures in HRS and WLS for the new phenotypes added in the current release. The new or updated UKB phenotype measures are described in Supplementary Table 2. Remaining phenotype measures (i.e. those that were included in the first release of the Repository) can be found in Supplementary Tables 5 and 12 of Becker et al.^7^. Descriptive statistics for all phenotypes in the validation cohorts can be found in Supplementary Table 15.

We followed the same procedure as Becker et al.^7^ to construct the phenotypes. If a single measurement was available, we residualized the phenotype measure on a second-degree polynomial in age or birth year, sex, and their interactions. If multiple measurements in time were available, we did either of the following two: for quantitative phenotypes, we did the same residualization in each wave and took the mean across waves; for case-control phenotypes, we took the maximum across waves and then residualized on a second-degree polynomial in birth year, sex, and their interactions.

Our measure of predictive power is the incremental *R*^2^, calculated as the increase in the coefficient of determination when the PGI is added to a regression of the phenotype on the first 10 PCs of the genetic data in HRS and WLS, and the first 20 PCs and 106 genotyping batch dummies in UKB. Supplementary Table 3 shows the results alongside the incremental *R*^2^’s from Becker et al.^7^ for phenotypes included in incremental the first release to demonstrate the improvement in the PGIs. The 95% confidence intervals around the *R*^2^’s are obtained by bootstrapping with 1000 repetitions.

### IV. Parental PGIs

#### Imputation of Missing Parental Genotypes

We impute the genotypes of missing parent(s) based on Mendelian laws using the software tool *snipar*^12^. Prior to imputation, as a first step, we used *vcftools*^55^ v.4.2 (for vcf files) or *qctool* v.2.0.7 (for bgen files) to restrict the sample to individuals of European genetic ancestries as defined above, and remove indels, multiallelic variants, SNPs with imputation quality (INFO score or R^2^) less than 0.99, average posterior call (if available) less than 0.99, minor allele frequency less than 1% as well as SNPs out of Hardy-Weinberg equilibrium, for which we imposed various *P*-value cutoffs depending on the sample size of the cohort. We used a stringent imputation quality threshold because previous analyses indicated that all but the highest quality imputed variants show deviations from expected Mendelian relations between relatives^33^. For two cohorts, SOEP-G and TwinLife, we relaxed the imputation accuracy and average posterior call cutoffs to 0.95 since the number of SNPs remaining after filters were too few with the 0.99 cutoffs. Next, we converted the files into two different formats: “bgen” if the genotypes were phased (using *qctool*, unless the original files were in bgen format already) since bgen format preserves phasing information and is accepted by *snipar*, and “plink (bed/bim/fam)” (using plink2). We merged the per-chromosome plink files in plink1.9 and used this as input for KING^56^ to estimate the relatedness between individuals. In order to infer identity-by-descent (IBD) segments shared by siblings, we first pruned the variants in the per chromosome plink files using a 1Mb rolling window (incremented in steps of one variant) and an *R*^2^ threshold of 0.3. With these files, the KING output, and a file containing the age and sex of the individuals in the sample, we used *snipar* (ibd.py) to infer the IBD segments. When parent-offspring pairs were available in the sample, we let *snipar* calculate the per-SNP genotyping error probability and used the default cutoff of 0.01 for maximum allowed genotyping error. In cohorts with no parent-offspring data, we set the genotyping error probability to 0.001 using the -–p_error flag. Finally, using the bgen files for cohorts with phased data and plink files for those with unphased data, the KING output, the ibd segments inferred by *snipar*, and the file containing the age and sex of the individuals in the sample, we imputed the missing parental genotypes with *snipar* (impute.py).

For the UKB, we had access to phased haplotypes from the genotyping array and unphased imputed genotype dosages. In order to be able to utilize the phasing information without having to restrict the set of SNPs to those available only on the genotyping array, we conducted two separate parental genotype imputations: one using phased haplotypes from the genotyping array and another using the unphased dosages. For both imputations, we started by filtering out heterozygosity outliers, samples with sex chromosome aneuploidy, excess relatives, or excess genotype missingness. Next, we imposed the variant-level quality control filters described above. We used the filtered phased haplotypes for IBD inference without any LD-pruning. These inferred IBD segments were used in both imputations. The rest of the imputation followed the pipeline described above. Finally, we merged the two sets of imputed parental genotypes, with the one using phased haplotypes taking precedence when the SNP was available in both imputations.

The specific QC cutoffs we used for each cohort, the number of genotyped parent-offspring or sibling pairs, the number of parent-offspring trios available after imputation and the number of SNPs imputed for the parent(s) are summarized in Supplementary Table 16.

#### Constructing parental PGIs

We constructed PGIs for the genotyped or imputed parents using *snipar* (pgs.py). The *snipar* pgs.py script requires as input (i) a file set containing the observed genotypes for the proband individual and their genotyped parents (if available), (ii) a file set containing the imputed genotypes for the imputed parents (i.e., the .hdf5 files produced by the impute.py script explained in the previous section), and (iii) a file containing the SNP weights for the phenotype. For cohorts with phased genotypes, we used, as (i), the filtered per chromosome bgen files we had used as input to the impute.py script for the parental genotype imputation. For cohorts with unphased genotypes, we used the filtered per-chromosome plink bed/bim/fam files described in the previous section. These two sets of files contain the same set of SNPs. However, *snipar* does not accept unphased genotypes in bgen format, which led us to use different formats for phased and unphased data. As (iii), we used the LD-adjusted SNP weights from SBayesR that we describe above under “Constructing PGIs”. We used the default option in *snipar* to apply assortative-mating adjustment to the imputed parental PGIs if the correlation between the maternal and paternal PGIs could be estimated with sufficient precision.

The *snipar* pgs.py output contains, in addition to the family and individual identifier for the proband and their genotyped parents (if available), a PGI for the proband and either a single “parental” PGI, or separate “maternal” and “paternal” PGIs. The former situation occurs when there are no genotyped parent-offspring pairs in the data. The imputation of separate maternal and paternal genotypes from a sibling pair is not generally possible. Therefore, the maternal and paternal PGIs are set to the same value by *snipar*. If there are no genotyped parent-offspring pairs in the data, the imputed maternal and paternal PGIs are the same, so *snipar* outputs a single “parental” PGI — the sum of paternal and maternal PGIs — to avoid collinearity issues in downstream analyses. It is also important to note here that the “proband” PGI in the output is not identical to the regular PGIs that we created for the Repository. The “proband” PGI is based on the same set of SNPs as the parental PGIs, i.e., the intersection between the weights and the set of SNPs that were imputed for the parents. These are smaller sets of SNPs that remain after strict quality control filters (see the previous section). Although these proband PGIs will generally be less predictive because they are based on fewer SNPs, we recommend using these PGIs in analyses that control for parental PGIs to ensure that the parent and offspring PGIs preserve expected Mendelian relations required for the expected properties of family-based analyses to apply. Supplementary Table 17 provides, for each cohort and phenotype, the number of SNPs included in the *snipar* PGIs, as well as the correlation between maternal/paternal/parental and proband PGIs and between the *snipar* proband PGI and the regular Repository PGI.

#### Within-Family Prediction Analyses

We utilized two of our Repository cohorts with family data, WLS and UKB, to compare the “population association” and “causal effect” of PGIs across various phenotypes. The “population association” of a PGI, denoted *ψ*, is the coefficient from a regression of an individual’s phenotype on that individual’s PGI:

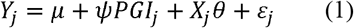

where *PGI*_*J*_ is the PGI of an individual (i.e., proband) *j*; μ is the intercept; *X*_*j*_ represents a vector of individual-level covariates (sex, age, age^2^, their interactions with sex, and the top 20 PCs); and *ε*_*j*_ is the residual term. In UKB, we use the third partition, UKB3, which does not include any related individuals, to obtain the population effect. In WLS, we use a subset of the cohort that contains no related individuals (no siblings). We ran logistic regression for binary phenotypes and linear regression for continuous phenotypes. To interpret effect sizes from logistic regression more intuitively, we transformed 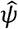 into an odds ratio.

In UKB, the “causal effects” of PGIs, denoted *δ*, were estimated in the first-degree relatives subsample of UKB1 using the following regression model:

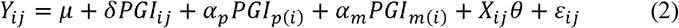

Where *PGI*_*p*(*i*)_ and *PGI*_*m*(*i*)_ are the PGIs of the father and mother in family *i*, respectively.

In WLS, the “causal effects” of PGIs were estimated in the sibling subsample using a slightly modified version of equation (2):

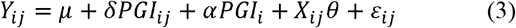

We employed Equation (2) for UKB as we have the observed or imputed PGIs for both parents of the proband. In the WLS cohort, however, we implemented Equation (3), which controls for the parental PGI (sum of paternal and maternal PGIs) (*PGI*_*i*_) imputed from siblings as we do not have any genotyped parent-offspring pairs. *α*_*p*_ and *α* _*m*_ in Equation (2) and *α* (which is equal to the average of the coefficients on the maternal and paternal PGIs)^12^ in Equation (3) capture indirect genetic effects, gene-environment correlations arising from population stratification, and assortative mating. All other terms in Equation (2) are analogous to those in Equation (1). Note that *δ* exclusively captures causal effects because the variation in an individual’s PGI, once parental PGIs are controlled for, is due to random Mendelian segregation during meiosis^25^.

For non-binary phenotypes, *δ* and *α*_{*p, m*}_ were estimated using *snipar* (pgs.py), which implements a mixed linear model that accounts for relatedness. However, *snipar* does not have an option for an equivalent logistic regression. Therefore, for binary phenotypes, we estimated a generalized linear mixed model with a logistic link using the -glmer function from the -lme4 R package, including a random intercept for family ID to account for within-family correlation. As in the models evaluating “population associations”, we standardized the PGIs before running all regressions and transformed the coefficients from the logistic regressions into odds ratios.

We estimated the population associations in separate samples from the ones used for estimating the causal effects to obtain independent estimates of *δ* and *ψ*. This allowed us to calculate standard errors for the ratio 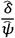 using the delta method, without needing the covariance between the two estimates. (The alternative of bootstrapping the ratio was not feasible due to the computational demands of fitting mixed linear models.) To ensure the two samples were comparable, we also estimated population associations in the first-degree-relatives subsamples of UKB1 and WLS. These estimates are very similar to those in the unrelated subsamples.

Supplementary Table 4 shows the estimated population associations and causal effects for the two cohorts, as well as the causal effect to population association ratio. For binary phenotypes, we obtained the ratio using logistic regression coefficients (prior to transforming them into odds ratios). To obtain standard errors for the ratio of causal effect to population association estimates, we used the delta method:

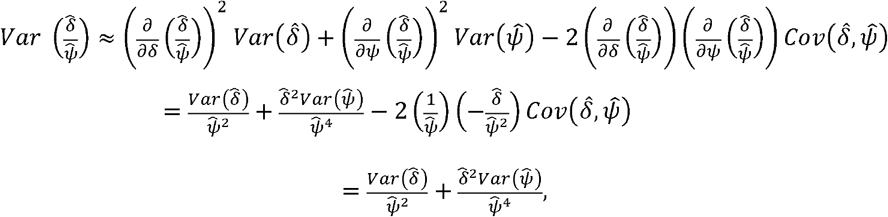

where the last equality follows because (i) *δ* and *ψ*. are treated as fixed (but unknown) parameters in our frequentist framework, and hence 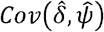 can be non-zero only if 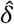 and 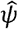 have correlated estimation error; and (ii) *δ* and *ψ*. are estimated in two independent samples and therefore have independent estimation error. Thus, we compute the standard errors as:

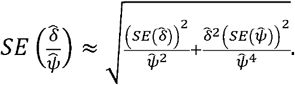

### V. Analysis of PGI Effect Heterogeneity

To provide an illustrative application of parental PGIs, we revisit the analysis of PGI effect heterogeneity across subgroups in the UK Biobank (UKB) originally conducted by Mostafavi et al.^57^.

The main text highlights three primary differences between our approach and theirs; here, we note two more. First, whereas Mostafavi et al. analyzed BMI stratified by four age quartiles, we divide the prediction sample into two age groups using the median age as the cutoff. Second, the prediction samples differ. For their analysis of diastolic blood pressure by sex, Mostafavi et al. used a mixed-sex prediction sample comprising 20,000 men and 20,000 women. For their BMI-by-age analysis, they sampled 10,000 individuals from each age quartile. In both cases, individuals were randomly selected from among “White British”^58^ UKB participants who were not included in the GWAS used to estimate the relevant PGI weights. In contrast, we construct PGIs using a meta-analysis of GWAS summary statistics from the second and third UKB partitions along with external datasets (see “Input GWAS”) to maximize prediction accuracy. Our prediction sample for all analyses consists of the first-degree relatives subsample from the first partition of the UKB (*N* = 39,687), all of whom have observed or imputed parental genotypes.

#### Population Associations

To estimate population associations, we fit linear mixed models that include an interaction term between the PGI and the relevant subgroup indicator:

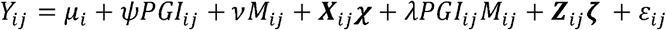

where *Y*_*ij*_ is the phenotype of individual *j* in family *i*; *μ*_*i*_ is a random intercept for family *i*; *PGI*_*ij*_ is the standardized PGI of focal individual *j* in family _*i*_; *M*_*ij*_ is the subgroup indicator; ***X*** _*ij*_ is a vector of mean-centered individual-level covariates (age and the first 20 PCs for diastolic blood pressure and additionally sex for BMI); ***Z*** _*ij*_ is the vector of all two-way interactions between (i) *PGI*_*j*_ and the covariates in ***X*** _*ij*_ and (ii) between *M*_*ij*_ and the covariates in ***X*** _*ij*_; and *ε*_*ij*_ is the portion of the residual that is drawn independently for members of the same family. The phenotype variable *Y*_*ij*_ is standardized within each subgroup defined by *M*_*ij*._

#### Causal Effects

To estimate causal effects, we additionally control for parental PGIs:

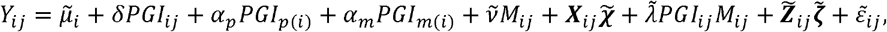

where *PGI*_*p*(*i*)_ and *PGI*_*m* (*i*)_ are the standardized PGIs of the father and mother of focal individual *j* in family *i*, and 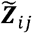 additionally includes all two-way interactions between (i) each parental PGI (*PGI*_*p* (*i*)_ and *PGI*_*m* (*i*)_ and *PGI*_*ij*_ and (ii) between each parental PGI and *M*_*ij*_. (The coefficients on the parental PGIs are separately identified by the small number of cases where at least one parental PGI is observed rather than imputed.)

#### Comparing Standardized Coefficients

For diastolic blood pressure, *M*_*ij*_ is an indicator equal to one for males; for BMI, it is an indicator equal to one for individuals above the median age. The population association in the baseline group defined by *M*_*ij*_ = 0 is given by the coefficient on the *PGI* (*ψ*) in the first model above. The association in the comparison subgroup defined by *M*_*ij*_ = 1 is equal to *ψ* + *λ* where *λ* is the coefficient on the interaction between the subgroup indicator *M*_*ij*_ and the PGI. We assess heterogeneity by examining if the ratio of the two population associations, *ψ*/(*ψ* + *γ*), is distinguishable from 1. In the second model with controls for parental PGIs, *δ* is the causal effect in the baseline group defined by *M*_*ij*_ = 0 and 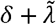 is the causal effect in the comparison subgroup. The ratio of the causal parameters is given by 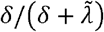 In our empirical analyses, we estimate the two ratios by plugging in our estimates of the population parameters into the ratio formula, and we use the delta method^59^ to obtain standard errors for each estimated ratio.

#### Comparing Incremental *R*^*2*^

To compare incremental *R*^2^*s*, we estimate one linear mixed-effects model for each subgroup. For the population associations, we split the sample into two based on the value of *M*_*ij*_ and estimate the following two models:

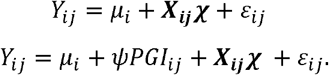

We calculate the incremental *R*^2^ as the difference in the coefficient of determination between the two. For the causal effect, we estimate incremental *R*^2^ as the difference in *R*^2^ of the following two models:

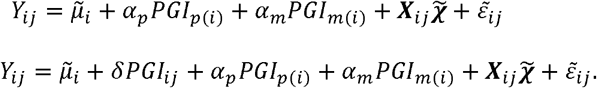

We obtain confidence intervals for these incrementa *R*^2^ estimates and their ratios via a non-parametric bootstrap. For each phenotype, we resample families with replacement within each subgroup 1,000 times.

We estimate the four models above in each iteration, generating bootstrap distributions for the incremental *R*^2^ in each group and the ratio of the incremental *R*^2^ values between groups. The 95% confidence intervals for both the individual incremental *R*^2^ values and their ratios are the 2.5^th^ and 97.5^th^ percentiles of their respective bootstrap distribution.

### VI. Categorization of BGA Annual Meeting Presentations

To categorize the 2,999 Behavior Genetics Association (BGA) meeting presentations from 2009 to 2024 for the main text Figure 1, we used a rule-based classification algorithm based on Becker et al.^7^, with the following improvements and modifications:

#### Algorithm Enhancements

1. General Preprocessing: any double spaces in the text were replaced with single spaces before applying the algorithm to ensure uniformity in keyword matching.
2. Categorization of “Twin/Family/Adoption” Studies:
  - Added Keywords: The following keywords were added to the algorithm to improve detection: “adult twins,” “twins,” “twin population,” “twin sample,” “twin and sibling pairs,” “twin design.”
  - Refined Criteria: The keyword “twin” was removed as a standalone term to reduce false positives.
  - A study was categorized as “Twin/Family/Adoption” if:
    - The title contained either “twin” or “adoption”, OR
    - The abstract mentioned the word “twin” at least twice, excluding mentions of “twin studies.”
3. Categorization of “Candidate-Gene” Studies:
  - Minor adjustments were made to keyword matching to enhance accuracy: Changed “tph” to “tph” and “akt” to “akt” (both beginning with a space) to avoid erroneous matches with non-gene-related text.
4. Categorization of studies “using within-family variation”: to construct the right panel of Figure 1, we introduced a new algorithmic component to identify PGI studies examining causal effects:
  - Keywords Added: “direct effects,” “direct-effects,” “direct genetic effects,” “sibling pairs,” “sib-difference,” “parental pgi,” “parental pgs,” “parental polygenic,” “within-sibship,” “within sibship,” “genetic nurture,” “indirect effects,” “indirect genetic effect,” “within family,” “within-family,” “tdt,” “polygenic transmission disequilibrium,” “trio,” “trios.”
  - Conditional Keyword Handling: If the text included keywords containing the substring “indirect”, the study was only categorized as “using within-family variation” if the keyword “direct” was also present to prevent misclassification.

#### Validation of the Algorithm

To evaluate the accuracy of our algorithm, we followed the procedure outlined in Becker et al., with modifications to account for the inclusion of studies that evaluate the causal effects of PGIs. The validation process was conducted in two stages:

##### 1. Validation of 2009–2022 Presentations

A total of 65 randomly sampled BGA presentations from 2009 to 2022 were independently categorized by expert raters. PGI studies using within-family variation were oversampled, constituting approximately 30% of the sample to ensure sufficient representation. Error rates were defined as the percentage of presentations where the algorithm’s categorization disagreed with the final expert consensus. The observed error rates were 9.2% for PGI studies, 9.2% for PGI studies using within-family variation, 16.9% for twin/family/adoption studies, and 7.7% for candidate-gene studies. These results indicate that the algorithm performs with reasonable accuracy, particularly for PGI and candidate-gene studies.

##### 2. Review and Recategorization of 2023–2024 Presentations

The dataset was subsequently updated to include presentations from 2023 and 2024. To assess the accuracy of the updated dataset, an expert rater conducted a targeted review of:

- All presentations across all years marked as “PGI studies using within-family variation.”
- All presentations from 2023 to 2024 marked as “PGI studies.”

Based on this review, the expert rater made the following recategorizations:

- 15 presentations were moved from “PGI studies using within-family variation” to “PGI studies” (not using within-family variation).
- 8 presentations were moved from “PGI studies” (not using within-family variation) to “PGI studies using within-family variation.”
- 5 presentations were removed from the “PGI studies” category.

These refinements ensured that the classification process remained robust, particularly for PGI studies utilizing within-family variation, which can significantly impact the interpretation of trends over time. Full details on the validation procedure and consensus-building process are available in Becker et al.

## Supporting information

Frequently Asked Questions

Supplementary Methods and User Guide

## Data availability

For how to access the Repository PGIs and other data from each participating dataset, see Supplementary Note; upon publication, an up-to-date list of participating datasets and data access procedures will be maintained at https://www.thessgac.org/pgi-repository. For each phenotype that we analyze, we report GWAS summary statistics and PGI (SBayesR) weights for all SNPs from the largest discovery sample for that analysis, unless the sample includes 23andMe. SNP-level summary statistics from analyses based entirely or in part on 23andMe data can only be reported for up to 10,000 SNPs. Therefore, if the largest GWAS for a phenotype includes 23andMe, we report summary statistics for only the genome-wide significant SNPs from that analysis. In addition, we report summary statistics for all SNPs from a version of the largest GWAS analysis that excludes 23andMe. These summary statistics and PGI weights can be downloaded from https://www.thessgac.org/pgi-repository upon publication. Researchers at non-profit institutions can obtain access to the genome-wide summary statistics from 23andMe used in this paper by completing the 23andMe Publication Dataset Access Request Form, available at https://research.23andme.com/dataset-access/.

## Code availability

The code for constructing PGIs and principal components will be available at https://github.com/aysuo/PGI_Repo upon publication.

## Acknowledgements

This research was carried out under the auspices of the Social Science Genetic Association Consortium (SSGAC). This research was conducted using the UK Biobank Resource under application number 11425. A.O. and R.Ah. were supported by the Swedish Research Council (grant number 2023-01343) awarded to A.O., R.Ah., and Felix Tropf.; A.O., R.Ah. and S.O. by the Swedish Research Council (grant number 2024-06499) to A.O., R.Ah., S.O., Abdel Abdellaoui and Eivind Ystrom.; A.O. by the Amsterdam UMC Fellowship to A.O.; D.J.B., D.C., P.T., and A.Y. were supported by Open Philanthropy and National Institute on Aging/National Institutes of Health grants R24-AG065184 and R01-AG042568 to D.J.B., R01-AG083379 to A.Y., and R00-AG062787 and R01-AG081518 to P.T.; U.V by Estonian Research Council’s personal research funding start-up grant PSG759; T.E. by Estonian Research Council’s team grant PRG1291; R.F.K. by the U.S. National Institute on Aging grants R01AG053217, R01AG077742 and U19AG051426. We thank the following consortia for sharing GWAS summary statistics: Reproductive Genetics (ReproGen) Consortium for age at first menses; Genetics of Personality Consortium (GPC) for neuroticism, extraversion, and openness; Psychiatric Genomics Consortium (PGC) for ADHD, Alzheimer’s disease, alcohol misuse, anorexia nervosa, autism spectrum disorder, bipolar disorder, depressive symptoms, insomnia and schizophrenia; International Genomics of Alzheimer’s Project (IGAP) for Alzheimer’s disease, GWAS & Sequencing Consortium of Alcohol and Nicotine use (GSCAN) for age smoking initiation, cigarettes per day, drinks per week, ever smoker, and smoking cessation; Genetic Investigation of Anthropometric Traits (GIANT) Consortium for height and BMI; Cognitive Genomics (COGENT) Consortium for cognitive performance; CARDIoGRAMplusC4D Consortium for coronary artery disease; Global Lipids Genetics Consortium for blood lipids; International Consortium for Blood Pressure for blood pressure, Global Biobank Meta-analysis Initiative for asthma and COPD, Breast Cancer Association Consortium for breast cancer, The International Inflammatory Bowel Disease Genetics Consortium for inflammatory bowel disease, International Headache Genetics Consortium for migraine, The Prostate Cancer Association Group to Investigate Cancer Associated Alterations in the Genome (PRACTICAL) consortium for prostate cancer, and DIAMANTE (DIAbetes Meta-ANalysis of Trans-Ethnic association studies) and DIAbetes Genetics Replication And Meta-analysis (DIAGRAM) Consortia for type-II diabetes. We thank the research participants and employees of 23andMe for making this work possible. A full list of acknowledgements is provided in the Supplementary Note.

## Author contributions

D.J.B., A.O., P.T. and A.Y. designed and oversaw the study. A.O. supervised all analyses and led the writing of the manuscript. R.A. was responsible for the within-family validation analyses and making all figures except Figure 1. A.O. ran the UKB GWAS, conducted the quality control of GWAS summary statistics, ran the MTAG analyses, constructed the PGIs and PCs, and conducted the between-family PGI validation analyses. J.G. and A.Y. did the imputation of parental genotypes for UKB. A.T. developed the measurement error correction estimator for family-based analyses. D.J.B., P.T. and A.T developed the theoretical framework for the causal PGI study. M.H. designed and implemented the algorithm used to generate Figure 1. A.K., D.A.H., and the 23andMe Research Group conducted genome-wide association analyses for 23andMe. D.L., M.M., and D.C. contributed to study design. J.J. oversaw the data transfers and storage. All authors contributed to and critically reviewed the manuscript. R.A., D.J.B., D.C., A.O., and P.T. made especially major contributions to the writing and editing.

## 23andMe, Inc. Research Group

Stella Aslibekyan^5^, Adam Auton^5^, Robert K. Bell^5^, Katelyn Kukar Bond^5^, Zayn Cochinwala^5^, Sayantan Das^5^, Kahsaia de Brito^5^, Emily DelloRusso^5^, Chris Eijsbouts^5^, Sarah L. Elson^5^, Chris German^5^, Julie M. Granka^5^, Barry Hicks^5^, David A. Hinds^5^, Reza Jabal^5^, Aly Khan^5^, Matthew J. Kmiecik^5^, Alan Kwong^5^, Yanyu Liang^5^, Keng-Han Lin^5^, Matthew H. McIntyre^5^, Shubham Saini^5^, Anjali J. Shastri^5^, Jingchunzi Shi^5^, Suyash Shringarpure^5^, Qiaojuan Jane Su^5^, Vinh Tran^5^, Joyce Y. Tung^5^, Catherine H. Weldon^5^, Wanwan Xu^5^

## Estonian Biobank Research Team

Andres Metspalu^24^, Lili Milani^24^, Tõnu Esko^24^, Reedik Mägi^24^, Mari Nelis^24^ and Georgi Hudjashov^24^

## Figures

**Supplementary Figure 1.**
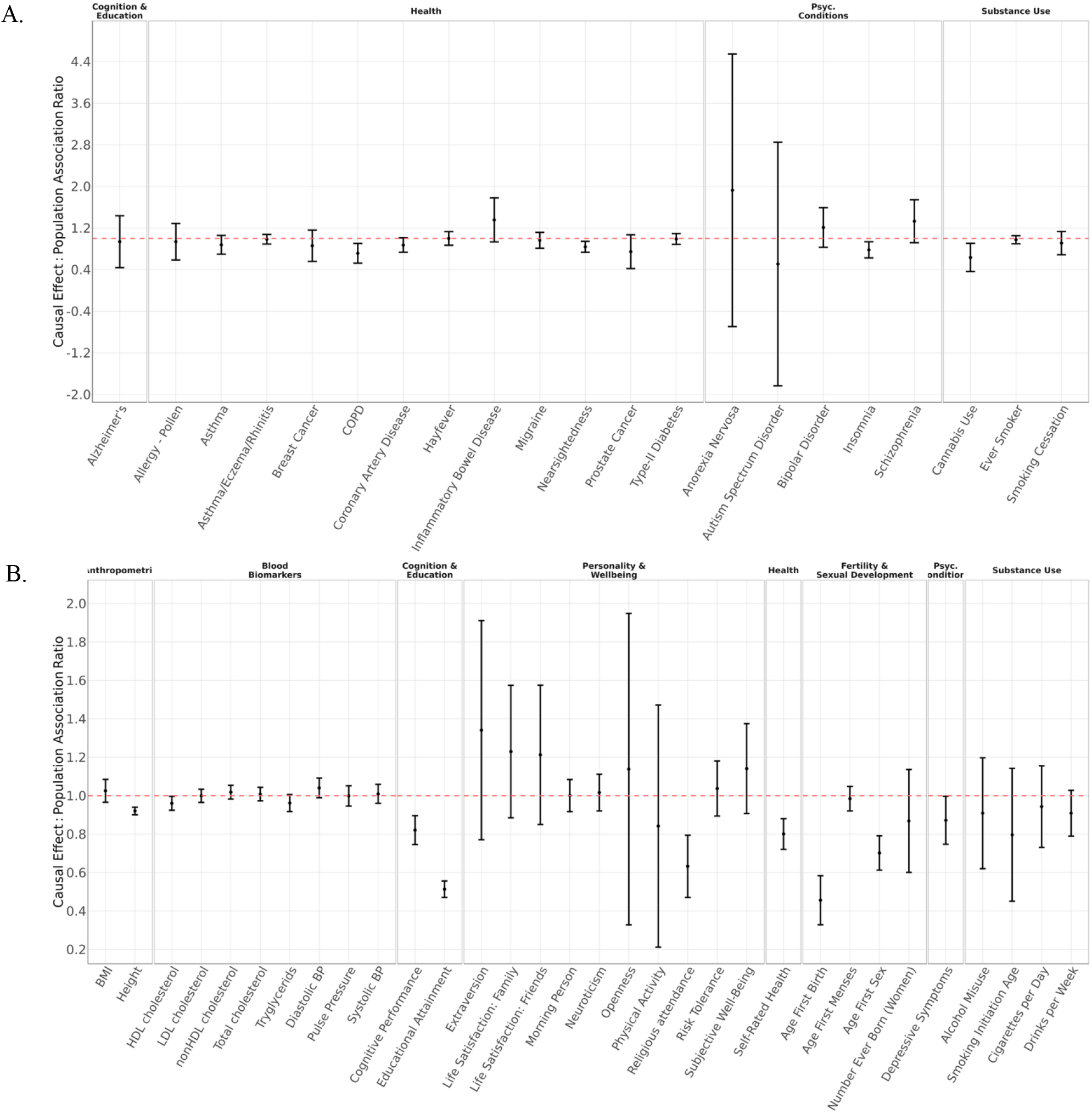
Ratio of causal effects of PGIs to their population associations meta-analyzed across UKB and WLS. *Notes:* Ratio of causal effects of PGIs to their population associations among binary (Panel A) and quantitative (Panel B) phenotypes. The causal effects were estimated in a sample of first-degree relatives, while the population associations were estimated in a sample of unrelated individuals in the third partition (UKB3). The ratios were calculated by dividing the direct effects by the population associations. Error bars represent 95% confidence intervals. Standard errors were computed using the delta method (Methods).

**Supplementary Figure 2.**
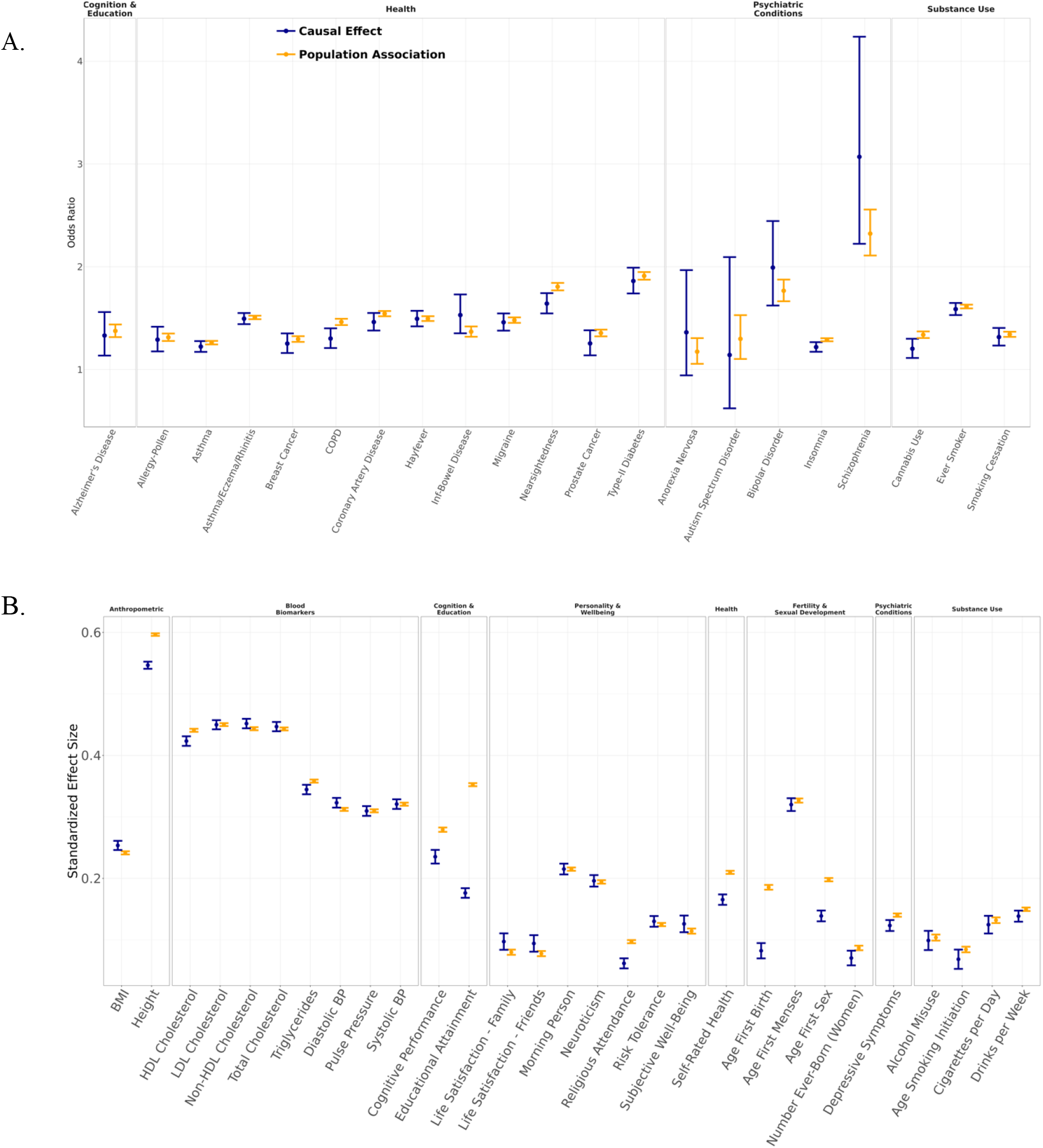
Causal effects of PGIs versus their population associations in UKB. *Notes*: Causal effects and population associations of PGIs in UKB. Causal effects were estimated in the sample of first-degree relatives, and population associations in a sample of unrelated individuals (third partition of UKB). For binary phenotypes, population associations were estimated using logistic regression and causal effects were estimated using a generalized mixed linear model with logistic link function in order to account for residual correlations between siblings (Panel A). For quantitative phenotypes, population associations were estimated using linear regression, and causal effects were obtained from a mixed linear model (Panel B). Error bars represent 95% confidence intervals.

**Supplementary Figure 3.**
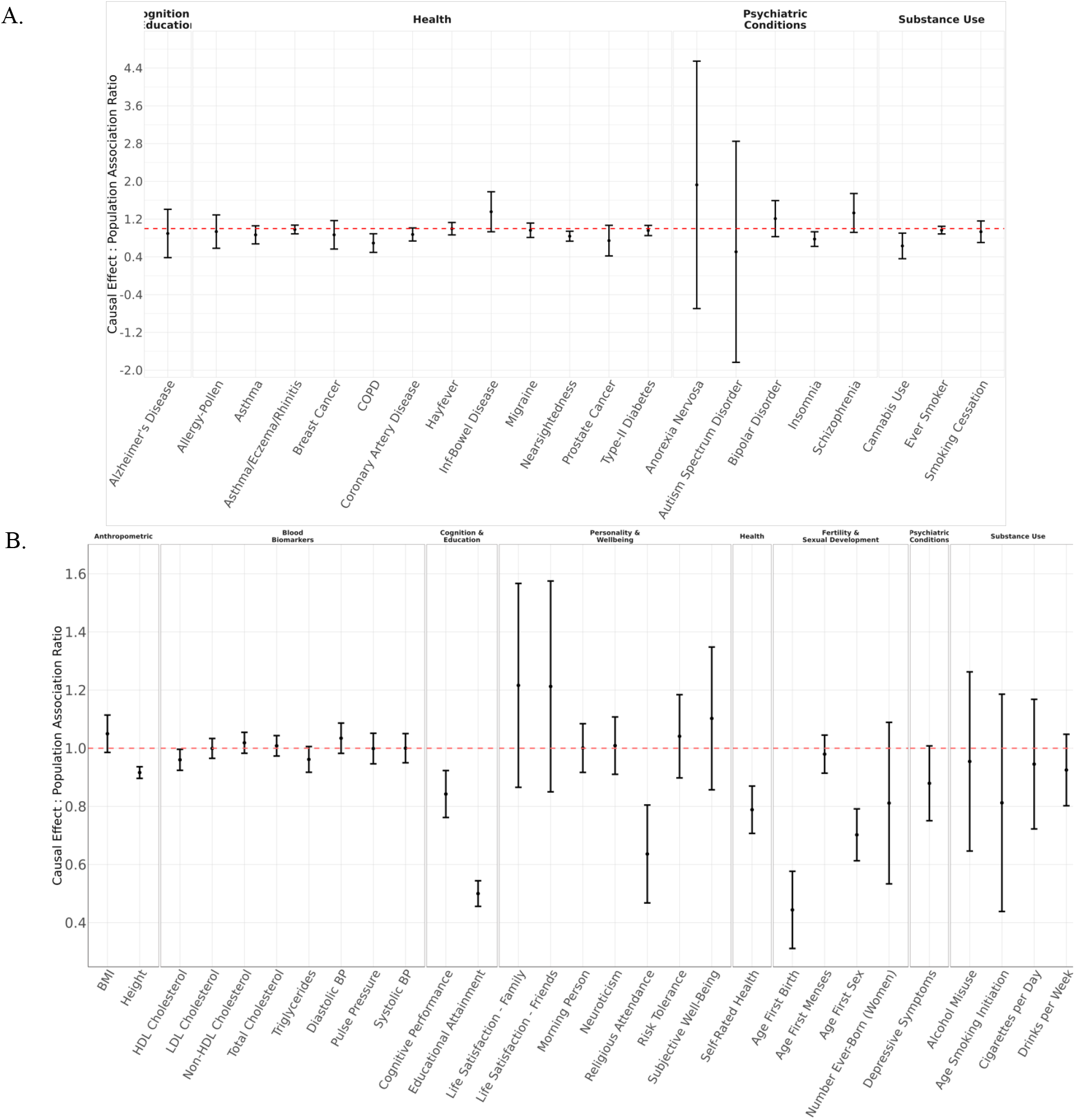
Ratio of causal effects of PGIs to their population associations in UKB. *Notes:* Ratio of causal effects of PGIs to their population associations among binary (Panel A) and quantitative (Panel B) phenotypes. The causal effects were estimated in a sample of first-degree relatives, while the population associations were estimated in a sample of unrelated individuals in the third partition (UKB3). The ratios were calculated by dividing the direct effects by the population associations. Error bars represent 95% confidence intervals. Standard errors were computed using the delta method (Methods).

**Supplementary Figure 4.**
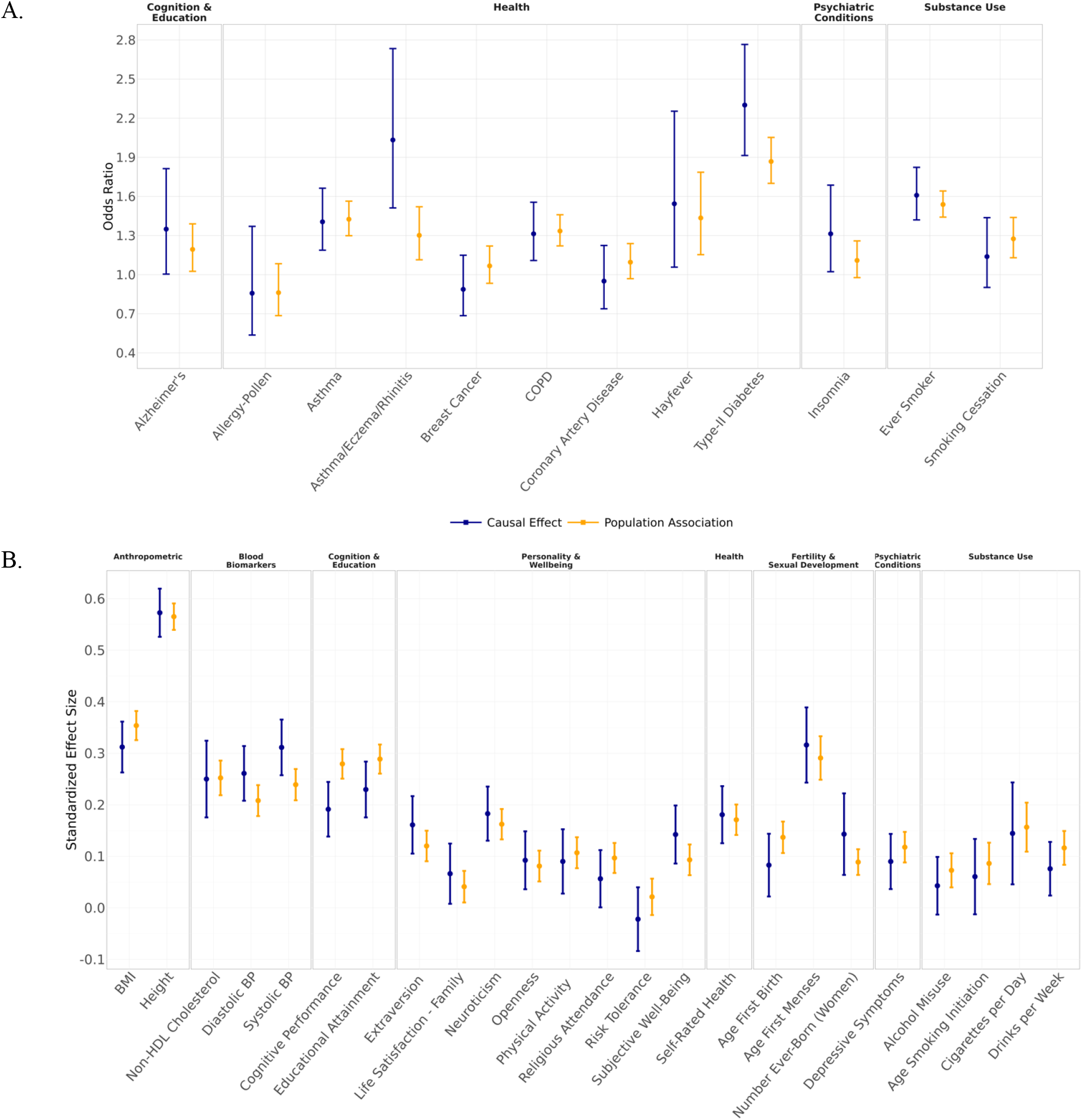
Causal effects of PGIs versus their population associations in WLS. *Notes:* Causal effects of PGIs versus their population associations in the WLS cohort. Causal effects were estimated using first-degree relatives (those with sibling genotype data), and population associations were estimated in the sample of unrelated individuals. For binary phenotypes, population associations were estimated using logistic regression and causal effects were estimated using a generalized mixed linear model with logistic link function in order to account for residual correlations between siblings (Panel A). For quantitative phenotypes, population associations were estimated using linear regression, and causal effects were obtained from a mixed linear model (Panel B). Error bars represent 95% confidence intervals.

**Supplementary Figure 5.**
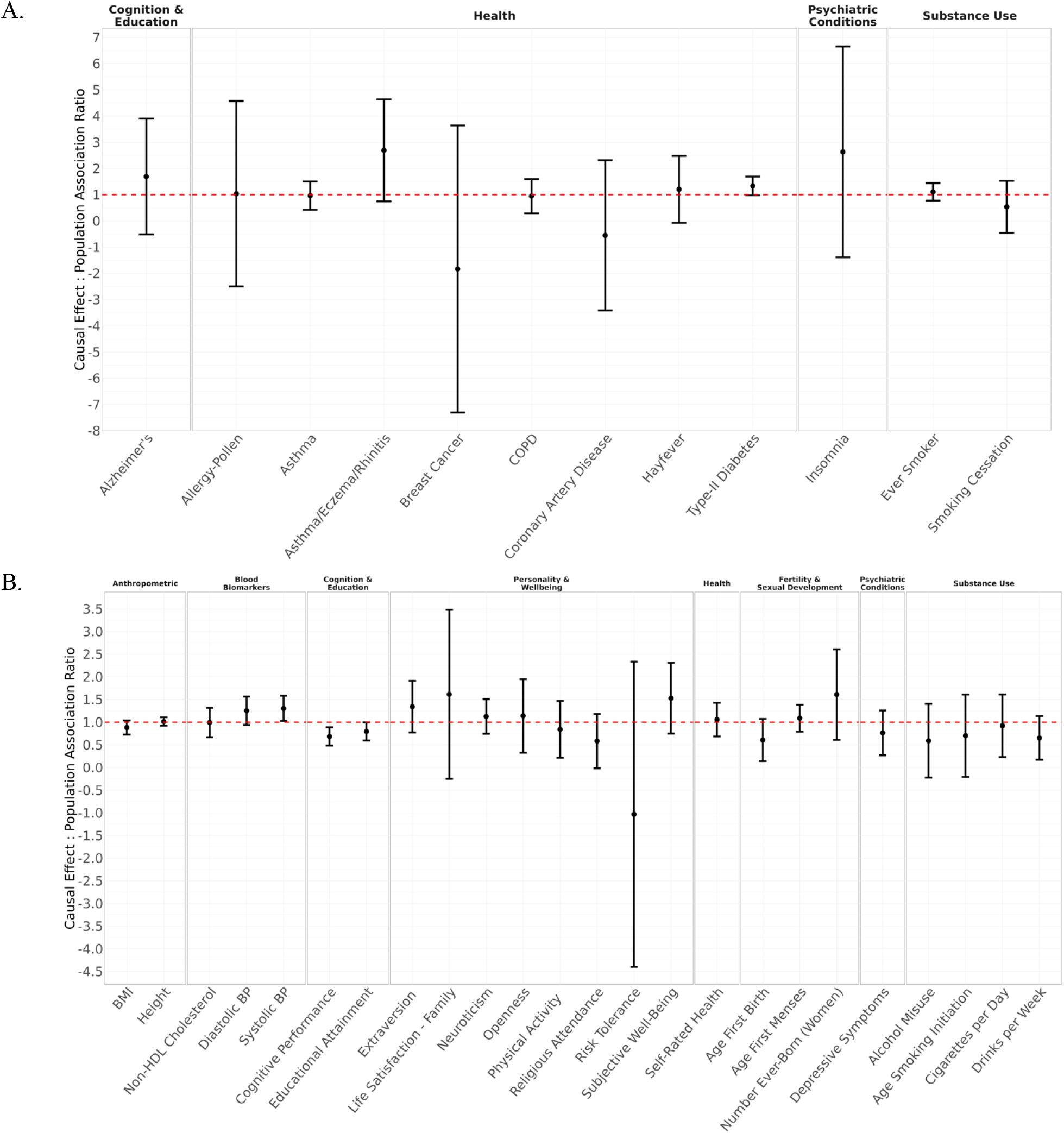
Ratio of causal effects of PGIs to their population associations in WLS. *Notes:* Ratio of causal effect to population association of PGIs among binary (Panel A) and quantitative (Panel B) phenotypes. Causal effects were estimated using first-degree relatives (those with sibling genotype data), and population associations were estimated in the sample of unrelated individuals. Ratios were obtained by dividing the estimates of causal effects by those of population association effects. Error bars represent 95% confidence intervals. Standard errors were computed using the delta method (Methods).

**Supplementary Figure 6.**
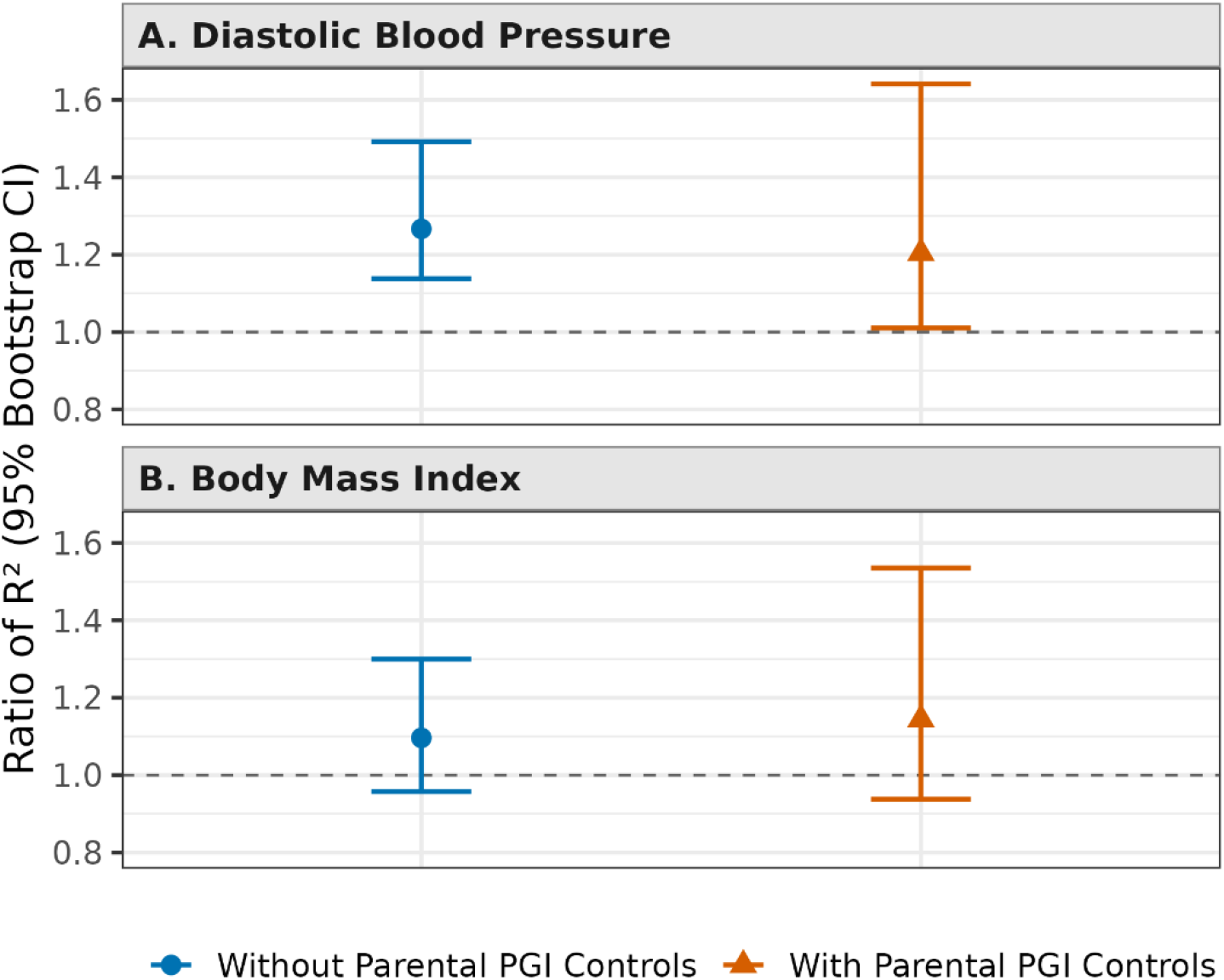
Ratios of incremental *R*^2^ between subgroups, with and without parental PGI controls. *Notes*: Ratio of PGI predictive accuracy (incremental *R*^2^) in subgroups of UKB for two traits, comparing models without parental PGI controls (blue circles) to models with parental PGI controls (orange triangles). Panel A shows the female / male incremental *R*^2^ ratio for diastolic blood pressure; Panel B shows the younger / older incremental *R*^2^ ratio for body mass index. Error bars are 95% confidence intervals obtained by the percentile method from 1,000 non-parametric bootstrap resamples of families within each subgroup. The dashed horizontal line at 1 indicates equal predictive accuracy across subgroups. Full model and bootstrap details are provided in the Methods.

## Footnotes

a The first release included PGIs based on multi-trait analyses of genetically correlated phenotypes. For some phenotypes, only the multi-trait (and not the single-trait) analyses yielded PGIs with expected predictive power greater than our inclusion cutoff of 1%. In the current release, we do not include multi-trait analyses, and therefore some of the 47 first-release phenotypes are not included among the current 61 phenotypes. These are: Number ever born (men), Age Voice Deepened, Allergy - Cat, Allergy - Dust, Cognitive Empathy, Delay Discounting, Life satisfaction – Finance, and Life Satisfaction – Work.

b To the extent that LD score regression^36^ successfully adjusts for population stratification, Corr(***x***_*i*_***γ, x***_*i*_ ***μ***) estimated using LDSC can also approach 1 when population stratification is the only source of confounding in ***µ***.

c Allergy - Cat, Allergy - Dust, Alzheimer’s Disease, Anorexia Nervosa, Attention Deficit and Hyperactivity Disorder, and Autism Spectrum Disorder.

d As in Becker et al., we recommend that researchers only do PGI analyses on sets of individuals within a partition. If researchers choose to do analyses with individuals across different partitions, their standard errors may be poorly calibrated; see Becker et al. (Methods, section “Using the UK Biobank split-sample PGI”) for discussion.

e Specifically, the following versions (see Supplementary Table 11): BL_CHOL3 - BL_CHOL5, BL_HDL3 - BL_HDL5, BL_LDL3 - BL_LDL5, BL_nonHDL2 - BL_nonHDL5, BMI1 - BMI5, EA1 - EA10, EA14, EA15, HEIGHT1 - HEIGHT6.

f EVERSMOKE1, EVERSMOKE4-6, CPD1, and CPD4 - CPD6 (see Supplementary Table 11).

