## Supplementary Note for "An Updated Polygenic Index Repository: Expanded Phenotypes, New Cohorts, and Improved Causal Inference"

May 9, 2025

#### Contents

**1 Data Access Procedures** **2**

**2 Dataset Profiles** **5**

**3 Dataset-Specific Acknowledgments** **7**

**4 Dataset Authorship Contributions** **13**

### 1 Data Access Procedures

#### 1958BC (NCDS)

Access to the polygenic indexes used in this manuscript, the underlying genetic data and all phenotypic data can be obtained free of charge by application to the Centre for Longitudinal Studies (CLS).(https://cls.ucl.ac.uk/data-access-training/data-access/accessing-data-directly-from-cls/). Please see the CLS Genomics GitHub repository (https://cls-genetics.github.io/) for more information.

#### 1970BC (BCS70)

Access to the polygenic indexes used in this manuscript, the underlying genetic data and all phenotypic data can be obtained free of charge by application to the Centre for Longitudinal Studies (CLS).(https://cls.ucl.ac.uk/data-access-training/data-access/accessing-data-directly-from-cls/). Please see the CLS Genomics GitHub repository (https://cls-genetics.github.io/) for more information.

#### 2001BC (MCS)

Access to the polygenic indexes used in this manuscript, the underlying genetic data and all phenotypic data can be obtained free of charge by application to the Centre for Longitudinal Studies (CLS).(https://cls.ucl.ac.uk/data-access-training/data-access/accessing-data-directly-from-cls/). Please see the CLS Genomics GitHub repository (https://cls-genetics.github.io/) for more information.

#### 23andMe, Inc.

Upon publication of this paper, investigators at non-profit institutions can obtain access to the genome-wide summary statistics from 23andMe used in this paper by completing the 23andMe Publication Dataset Access Request Form. The information provided on this form will be used to generate a Statement of Work (SOW) that will allow 23andMe to transfer data for use in the described research project. The SOW and a Data Transfer Agreement will need to be signed by the institution and 23andMe before data can be shared. The form, as well as additional information and requirements, are available at https://research.23andme.com/dataset-access/#how-to.

#### Add Health

Access to the polygenic indexes and full phenotype data in Add Health is publicly available via a restricted data use contract with the University of North Carolina at Chapel Hill. Obtain access by visiting the CPC Data Portal at data.cpc.unc.edu/projects/2/view or see the Add Health contracts page at www.cpc.unc.edu/projects/addhealth/contracts. Add Health genotype data can be accessed via the database of Genotypes and Phenotypes (dbGaP, www.ncbi.nlm.nih.gov/gap, accession number phs001367.v1.p1).

#### ALSPAC

The ALSPAC data management plan describes in detail the policy regarding data sharing, which is through a system of managed open access. Full instructions for applying for data access can be found on the study website (https://www.bristol.ac.uk/alspac/researchers/access/) and detailed in the ALSPAC access policy (https://www.bristol.ac.uk/media-library/sites/alspac/documents/researchers/data-access/ALSPAC_Access_Policy.pdf).

#### Dunedin

The datasets reported in the current article are available on request by qualified scientists. Requests require a concept paper describing the purpose of data access, ethical approval at the applicant's university, and provision for secure data access. We offer secure access on the Duke, Otago and King's

College campuses. All data analysis scripts and results files are available for review. For more information, see moffittcaspi.trinity.duke.edu/research-topics/dunedin.

#### ELSA

Polygenic indexes and genotype data are publicly available and are available here: https://www.elsa-project.ac.uk/genetics. Phenotype and other publicly available data can be downloaded from the UK Data Service: https://beta.ukdataservice.ac.uk/datacatalogue/studies/study?id=5050. Use is limited to non-profit research use only. For more information regarding the data please contact.

#### E-Risk

The datasets reported in the current article are available on request by qualified scientists. Requests require a concept paper describing the purpose of data access, ethical approval at the applicant's university, and provision for secure data access. We offer secure access on the Duke and King's College campuses. All data analysis scripts and results files are available for review. For more information, see moffittcaspi.trinity.duke.edu/research-topics/erisk.

#### EstBB

PGIs can be accessed for research and development purposes in accordance with the Estonian Human Genome Research Act (https://www.riigiteataja.ee/en/eli/ee/531102013003/consolide/current). To access data, the research proposal must be approved by the Scientific Advisory Committee of the Estonian Biobank as well as by the Estonian Committee on Bioethics and Human Research. Access to samples requires the same approval process and an additional approval from the Senate of the University of Tartu. For more details on data access and relevant documents, please see https://genomics.ut.ee/en/content/estonian-biobank#dataaccess.

#### FINBB

Researchers can apply for the PGIs in Finnish biobanks via the Fingenious portal (https://site.fingenious.fi/en/) hosted by the Finnish Biobank Cooperative FINBB (https://finbb.fi/en/). All Finnish biobanks can provide access for research projects within the scope regulated by the Finnish Biobank Act, which is research utilizing the biobank samples or data for the purposes of promoting health, understanding the mechanisms of disease or developing products and treatment practices used in health and medical care.

## GS

Generation Scotland data is available to researchers on application to. Further details can be found at https://genscot.ed.ac.uk/for-researchers/access.

#### HRS

PGIs are publicly available and can be downloaded here: https://hrsdata.isr.umich.edu/data-products/ssgac-polygenic-index-pgi-repository. Phenotype and other publicly available data can be downloaded here: hrs.isr.umich.edu/data-products. Genotype data can be accessed via NIAGADS (www.niagads.org/). Use is limited to non-profit research use only.

#### MCTFR

Access to the MCTFR PGIs is available by contacting Matt McGue, who will provide access authorization. Access to MCTFR phenotypic data will require a research proposal the structure of which can be provided by Matt McGue. Please note that the MCTFR is a complex, longitudinal study with thousands of relevant phenotypes assessed at multiple points in time. An overview of the range of phenotypes and developmental periods can be found in Wilson et al. (2019). Use of phenotypic data requires an approved proposal that is approved by the MCTFR Principal Investigator Committee; access to the MCTFR PGIs does not require an approved proposal. Because of the complexities involved, developing a proposal typically involves multiple iterations with MCTFR staff and are dealt with on a case-by-case basis.

##### MIDUS

Researchers interested in using the MIDUS data can find all pertinent information regarding data access at the following link: https://midus.wisc.edu/?page_id=89

##### SOEP-G

The PGIs will be shared by DIW Berlin in a standard phenotype file that can be accessed via user agreements with DIW Berlin (https://www.diw.de/en/diw_01.c.601584.en/data_access.html, contact).

##### STR

Researchers interested in using STR data must obtain approval from the Swedish Ethical Review Authority and from the Expert Committee of the Swedish Twin Registry. Researchers using STR data are required to follow the terms of a number of clauses designed to ensure protection of privacy and compliance with relevant laws. For further information please visit https://ki.se/en/research/research-infrastructure-and-environments/core-facilities-for-research/swedish-twin-registry-core-facility.

##### TTP

Access to the polygenic indexes and phenotype data from the Texas Twin Project is available via a restricted data use contract with the University of Texas at Austin. Restricted data users must develop an IRB-approved research proposal and security plan that ensures secure use of the data to minimize deductive disclosure risks. To apply for restricted-use data, please.

##### TwinLife

Access to the phenotype data from the TwinLife study is available via GESIS – Leibniz Institute for the Social Sciences. Researchers can request access by visiting the GESIS data catalog at https://search.gesis.org/research_data/ZA6701. Detailed documentation of the TwinLife phenotype data is available at https://www.twin-life.de/documentation/. PGIs generated for the TwinLife study are not hosted via GESIS and require a separate Data Transfer Agreement. Qualified researchers interested in accessing these PGIs must contact the TwinLife Data Center via email at. Requests should include a brief description of the proposed research project and institutional information to initiate the process.

##### UKB

All bona fide researchers can apply to use the UK Biobank resource for health related research that is in the public interest. Researchers can register and apply for data access at https://www.ukbiobank.ac.uk/register-apply/. Prior to publication of this paper, we will return the Repository PGIs to the UKB in accordance with their "returning results" procedure: https://biobank.ndph.ox.ac.uk/showcase/exinfo.cgi?src=returning_results. UKB will subsequently make the PGIs available to researchers as "Derived data-fields."

##### WLS

In addition to phenotype data, the polygenic index data is publicly available. As of February 2019, researchers who wish to use these polygenic indexes should email a brief research proposal and a copy or link to their CV to. Given the need to preserve participant confidentiality, to access the complete genotyped data, researchers will additionally need to receive IRB approval from their home institution and enter into a Data Use Agreement between the researcher's home institution and the University of Wisconsin-Madison. For the most up-do-date instructions, see `www.ssc.wisc.edu/wlsresearch/documentation/GWAS/`.

#### 2 Dataset Profiles

##### 1958BC (NCDS)

Power, C. & Elliott, J. Cohort profile: 1958 British birth cohort (National Child Development Study). *Int. J. Epidemiol.* **35**, 34–41 (2006).

Shireby, G. *et al.* Data Resource Profile: Genomic Data in Multiple British Birth Cohorts (1946-2001)—Health, Social, and Environmental Data from Birth to Old Age. *medRxiv* 2024.11.06.24316761 (2024) doi:10.1101/2024.11.06.24316761

##### 1970BC (BCS70)

Sullivan, A., Brown, M., Hamer, M. & Ploubidis, G. B. Cohort Profile Update: The 1970 British Cohort Study (BCS70). *Int. J. Epidemiol.* **52**, e179–e186 (2023).

Shireby, G. *et al.* Data Resource Profile: Genomic Data in Multiple British Birth Cohorts (1946-2001)—Health, Social, and Environmental Data from Birth to Old Age. *medRxiv* 2024.11.06.24316761 (2024) doi:10.1101/2024.11.06.24316761

##### 2001BC (MCS)

Joshi, H. & Fitzsimons, E. The Millennium Cohort Study: the making of a multi-purpose resource for social science and policy. *Longit. Life Course Stud.* **7**, 409–430 (2016).

Shireby, G. *et al.* Data Resource Profile: Genomic Data in Multiple British Birth Cohorts (1946-2001)—Health, Social, and Environmental Data from Birth to Old Age. *medRxiv* 2024.11.06.24316761 (2024) doi:10.1101/2024.11.06.24316761

##### 23andMe, Inc

Eriksson, N. *et al.* Web-based, participant-driven studies yield novel genetic associations for common traits. *PLoS Genet.* **6**, e1000993 (2010).

##### Add Health

Harris, K. M. *et al.* Cohort Profile: The National Longitudinal Study of Adolescent to Adult Health (Add Health). *Int. J. Epidemiol.* **48**, 1415-1415k (2019).

##### ALSPAC

Present study uses data from the G01 mothers and G01 children samples1–3. Pregnant women resident in Avon, UK with expected dates of delivery between 1st April 1991 and 31st December 1992 were invited to take part in the study. 20,248 pregnancies have been identified as being eligible and the initial number of pregnancies enrolled was 14,541. Of the initial pregnancies, there was a total of 14,676 foetuses, resulting in 14,062 live births and 13,988 children who were alive at 1 year of age. When the oldest children were approximately 7 years of age, an attempt was made to bolster the initial sample with eligible cases who had failed to join the study originally. As a result, when considering variables collected from the age of seven onwards (and potentially abstracted from obstetric notes) there are data available for more than the 14,541 pregnancies mentioned above: The number of new pregnancies not in the initial sample (known as Phase I enrolment) that are currently represented in the released data and reflecting enrolment status at the age of 24 is 906, resulting in an additional 913 children being enrolled (456, 262 and 195 recruited during Phases II, III and IV respectively). The phases of enrolment are described in more detail in the cohort profile paper and its update (see footnote 5 below). The total sample size for analyses using any data collected after the age of seven is therefore 15,447 pregnancies, resulting in 15,658 foetuses. Of these 14,901 children were alive at 1 year of age. Of the original 14,541 initial pregnancies, 338 were from a woman who had already enrolled with a previous pregnancy, meaning 14,203 unique mothers were initially enrolled in the study. As a result of the additional phases of recruitment, a further 630 women who did not enrol originally have provided data since their child was 7 years of age. This provides a total of 14,833 unique women (G0 mothers) enrolled in ALSPAC as of September 2021. Please note that the study website contains details of all the data that is available through a fully searchable data dictionary and variable search tool (`http://www.bristol.ac.uk/alspac/researchers/our-data/`) Ethical approval for the study was obtained from the ALSPAC Ethics and Law Committee and the Local Research Ethics Committees. Consent for biological samples has been collected in accordance with the Human Tissue Act (2004).

Boyd, A. *et al.* Cohort Profile: the 'children of the 90s'--the index offspring of the Avon Longitudinal Study of Parents and Children. *Int. J. Epidemiol.* **42**, 111–127 (2013).

Fraser, A. *et al.* Cohort Profile: the Avon Longitudinal Study of Parents and Children: ALSPAC mothers cohort. *Int. J. Epidemiol.* **42**, 97–110 (2012).

Northstone, K. *et al.* The Avon Longitudinal Study of Parents and Children (ALSPAC): an update on the enrolled sample of index children in 2019. *Wellcome Open Res.* **4**, (2019).

#### Dunedin

Poulton, R., Moffitt, T. E. & Silva, P. A. The Dunedin Multidisciplinary Health and Development Study: overview of the first 40 years, with an eye to the future. *Soc. Psychiatry Psychiatr. Epidemiol.* **50**, 679–693 (2015).

#### ELSA

Steptoe, A., Breeze, E., Banks, J. & Nazroo, J. Cohort Profile: The English Longitudinal Study of Ageing. *Int. J. Epidemiol.* **42**, 1640–1648 (2013).

#### E-Risk

Moffitt, T. E. *et al.* Teen-aged mothers in contemporary Britain. *J. Child Psychol. Psychiatry* **43**, 727–742 (2002).

#### EstBB

Milani, L. *et al.* The Estonian Biobank's Journey from Biobanking to Personalized Medicine. *Nat. Commun.* 2025 161 **16**, 1–13 (2025).

#### FINBB

Kurki, M. I. *et al.* FinnGen provides genetic insights from a well-phenotyped isolated population. *Nat.* 2023 6137944 **613**, 508–518 (2023).

### GS

Milbourn, H. *et al.* Generation Scotland: an update on Scotland's longitudinal family health study. *BMJ Open* **14**, e084719 (2024).

##### HRS

Sonnega, A. *et al.* Cohort profile: The Health and Retirement Study (HRS). *Int. J. Epidemiol.* **43**, 576–585 (2014).

##### MCTFR

Wilson, S. *et al.* Minnesota Center for Twin and Family Research. *Twin Res. Hum. Genet.* **22**, 746–752 (2019).

##### MIDUS

Ryff, C. D. *et al.* Midlife in the United States (MIDUS 2), 2004-2006. (2021) doi:10.3886/ICPSR04652.v8

##### SOEP-G

Koellinger, P. D. *et al.* Cohort profile: Genetic data in the German Socio-Economic Panel Innovation Sample (SOEP-G). *PLoS One* **18**, e0294896 (2023).

##### STR

Zagai, U., Lichtenstein, P., Pedersen, N. L. & Magnusson, P. K. E. The Swedish Twin Registry: Content and Management as a Research Infrastructure. *Twin Res. Hum. Genet.* **22**, 672–680 (2019).

##### TTP

Harden, K. P., Tucker-Drob, E. M. & Tackett, J. L. The texas twin project. *Twin Res. Hum. Genet.* **16**, 385–390 (2013).

##### TwinLife

Rohm, T. *et al.* Data from the German TwinLife Study: Genetic and Social Origins of Educational Predictors, Processes, and Outcomes. *J. Open Psychol. Data* **11**, 1–15 (2023).

##### UKB

Sudlow, C. *et al.* UK Biobank: An Open Access Resource for Identifying the Causes of a Wide Range of Complex Diseases of Middle and Old Age. *PLoS Med.* **12**, e1001779 (2015).

##### WLS

Herd, P., Carr, D. & Roan, C. Cohort Profile: Wisconsin Longitudinal Study (WLS). *Int. J. Epidemiol.* **43**, 34–41 (2014).


#### 3 Dataset-Specific Acknowledgments

We gratefully acknowledge research participants from all cohorts.

#### 1958 BC (NCDS)

The Centre for Longitudinal Studies is funded by the Economic and Social Research Council (grant numbers ES/M001660/1 and ES/W013142/1).

#### 1970 BC (BCS70)

The Centre for Longitudinal Studies is funded by the Economic and Social Research Council (grant numbers ES/M001660/1 and ES/W013142/1).

#### 2001 BC (MCS)

The Centre for Longitudinal Studies is funded by the Economic and Social Research Council (grant numbers ES/M001660/1 and ES/W013142/1; the Millennium Cohort Study (MCS) is additionally supported by grant reference ES/W001179/1).

#### 23andMe, Inc.

We gratefully acknowledge the contributions of members of 23andMe's Research Team, whose names are listed below: Stella Aslibekyan, Adam Auton, Robert K. Bell, Katelyn Kukar Bond, Zayn Cochinwala, Sayantan Das, Kahsaia de Brito, Emily DelloRusso, Chris Eijsbouts, Sarah L. Elson, Chris German, Julie M. Granka, Barry Hicks, David A. Hinds, Reza Jabal, Aly Khan, Matthew J. Kmiecik, Alan Kwong, Yanyu Liang, Keng-Han Lin, Matthew H. McIntyre, Shubham Saini, Anjali J. Shastri, Jingchunzi Shi, Suyash Shringarpure, Qiaojuan Jane Su, Vinh Tran, Joyce Y. Tung, Catherine H. Weldon, Wanwan Xu.

#### Add Health

The National Longitudinal Study of Adolescent to Adult Health (Add Health) is supported by grant P01 HD031921 to Kathleen Mullan Harris from the Eunice Kennedy Shriver National Institute of Child Health and Human Development (NICHD), with cooperative funding from 23 other federal agencies and foundations. Add Health GWAS data were funded by NICHD grants to Harris (R01 HD073342) and to Harris, Boardman, and McQueen (R01 HD060726). For information about access to the data from this study, contact.

#### ALSPAC

We are extremely grateful to all the families who took part in this study, the midwives for their help in recruiting them, and the whole ALSPAC team, which includes interviewers, computer and laboratory technicians, clerical workers, research scientists, volunteers, managers, receptionists and nurses. The ALSPAC study is funded by The UK Medical Research Council and Wellcome (Grant ref: 217065/Z/19/Z) and the University of Bristol provide core support for ALSPAC. This publication is the work of the authors and Aysu Okbay will serve as guarantor for the contents of this paper. A comprehensive list of grants funding is available on the ALSPAC website (`http://www.bristol.ac.uk/alspac/external/documents/grant-acknowledgements.pdf`); This research was specifically funded by grant from Wellcome Trust (WT088806) for the genotyping of ALSPAC mothers. For the children sample, GWAS data was generated by Sample Logistics and Genotyping Facilities at Wellcome Sanger Institute and LabCorp (Laboratory Corporation of America) using support from 23andMe.

#### Dunedin

Dunedin Multidisciplinary Health and Development Study research is supported by National Institute on Aging grants R01AG032282, R01AG049789, UK Medical Research Council grant MR/P005918, the New Zealand Health Research Council and New Zealand Ministry of Business, Innovation, and Employment.

#### ELSA

The English Longitudinal Study of Ageing is jointly run by University College London, Institute for Fiscal Studies, University of Manchester and National Centre for Social Research. Genetic analyses have been carried out by UCL Genomics and funded by the Economic and Social Research Council (ES/K005774/1) and the National Institute on Aging (R01 AG017644). All GWAS data has been deposited in the European Genome-phenome Archive. For more information please refer to`www.elsa-project.ac.uk/genetics`, or contact.

#### E-Risk

The E-Risk study is funded by grant G1002190 from the UK Medical Research Council and grant HD077482 from the National Institute of Child Health and Development.

#### EstBB

The activities of the EstBB are regulated by the Human Genes Research Act, which was adopted in 2000 specifically for the operations of the EstBB. Individual level data analysis in the EstBB was carried out under ethical approval 1.1-12/325, 23.01.2023 granted by the Estonian Committee on Bioethics and Human Research (Estonian Ministry of Social Affairs), using data according to release application 6-7/GI/30407 from the Estonian Biobank, transferred temporarily under Data Transfer Agreement number 6-2/GI/12127.

#### FINBB

We thank all those who contributed samples and data for the FinnGen scientific project.The FinnGen project is funded by two grants from Business Finland (HUS 4685/31/2016 and UH 4386/31/2016) and the following industry partners: AbbVie, AstraZeneca UK, Biogen, Bristol Myers Squibb (and Celgene Corporation & Celgene International II), Genentech, Merck Sharp & Dohme LLC, a subsidiary of Merck & Co., Inc., Rahway, NJ, USA, Pfizer, GlaxoSmithKline Intellectual Property Development, Sanofi US Services, Maze Therapeutics, Janssen Biotech, Novartis, and Boehringer Ingelheim. The following biobanks are acknowledged for delivering samples to FinnGen: Auria Biobank (`https://www.auria.fi/biopankki/`), THL Biobank (`https://www.thl.fi/biobank`), Helsinki Biobank (`https://www.helsinginbiopankki.fi`), Biobank Borealis of Northern Finland (`https://www.ppshp.fi/Tutkimus-ja-opetus/Biopankki/Pages/Biobank-Borealis-briefly-in-English.aspx`), Finnish Clinical Biobank Tampere (`https://www.tays.fi/en-US/Research_and_development/Finnish_Clinical_Biobank_Tampere`), Biobank of Eastern Finland (`https://www.ita-suomenbiopankki.fi/en`), Central Finland Biobank (`https://www.ksshp.fi/fi-FI/Potilaalle/Biopankki`), Finnish Red Cross Blood Service Biobank (`www.veripalvelu.fi/verenluovutus/biopankkitoiminta`) and Terveystalo Biobank (`https://www.terveystalo.com/fi/Yritystietoa/Terveystalo-Biopankk Biopankki/`). All Finnish biobanks are members of the BBMRI.fi infrastructure (`https://www.bbmri.fi`). The FINBB (`https://finbb.fi/`) is the coordinator of BBMRI-ERIC operations in Finland. The Finnish biobank data can be accessed through the Fingenious services (`https://site.fingenious.fi/en/`) managed by FINBB.

## GS

Generation Scotland received core support from the Chief Scientist Office of the Scottish Government Health Directorates [CZD/16/6] and the Scottish Funding Council [HR03006] and is currently supported by the Wellcome Trust [216767/Z/19/Z]. Genotyping of the GS:SFHS samples was carried out by the Genetics Core Laboratory at the Edinburgh Clinical Research Facility, University of Edinburgh, Scotland and was funded by the Medical Research Council UK and the Wellcome Trust (Wellcome Trust Strategic Award "STratifying Resilience and Depression Longitudinally" (STRADL) Reference 104036/Z/14/Z).

#### MCTFR

This project was led by William G. Iacono, PhD. and Matt McGue, PhD (Co-Principal Investigators) at the University of Minnesota, Minneapolis, MN, USA. Co-investigators from the same institution included: Irene J. Elkins, Margaret A. Keyes, James J. Lee, Lisa N. Legrand, Stephen M. Malone, William S. Oetting, Michael B. Miller, Saonli Basu and Scott Vrieze. Funding support for this project was provided through NIDA (U01DA024417). Other support for sample ascertainment and data collection came from several grants: R37DA05147, R01AA09367, R01AA11886, R01DA13240, R01MH66140.

#### MIDUS

Since 1995 the MIDUS study has been funded by the following: John D. and Catherine T. MacArthur Foundation Research Network; National Institute on Aging (P01-AG020166); National institute on Aging (U19-AG051426). Biomarker data collection was further supported by the NIH National Center for Advancing Translational Sciences (NCATS) Clinical and Translational Science Award (CTSA) program as follows: UL1TR001409 (Georgetown); UL1TR001881 (UCLA); 1UL1RR025011 (UW).

#### SOEP-G

We are deeply indebted to all individuals who have agreed to participate in the German Socio-Economic Panel Innovation Survey. The data collection was supported by the German Research Foundation (Leibniz Prize to Ralph Hertwig), a European Research Council Consolidator Grant (647648 EdGe to Philipp Koellinger), the Jacobs Foundation (Elliot M. Tucker-Drob, K. Paige Harden, Daniel W. Belsky), National Institute of Health/National Institute of Child Health and Human Development grant R01HD092548 (K. Paige Harden), the NORFACE DIAL Grant 462-16-100 (Pietro Biroli), the University of Basel (Rui Mata), and the Canadian Institute for Advanced Research (Daniel W. Belsky) and the Max Planck Institute for Human Development (Gert Wagner). The funders had no role in study design, data collection, and analysis, decision to publish, or manuscript preparation.

#### STR

The Swedish Twin Registry (STR) is managed by Karolinska Institutet and receives additional funding through the Swedish Research Council under the grant no 2017-00641 and 2021-00180. Other funding for the project come from the Ragnar Söderberg Foundation (E9/11), the Swedish Research Council (421-2013-1061).

#### TTP

The Texas Twin Project is supported by grants R01HD083613 and R01HD092548 from NIH/NICHD and Jacobs Foundation Research Fellowships.

#### TwinLife

The TwinLife study and its molecular genetic extension projects were funded by the German Research Foundation (DFG) (Grant numbers: 220286500, 428902522, and 458609264). The funder had no role in the study design, data collection and analysis, the decision to publish, or the preparation of the manuscript. We thank Shirin Zare for performing the DNA extraction.

#### WLS

This research uses data from the Wisconsin Longitudinal Study (WLS) of the University of Wisconsin-Madison. Since 1991, the WLS has been supported principally by the National Institute on Aging (AG-9775, AG-21079, AG-033285, and AG-041868, R01 AG041868-01A1), with additional support from the Vilas Estate Trust, the National Science Foundation, the Spencer Foundation, and the Graduate School of the University of Wisconsin-Madison. Since 1992, data have been collected by the University of

Wisconsin Survey Center. The opinions expressed herein are those of the authors. A public use file of data from the Wisconsin Longitudinal Study is available from the Wisconsin Longitudinal Study, University of Wisconsin-Madison, 1180 Observatory Drive, Madison, Wisconsin 53706 and at www.ssc.wisc.edu/WLSresearch/data/.

### 4 Dataset Authorship Contributions

| Dataset | Author | Study design & mgmt. | Data collection | Genotyping | Genotype prep. | Data analysis | Writing |
|---|---|---|---|---|---|---|---|
| **23andMe** | Aaron Kleinman | | | | | X | |
| **23andMe** | 23andMe Research Team | | X | X | | X | X |
| **1958BC (NCDS)** | David Bann | X | | | X | | X |
| **1958BC (NCDS)** | Tim Morris | X | | | X | | X |
| **1958BC (NCDS)** | George B. Ploubidis | X | X | X | | | X |
| **1970BC (BCS70)** | David Bann | X | | | X | | X |
| **1970BC (BCS70)** | Tim Morris | X | | | X | | X |
| **1970BC (BCS70)** | George Ploubidis | X | X | X | | | X |
| **2001BC (MCS)** | David Bann | X | | | X | | X |
| **2001BC (MCS)** | Tim Morris | X | | | | | X |
| **2001BC (MCS)** | Emla Fitzsimons | X | X | X | | | |
| **Add Health** | Kathleen Mullan Harris | X | X | X | | | |
| **Dunedin** | Avshalom Caspi | X | X | | | | |
| **Dunedin** | David L. Corcoran | | | X | X | | |
| **Dunedin** | Terrie E. Moffitt | X | X | | | | |
| **Dunedin** | Karen Sugden | | | X | X | | |
| **Dunedin** | Benjamin S. Williams | | | X | X | | |
| **ELSA** | Andrew Steptoe | X | X | | | | |
| **ELSA** | Olesya Ajnakina | | | | X | | |
| **E-Risk** | Avshalom Caspi | X | X | | | | |
| **E-Risk** | David L. Corcoran | | | X | X | | |
| **E-Risk** | Terrie E. Moffitt | X | X | | | | |
| **E-Risk** | Karen Sugden | | | X | X | | |
| **E-Risk** | Benjamin S. Williams | | | X | X | | |
| **EstBB** | Uku Vainik | X | X | X | X | | |
| **EstBB** | Tõnu Esko | X | X | X | X | | |
| **EstBB** | Estonian Biobank Research Team | X | X | X | X | | |
| **GS** | Archie Campbell | X | X | X | | | |
| **GS** | Caroline Hayward | X | X | X | | | |
| **MCTFR** | William G. Iacono | X | X | | | | |
| **MCTFR** | Matt McGue | X | X | | | | |
| **MIDUS** | Robert F. Krueger | X | X | X | X | | |
| **MIDUS** | Anna R. Docherty | X | X | | | | |
| **MIDUS** | Andrey A. Shabalin | X | X | | | | |
| **SOEP-G** | Ralph Hertwig | X | X | X | | | |
| **SOEP-G** | Philipp Koellinger | X | X | X | X | | |
| **SOEP-G** | David Richter | X | X | X | | | |
| **SOEP-G** | Jan Goebel | X | X | X | | | |
| **STR** | Rafael Ahlskog | X | | | | | |
| **STR** | Patrik K.E. Magnusson | X | X | | | | |
| **STR** | Sven Oskarsson | | | | | | X |
| **TTP** | K. Paige Harden | X | | | | | X |
| **TTP** | Elliot M. Tucker-Drob | X | | | | | |
| **TwinLife** | Charlotte K. L. Pahnke | X | | | | | |
| **TwinLife** | Carlo Maj | X | X | X | X | | |
| **TwinLife** | Frank M. Spinath | X | | | | | |
| **WLS** | Pamela Herd | X | X | X | | | |
| **WLS** | Jeremy Freese | X | X | X | | | |
