## Supplementary material for "An Updated Polygenic Index Repository: Expanded Phenotypes, New Cohorts, and Improved Causal Inference": Frequently Asked Questions

**Frequently Asked Questions (FAQs)**

This document provides information about the Social Science Genetic Association Consortium's (SSGAC) Polygenic Index Repository (https://www.thessgac.org/pgi-repository):

This version of the FAQs will become available upon publication of:

Alemu *et al.* (2025). "An Updated Polygenic Index Repository: Expanded Phenotypes, New Cohorts, and Improved Causal Inference" *bioRxiv*.

The document was prepared by Daniel Benjamin, David Laibson, Michelle N. Meyer, and Patrick Turley. It draws from and builds on the FAQs for earlier SSGAC papers. It has the following sections:

1. **Background**
2. **Study design and results**
3. **Social and ethical implications of the study**
4. **Appendices**

For clarifications or additional questions, please contact Daniel Benjamin.

1.10. How do different polygenic indexes for the same phenotypes differ? How comparable are results across studies that use different polygenic indexes for the same phenotypes? 11

### 1. Background

#### 1.1. Who conducted this study? What are the group's overarching goals?

The authors of the study are researchers affiliated with the Social Science Genetic Association Consortium (SSGAC) as well as data providers (i.e., individuals who act as stewards for datasets and provide other researchers with access to these data for research purposes). Social science is "the study of people: as individuals, communities and societies; their behaviours and interactions with each other and with their built, technological and natural environments" (Academy of Social Sciences, n.d.). It includes disciplines such as economics, psychology, sociology, political science, anthropology, and demography. Geneticists call an individual's observable characteristics *phenotypes*. Examples of phenotypes include height, blood type, Alzheimer's disease, dietary choice, religiosity (one's level of commitment to religious beliefs and practices), and educational attainment (how long someone went to school). The SSGAC is a multi-institutional, international research group that aims to identify statistically robust associations between differences in DNA and differences in social-science-relevant phenotypes like those involving human behavior, preferences, personality, cognition, and health. Many of the phenotypes studied by the SSGAC are also of interest to health researchers and other researchers.

SSGAC was founded on the belief that studying the links between genetics and social-science-relevant phenotypes can have substantial positive impacts across many research fields. This includes research that aims to better understand the effects of different environments (e.g., research on policy interventions) and the effects of interactions between genes and environments. The potential benefits also span a wide range of research questions in the biomedical sciences, such as why and how educational attainment is linked to longevity and better overall health phenotypes.

The SSGAC was formed in 2011 to address a specific set of scientific challenges. There are millions of common genetic differences across people (called SNPs, see FAQ 1.2). All *together*, these differences can be meaningful (see FAQs 1.3 & 2.3). But most phenotypes are, at best, only extremely weakly associated with any one particular common genetic difference. These associations are so weak that they are impossible to measure using sample sizes that are typical in social science research (e.g., hundreds of participants). To identify specific genetic differences with such small effects, scientists must study hundreds of thousands, if not millions, of people (to separate weak signals from sampling noise). One promising strategy for doing this is for many investigators to pool their data into one large study. This approach has borne considerable fruit when used by medical geneticists interested in a range of medical conditions (Visscher et al., 2017). Most of these advances would not have been possible without large research collaborations between multiple research groups interested in similar questions. The SSGAC was formed in an attempt by social scientists to adopt this research model.

The SSGAC is organized as a working group of the Cohorts for Heart and Aging Research in Genomic Epidemiology (CHARGE), a successful medical consortium. (In genetics research, a "cohort" is a group of research participants with something in common, such as having been enrolled into a biobank by a particular research institution.) The SSGAC was founded by three social scientists—Daniel Benjamin (University of California – Los Angeles), David Cesarini (New York University), and Philipp Koellinger (Vrije Universiteit Amsterdam and DeSci Labs). Today, the steering committee of the SSGAC includes Daniel Benjamin, David Cesarini, Aysu Okbay (Amsterdam University Medical Center), Patrick Turley (University of Southern California), and Alexander Young (University of California – Los Angeles).

To conduct such research, the SSGAC implements genome-wide association studies (GWAS) of social-scientific phenotypes. For example, to conduct a GWAS of educational attainment (Okbay et al., 2022) every participating cohort calculates for its participants the association between educational attainment and a genetic difference at a single location in the genome. This statistical analysis is repeated for each place on the genome where there are common genetic differences. The cohort-level results do not contain individual participant data—just aggregated, group-level "summary statistics" about these within-cohort statistical associations. The SSGAC then combines these cohort results to produce the overall GWAS results. By using existing datasets and combining cohort results, we can study the genetics of large numbers of individuals (for example, ~3 million people(Okbay et al., 2022). TheClick or tap here to enter text. SSGAC publicly shares overall, aggregated results (subject to some Terms of Service; see FAQ 3.6) so that other scientists can reproduce, further replicate (each GWAS publication contains its own replication), and build on this work. These publicly available data have already catalyzed many research projects and analyses across the social and biomedical sciences. Among the most useful products of these GWASs for other research are the polygenic indexes (PGIs) that are based on GWAS associations. PGIs are variables that aggregate the predictive power of many SNPs for predicting the phenotype of the GWAS (see FAQ 1.3).

The Advisory Board for the SSGAC is composed of prominent researchers representing various disciplines: Shawneequa Callier (Genomics Law and Policy, George Washington University), David Laibson (Economics, Harvard University), Alicia Martin (Population Genetics, Massachusetts General Hospital and Harvard Medical School) Michelle Meyer (Bioethics and Law, Geisinger College of Health Sciences), Melinda Mills (Sociology, University of Oxford), Peter Visscher (Statistical Genetics, University of Queensland), and Loic Yengo (Quantitative Genetics, University of Queensland, Australia).

The SSGAC is committed to the principles of reproducibility, transparency, and responsible communication of results. In 2013, SSGAC pioneered the practice of developing FAQ documents (such as this one) to accompany its major research publications and the practice has now become common in the field of social science genomics research (The Hastings Center, n.d.). FAQ documents aim to communicate research results less tersely and technically than in scientific papers, as well as what can and cannot be concluded from the research findings more broadly, and how they should and should not be used. FAQ documents produced for SSGAC publications are available on the SSGAC website, as well as in the Hastings Center FAQ Repository (along with FAQs from several other research teams).

To date, SSGAC-affiliated papers have studied educational attainment, cognitive performance, subjective well-being, reproductive behavior, risk tolerance, and dietary intake. The SSGAC website contains a list of our major publications, which have been published in journals such as *Science*, *Nature*, *Nature Genetics*, *Proceedings of the National Academy of Sciences*, *Psychological Science*, and *Molecular Psychiatry*.

##### 1.2. What is a single-nucleotide polymorphism (SNP)? What is a DNA base pair? What is an allele?

We study the most common sources of genetic difference, or variation: single-nucleotide polymorphisms (SNPs). SNPs are sites in the genome where single DNA base pairs commonly differ across individuals. A DNA "base pair" is a pair of two complementary nitrogen-containing bases—nucleotides—that are linked together in a DNA molecule, forming the "rungs" of the DNA ladder. Each SNP usually has two different possible variants of the DNA base pair at that particular site in the genome. These variants are called alleles. Although there are tens of millions of sites where SNPs are located in the human genome, our work (like most genetic research today that aims to link variation in DNA to variation in disease or other phenotypes) investigates only SNPs that can be easily measured with a high level of accuracy. These days, we can easily and accurately measure millions of SNPs, which together capture most of the common genetic variation across people.

##### 1.3. What is a polygenic index (PGI)? Why this terminology?

A polygenic index (PGI) is an index composed of a large number of SNPs (FAQ 1.3) from across the genome. Each PGI is associated with a particular phenotype (for details, see FAQ 1.4). Because a PGI aggregates the information from many SNPs, it can "predict" (see FAQ 1.7) vastly more of the variation among individuals than any single SNP. (Note that even PGIs are not good predictors of phenotypes for one person; see FAQ 3.3.) Often, the PGIs with the most predictive power are those created using millions of SNPs. A SNP array is the currently standard way of measuring common genetic differences across individuals. A SNP array does not measure the entire genetic sequence of each individual, but it does measure most of the places on the genome where individuals differ.

Our terminology of *polygenic index* is becoming more common, but it is still not the most common terminology. The traditional terms include polygenic risk score (PRS) and polygenic score (PGS). The word *risk* makes little sense when the polygenic index is for a non-disease phenotype (such as height). The word *score* was intended to echo statistical nomenclature but can instead convey an unintended value judgment or valence (i.e., "a higher score must be better"). The word *index* is at least as accurate statistically and does not convey a value judgment (Becker et al., 2021); Box 1.

##### 1.4. How is a polygenic index constructed?

A polygenic index (PGI) is constructed in three steps. First, a genome-wide association study (GWAS) is conducted, looking at SNPs (see FAQ 1.2) measured across the entire human genome to see which of them are associated with higher or lower levels of some phenotype. For each of these millions of SNPs, the GWAS generates an "effect size" corresponding to the (typically miniscule) magnitude of the association between that SNP and the phenotype. (We use the term "effect size" because it is a common scientific shorthand for "magnitude of association," but we emphasize that use of the term is not intended to imply that the SNP, or PGI, *causes* the phenotype; see FAQ 1.6.)

Second, the effect sizes are used to determine the "weight" each SNP will get in the PGI. The simplest scheme is to weight each SNP by its effect size as estimated in the GWAS. This simple weighting scheme has one main problem: because SNPs tend to be correlated with nearby SNPs on the genome (a phenomenon called linkage disequilibrium), if one SNP is associated with the phenotype, nearby SNPs will also be associated with the phenotype. Statistical approaches to determining the weights for constructing a PGI address this problem. We use an approach called SBayesR (Lloyd-Jones et al., 2019). Using the results of a GWAS, SBayesR generates a weight for each SNP. These weights are not equal to the SNPs' effect sizes as estimated in the GWAS, mostly because the weights take into account each SNP's correlation with other SNPs. (Even though SBayesR addresses the issue of linkage disequilibrium, it does so only for the purpose of generating weights for optimal prediction. SBayesR will not necessarily assign more weight to the SNP whose association with the phenotype is responsible for nearby SNPs' associations with the phenotype. Thus, SBayesR is a tool to address the issue of linkage disequilibrium for the purpose of prediction—which is the purpose of a PGI—but not for the purpose of unbiased estimation of SNPs' effect sizes. See FAQ 1.6.)

Third, the set of weights for the SNPs are used in a formula for calculating a PGI for any particular individual. The formula is a weighted sum of alleles at each SNP (using the weights from the second step). The formula is used to calculate a numerical value of the PGI for each individual in some dataset (that was not included in the GWAS).

The sample used for the GWAS in the first step is the training sample for the PGI. The larger the GWAS sample size, the greater the predictive power of a PGI constructed in the third step. However, this predictive power of a PGI has a maximum for each phenotype that the PGI can approach as the sample size gets bigger, but can never exceed no matter how large the same size is.

#### 1.5. How might polygenic indexes be useful?

Compared to other research tools in human genetics, polygenic indexes (PGIs) are relatively new, and we are just beginning to see their potential. The idea of using GWAS results to create a PGI was initially proposed in 2007 (Wray et al., 2007), and the first PGI was created in 2009 in a GWAS of schizophrenia and bipolar disorder (Purcell et al., 2009). Since then, PGIs have become a significant part of research that builds on genetics in the medical and social sciences. For example, in the current paper (Alemu et al., 2025) we analyze presentations at the annual meeting of the Behavior Genetics Association. We report that the fraction of presentations that used PGIs increased from 0% in 2009 to 40% in 2024.

A PGI for a phenotype (e.g., a subjective well-being PGI) provides one measure of the genetic influence on that phenotype that can be used in research in a variety of ways. For example, PGIs have been used to:

- partially control for genetic influences in order to generate less noisy estimates of how changes in school policy influence health phenotypes (Davies et al., 2018);

- examine how the effect of school policy on health phenotypes depends in part on genetic influences (Barcellos et al., 2018);
- study why SNPs predict educational attainment – for example, it appears that some genetic effects on educational attainment operate through associations with cognitive function and traits such as self-control (Belsky et al., 2016), which in turn affect educational attainment;
- investigate how genetic influences on educational attainment differ across environmental contexts (Barcellos et al., 2018; Schmitz & Conley, 2017);
- investigate how genetic influences on BMI vary over the lifecycle (Khera et al., 2019);
- infer the degree of assortative mating in particular groups (Robinson et al., 2017; Yengo et al., 2018);
- trace recent migration patterns (Abdellaoui et al., 2019; Domingue et al., 2018);
- examine whether PGIs for disease risk are sufficiently predictive to be incorporated into clinical practice for preventative medicine (Khera et al., 2018);
- study the intergenerational transmission of wealth (Barth et al., 2022; S. Y. T. Lee & Seshadri, 2019);
- investigate how parental investment responds to children's abilities (Houmark et al., 2024); and
- develop new statistical tools that may advance our understanding of how parenting and other features of a child's rearing environment influence his or her developmental phenotypes (Koellinger & Harden, 2018; Kong et al., 2018).

Almost all of the uses of PGIs in research, including in the above list, have focused on statistical associations, rather than causal effects of genetic variants (see FAQ 1.6). In the past few years, a growing number of datasets with genotyped family members—such as sibling pairs, or trios of parents and their offspring—have become available. With such family data, it is possible to use PGIs to study causal effects of genetic variants (see FAQ 1.8). In our analysis of presentations at the annual meeting of the Behavior Genetics Association, we report that among the presentations that used PGIs, the fraction that used family data to conduct causal analyses increased from 0% in 2019 to 27% in 2024.

As discussed in FAQ 1.10, one goal of the Polygenic Index Repository is to facilitate further work using PGIs by making a much wider range of more predictive PGIs available to researchers. As discussed in FAQ 2.4, another goal is to facilitate causal analyses by making PGIs for individuals' parents available in family-based datasets.

#### 1.6. Why are most polygenic index studies not "causal"?

Polygenic indexes (PGIs) available today, including those we construct in this paper, should not be interpreted as a measure of causal mechanisms, for the reasons we explain below. Therefore, unless the study is designed in a specific way that gives it a causal interpretation (see FAQ 1.8), the results of a study should be interpreted as "predictive" rather than causal (see FAQ 1.7).

The genome-wide association studies (GWASs) used as the training data for the PGIs (see FAQ 1.4) identify SNPs that are associated with the phenotype, but an empirical correlation with a specific SNP need not imply that the SNP *causes* the phenotype, for a variety of reasons. First, SNPs are often highly correlated with other, nearby SNPs on the same chromosome. As a result, when one or more SNPs in a region causally influence a phenotype (in that particular environment), many non-causal SNPs in that region may also be identified as associated with the phenotype (in FAQ 1.4, see the parenthetical "Even though SBayesR…" for why SBayesR does not solve this problem for the purpose of identifying the causal SNP). In fact, the causal SNP may not have even been measured directly. For example, GWAS that focus on common SNPs would not be able to identify rare or structural types of genetic variation (e.g., deletions or insertions of an entire genetic region) that are causal, but they may identify SNPs that are correlated with these unobserved variants. For these and other reasons, PGIs are likely to be composed of a mix of causal and non-causal SNPs, and the weights used in the formula for constructing the PGI (see FAQ 1.4) should not be interpreted as estimates of the causal effects of the SNPs.

Second, at a particular SNP the frequency of different alleles might vary systematically across environments. If those environmental factors are not accounted for in the association analyses, some of the measured SNP associations with social-science phenotypes may be spurious. To use a well-known example often used to explain this idea (Lander & Schork, 1994), any genetic variants common in people of Asian ancestries will be associated statistically with more frequent than average chopstick use, but these variants would not *cause* greater chopstick use; rather, these genetic variants and the phenotype of chopstick use are both distributed unevenly among people with different ancestries. This is called the problem of "population stratification." The GWAS underlying the PGIs in the Repository employ standard strategies to try to minimize this problem, but the issues raised by population stratification cannot be ruled out entirely. As a result, the PGIs likely reflect population stratification to some extent. In the User Guide that accompanies the Polygenic Index Repository (reproduced in the Supplementary Methods of the paper), we discuss this problem in more detail and discuss strategies for addressing population stratification that is reflected in the PGIs.

Even in GWASs (such as those we rely on or conduct ourselves) that attempt to address and correct for heterogeneity in genetic ancestry, allele frequencies may nonetheless vary systematically with environmental factors even *within* a group of people of similar genetic ancestry. For example, a SNP that is associated with improved educational outcomes in the parental generation may have downstream effects on parental income and other factors known to influence children's educational outcomes (such as neighborhood characteristics). This same SNP is likely to be inherited by the children of these parents, creating a correlation between the presence of the SNP in a child's genome and the extent to which the child was reared in an environment with specific characteristics. A study of Icelandic families showed that a parental allele associated with higher educational attainment of the parent that is *not* passed on to the parent's offspring is still associated with the child's educational attainment, suggesting that GWAS results for educational attainment partly represent these intergenerational environmental pathways (Kong et al., 2018).

The second and third reasons above relate to confounds: reasons why at least part of the predictive power of a polygenic index is typically driven by factors *other than* the causal effects of genetic variants. As a very rough estimate, for social and behavioral phenotypes, no more than about one-third of the predictive power of a PGI (i.e., the percentage of the variance in the phenotype among individuals that the PGI explains) is explained by causal genetic effects taken together (Howe et al., 2022) i.e., the causal effects of SNPs included in the PGI together with the causal effects of genetic variants correlated with included SNPs. For instance, the most predictive PGI for educational attainment currently available explains about 12% of the variance between people, but only one-third of that – about 4%– is causal. These causal SNPs may be among the SNPs included in the PGI or may be physically close to, and therefore correlated with, SNPs that are included. In contrast, for anthropometric phenotypes such as height, it is possible that nearly all of the predictive power of a PGI is explained by causal SNPs in the PGI or correlated with those included SNPs.

Third, a SNP's effects on a phenotype may be indirect, so a SNP that may be "causal" in one environment may have a diminished effect or no effect at all in other environments. For example, variation in a particular SNP on chromosome 15 is associated with lung cancer (Amos et al., 2008; Hung et al., 2008; Thorgeirsson et al., 2008). From this observation alone we cannot conclude that variation in this SNP can cause lung cancer through some direct *biological* mechanism. In fact, it is likely that variation in this SNP, which is part of the nicotinic acetylcholine receptor gene cluster that affects nicotine metabolism, increases lung cancer risk through effects on smoking behavior. In a tobacco-free environment, it is plausible that this association with lung cancer would be substantially weaker and perhaps disappear altogether. Thus, even *if* we have credible evidence that a specific association is not spurious, it is entirely possible that the SNP in question influences the phenotype through channels that we, in common parlance, would label environmental (e.g., smoking). Nearly forty years ago, the sociologist Christopher Jencks criticized the widespread tendency to mistakenly treat environmental and genetic sources of variation as mutually exclusive (Turkheimer, 2000). As the example of smoking illustrates, it is often overly simplistic to assume that "genetic explanations of behavior are likely to be exclusively physical explanations while environmental explanations are likely to be social" (Jencks, 1980).

#### 1.7. In what sense does a polygenic index "predict" the phenotype of interest?

When we and other scientists say that polygenic indexes (and other variables, such as demographics or other environmental factors) "predict" certain phenotypes, our use of "predict" differs in several important ways from common parlance. First, we do not mean that the polygenic index (PGI) guarantees an outcome with 100% probability, or even with a high degree of likelihood. Rather, we mean that the PGI is, on average across people, statistically associated with a phenotype. In other words, on average, people with a higher numerical value of the PGI have a higher likelihood of the phenotype compared to people with a lower numerical value. A PGI is said to be statistically "predictive" of a phenotype even if the PGI has only a *weak* association with the phenotype—as is the case, for instance, with almost all of the PGIs in the Repository. In such cases, the PGI is only weakly predictive of the phenotype.

Second, in standard language, "prediction" usually refers to the future. In contrast, when scientists say that a PGI "predicts" a phenotype, they mean that they expect to see the association in *new data*. "New data" means data that haven't been analyzed yet—regardless of whether those data will be collected in the future or have already been collected. In other words, in social science, it makes perfect sense to ask how well a PGI predicts phenotypes that have already occurred, like how many years of education were attained by older adults.

Finally, in standard language, a "prediction" is often an unconditional guess about what will happen. Instead of meaning it unconditionally, scientists mean that they expect to see an association in new data under certain conditions, for example, that the environment for the new data is the same as the environment in which the GWAS that underlies the PGI (see FAQ 1.4) was conducted. In the example given in FAQ 1.6, in which a SNP is associated with lung cancer due to an effect on smoking, we would *not* expect the SNP to be as strongly predictive of lung cancer, or predictive at all, in an environment where tobacco-based products are hard to obtain or absent entirely.

#### 1.8. What is a "causal polygenic index study," and how does it differ from a (usual) population-based polygenic index study?

A "causal polygenic index (PGI) study" refers to a study that relates variation in a phenotype to variation in a PGI *within families*. (There are many caveats to referring to these PGI studies as "causal," as we discuss in detail below.) As a concrete example, (Okbay et al., 2022) examined how a PGI for educational attainment is causally related to a wide range of phenotypes. In one analysis, using ~3,500 individuals with both parents genotyped, they looked at how an individual's PGI is related to the individual's self-rated overall health, controlling for the average PGI of the individual's parents. This kind of study is causal because genetic inheritance from parents to offspring is a natural experiment. *On average*, every individual's PGI is halfway in between their mother's PGI and their father's PGI. But because it is random (like a coin flip) which of their parent's two alleles an individual inherits, some individuals end up with a higher PGI and others with a lower PGI than the parental average. It is therefore as if an experimenter randomly assigned some people to have higher PGIs than their parental average, and others to have lower PGIs. By examining how this random component of an individual's PGI relates to their overall health, (Okbay et al., 2022) could isolate the causal effects of genetic variants. This is in contrast to a usual, *population-based* PGI study, which in this example would merely examine the association between an individual's PGI and their overall health (without controlling for the average PGI of the individual's parents). In a population-based PGI study, researchers cannot conclude to what extent genetic variants, as opposed to environments that are correlated with those genetic variants, are responsible for any observed association between an individual's PGI and their overall health.

The correct interpretation of a causal PGI study is nuanced—and, indeed, one goal of our paper that introduces version 2 of the Repository (Alemu et al., 2025) is to help clarify the correct interpretation. A causal PGI study estimates a complicated weighted average of causal effects of genetic variants, including genetic variants that are not themselves part of the PGI but which are correlated with SNPs included in the PGI (i.e., in LD with those SNPs). Three natural ways that "causal PGI study" might be interpreted are incorrect, as we now explain.

First, as already noted, it is incorrect to think that a "causal PGI study" captures only the causal effects of genetic variants that are included in the PGI. It also captures causal effects of genetic variants that are not themselves in the PGI if they are correlated with the SNPs included in the PGI.

Second, in most cases it is incorrect to think of a PGI itself as having a causal effect. This is because a PGI is, by construction, an index, meaning that it is a weighted sum of genotypes from many genetic variants (typically, omitting all interaction effects). To give an extreme but illustrative example, suppose that a PGI was composed of two SNPs. Suppose an increase of one minor allele for either SNP would increase the PGI by one standard deviation (a statistical unit), but the two SNPs have different causal effects on the phenotype. (Hypothetically, suppose one allele affects memory while the other affects preference for learning, and the phenotype of educational attainment is only affected when both memory and preference for learning change, because they interact with each other.) Then there is no single answer to the question, *What is the effect of increasing the PGI by one standard deviation?*, because the answer depends on which SNP is changed. (For instance, if the one standard deviation increase is due only to the memory-affecting allele, or only to the preference for learning allele, then there would be no effect on educational attainment.) In a causal PGI study, the correct way to interpret the effect of “increasing the PGI by one standard deviation” is that it corresponds to a particular weighted sum of the causal effects of the genetic variants captured by the index. Recent research has characterized what these weights are (Benjamin et al., 2024; Veller et al., 2024). Put another way, it is not technically meaningful to talk about “the effect of a PGI” (Veller et al., 2024); when researchers use that language, it is—or should be—shorthand for referring to a weighted sum of the causal effects of genetic variants. (This kind of shorthand is standard in the economics literature, where analogous situations often arise. For example, it is not technically meaningful to talk about “the effect of an additional year of schooling” because it depends on how much schooling a person already has, which school it is, what classes are taken, who the teachers are, etc. However, economists use this language as shorthand to refer to a weighted average of the causal effects across these variables.)

##### 1.9. What polygenic indexes were available to researchers prior to the Polygenic Index Repository?

Prior to the Polygenic Index Repository, only a few datasets had constructed polygenic indexes (PGIs) that researchers could download and use. Notable examples of data providers that did make PGIs directly available to researchers—all of which recognized early on the value of doing so—are the Health and Retirement Study, the Wisconsin Longitudinal Study, and the National Longitudinal Adolescent to Adult Health Study. The UK Biobank does not construct PGIs for its users, but it provides a mechanism by which researchers who use the data and construct PGIs can “return” them to the UK Biobank for use by other researchers. Through this mechanism, PGIs constructed from several GWASs have been made available for researchers to download from the UK Biobank.

To study PGIs in other datasets or for other phenotypes, prior to the Polygenic Index Repository, researchers would need to construct the PGIs themselves, following the steps described in FAQ 1.4. For the first step, most researchers would need to rely on publicly available GWAS results, which include less data and are therefore less predictive than some PGIs in published work that rely on non-public GWAS results (see FAQ 2.3). To make it easier for researchers to construct PGIs themselves, the Polygenic Score Catalog (Lambert et al., 2021) collects together weights for a range of PGIs (also based on publicly available GWAS results).

As we discuss in more detail in FAQ 2.1, for the Polygenic Index Repository, we constructed a large number of PGIs in each of a growing number of datasets (including the four mentioned above)—initially 11 datasets, and now 20—and have made the PGIs directly available for researchers to download. The PGIs are often based on more data than is publicly available, and the PGIs are constructed according to a uniform methodology across both phenotypes and datasets. For examples of Repository PGIs that were previously not available at all or that were less accurate (i.e., predictive), see FAQ 2.3.

##### 1.10. How do different polygenic indexes for the same phenotypes differ? How comparable are results across studies that use different polygenic indexes for the same phenotypes?

There are several reasons why polygenic indexes (PGIs) for the same phenotype can differ from each other. As described in FAQ 1.4, there are three steps to creating a PGI, and differences can arise at each of these steps. For example, in the first step, researchers could base the PGI on different GWAS studies of the same phenotype. Different GWAS studies may be based on samples who live under different environmental conditions, may have different measures of the phenotype, and/or may have measured different SNPs. As another example, in the second step, researchers could use a different method of determining polygenic-index weights from the results of a GWAS. For these and other reasons, it has been common for different studies to use different PGIs, even when the PGIs are for the same phenotype and are being studied in the same dataset.

The results are typically difficult to compare across such studies for three main reasons:

1. If the PGIs are constructed using different methods, then even though they are both measuring genetic influences on the same phenotype, the precise definition of these "genetic influences" may differ (see FAQs 3.1 and 3.2).
2. The units for measuring the strength of associations between the PGI and other variables generally differ across studies. Researchers usually report results in terms of standard deviations (a statistical unit) of the PGI, but if the PGI in one study is a more powerful predictor than that in the other study, then one standard deviation of one PGI means something different than one standard deviation of the other.
3. If one of the PGIs is a more powerful predictor than the other, then they differ in their signal-to-noise ratio for capturing genetic influences on the phenotype. Whenever an explanatory variable is measured with noise, results based on that variable will be distorted, sometimes in unanticipated ways. Since the signal-to-noise ratio differs across the PGIs, results based on them are distorted differentially, further making the results difficult to compare.

#### 1.11. Why create the Polygenic Index Repository?

In brief, from when it was initially introduced, the Polygenic Index Repository has had three main goals: (i) to make polygenic indexes (PGIs) for a large number of phenotypes more accessible to a wider range of researchers from many fields and disciplines, including early career researchers, researchers without access to the data and/or training required to create the most state-of-the-art PGIs, and researchers who wish to probe the limitations of PGIs; (ii) to increase the use of PGIs that are more accurate (i.e., predictive) than PGIs researchers could construct from publicly available GWAS results and which therefore have greater statistical power (and are less prone to false positives); and (iii) to facilitate the comparability of results across studies that use these PGIs. The second release of the Polygenic Index Repository added an additional goal: (iv) in family-based datasets, to make available PGIs for individuals' parents, in order to facilitate causal analyses.

In more detail, the Polygenic Index Repository addresses several practical obstacles that researchers interested in using PGIs must often confront, including:

1. Constructing a PGI from genotype data requires special expertise. Even for researchers with that expertise, it can be a time-consuming process.
2. It is generally desirable to generate polygenic-index weights from the GWAS with the largest sample size because the predictive accuracy of a PGI is expected to be largest in that case. However, there are administrative hurdles for accessing some GWAS results, such as those from 23andMe. In practice, researchers often end up constructing PGIs using only publicly available GWAS results. Such PGIs tend to have less predictive power.

3. Publicly available GWAS results are sometimes based on a sample that includes the dataset (or close relatives of dataset members) in which the researcher wants to analyze the PGI. Such "sample overlap" spuriously inflates the predictive power of the PGI, which can lead to highly misleading results.
4. Because different researchers construct PGIs in different ways, it is hard to compare and interpret results from different studies (see FAQ 1.9).

As we explain in the paper that introduced the Polygenic Index (Becker et al., 2021):

> We overcome no. 1 by constructing the PGIs ourselves and releasing them to the data providers, who in turn will make them available to researchers. This simultaneously addresses no. 2 because we use all the data available to us that may not be easily available to other researchers or to the data providers, including genome-wide summary statistics from 23andMe. Using these genome-wide summary statistics from 23andMe is what primarily distinguishes our Repository from existing efforts by data providers to construct PGIs and make them available…It also distinguishes our Repository from efforts to make publicly available PGI weights directly available for download (although we also do that, for weights constructed without 23andMe data). To deal with no. 3, for each phenotype and each dataset, we construct a PGI from GWAS summary statistics that excludes that dataset. We overcome no. 4 by using a uniform methodology across the phenotype.

In addition to providing PGIs constructed using a uniform methodology (which deals with problem #1 listed in FAQ 1.9), we aim to improve comparability of results based on PGIs in another way (which deals with problems #2 and #3 listed in FAQ 1.9): we derive a "measurement-error-corrected estimator" and provide software for calculating it. This estimator deals with the fact that PGIs can differ from each other in their signal-to-noise ratios. It estimates what the results of an analysis would be if the PGI had no noise. It thereby avoids the distortions in results that arise from having a noisy measure. Because it puts results about the PGI in the units of the "noiseless" PGI, the results from PGIs with different signal-to-noise ratios are expressed in the same units. For more details, see FAQ 2.5.

### 2. Study Design and Results

#### 2.1. What phenotypes are included in the Polygenic Index Repository? How did you choose the phenotypes?

In the initial release of the Repository, we constructed polygenic indexes (PGIs) for 47 phenotypes in 11 datasets, using a consistent methodology. In the second release, we expanded the Repository to PGIs for 61 phenotypes in 20 datasets. The phenotypes (listed in Table 1 of (Alemu et al., 2025) can be categorized into eight somewhat overlapping groups:

- anthropometric (height and body mass index);
- biomarkers (cholesterol, triglycerides, and blood pressure);
- cognition and education (including number of years of formal schooling and performance on cognitive tests, as well as Alzheimer's disease);

- fertility and sexual development (including number of children for women, and age at first menses);
- health (the second largest category, which includes diseases such as Type II diabetes and coronary artery disease, other health phenotypes such as getting migraines and being allergic to pollen, and self-rated overall health);
- personality, well-being, and skills (the largest category, which includes a miscellaneous collection of phenotypes, including risk tolerance, subjective well-being, physical activity, and musical beat synchronization);
- psychiatric (including depressive symptoms and bipolar disorder); and
- substance use (including several alcohol and smoking-related behaviors, as well as cannabis use).

The set of 61 phenotypes we studied was selected from a larger set of 78 phenotypes; we did not create PGIs for the other phenotypes because statistical calculations indicated that, based on the GWAS results we had available, a PGI was predicted to explain less than 1% of the variation across individuals. Although the specific threshold of 1% is somewhat arbitrary (but see further discussion in FAQ 2.3 below), PGIs with low predictive power are less useful and more likely to generate misleading results (such as false positives) if used.

#### 2.2. How did you create these polygenic indexes?

In order to construct the polygenic indexes (PGIs), we combined GWAS results from three sources. First, for the 47 phenotypes where we could find previously published GWAS, we obtained the publicly available results. Second, we collaborated with the personal genomics company 23andMe, which contributes to academic research by analyzing the data of customers who consent to participate in research. For this paper, 23andMe provided GWAS results for 44 phenotypes, 9 of which had not previously been published. Third, for 46 phenotypes, we conducted a GWAS ourselves in the UK Biobank, a large-scale biomedical database accessible to researchers. When more than one of these sources of GWAS results was available for a phenotype, we combined the GWAS results together using a statistical method called meta-analysis.

In the first (but not the second) release of the Repository, we constructed “multi-trait PGIs” using GWAS results for multiple phenotypes (Turley et al., 2018). These PGIs are often more predictive than a standard “single-trait polygenic index” constructed from GWAS results from a single phenotype (FAQ 1.4), but the results from analyzing multi-trait PGIs are sometimes more difficult to interpret (FAQ 2.6).

#### 2.3. How predictive are the polygenic indexes in the Repository?

To assess the predictive power of the polygenic indexes (PGIs), in the most recent paper (Alemu et al., 2025) we used data from 3 of the 20 participating datasets (those for which we had access to both the phenotype and genotype data we needed to construct the PGIs). In each of these 3 datasets, we calculated the predictive power of every PGI for which the dataset contained data on the relevant phenotype (see FAQ 2.1).

The predictive power of the PGIs varies substantially across the phenotypes and validation datasets. The PGI for height has the greatest predictive power. It predicts 36% to 43% of the variation across individuals, depending on the validation dataset. In the UK Biobank, the polygenic indices for the blood phenotypes have predictive powers ranging from 11% to 23%, although the predictive powers are notably lower in the other two validation datasets, which have less precise measures of blood phenotypes. Next are the polygenic indices for body mass index (BMI) and educational attainment, whose predictive power ranges from 8% to 14% in our validation datasets. Two other phenotypes—cognitive performance and age at first menses—have a PGI with predictive power in the range of 7% to 13%. Among the least predictive are the PGIs for autism spectrum disorder, number ever born (for women), satisfaction with family, and satisfaction with friendships, whose predictive powers in our validation datasets range from 0.01% to 0.9% (they were included because their predictive power was statistically expected to exceed 1%; see FAQ 2.1). The predictive powers for the other PGIs in the Repository lie mostly between 1% and 9%.

Although the effects explained by these PGIs are small-to-modest, they can nevertheless be useful in research. For instance, the environmental factors studied in economics research typically have predictive power less than 5%, and often 1% or less. Among the strongest predictors of educational attainment is family socioeconomic status, which has predictive power of roughly 15%. In a standard categorization used in psychology (Cohen, 1992) predictive power less than 9% is "small" while predictive power greater than 25% (rarely attained in psychological research) is "large." We caution, however, that these comparisons of the effect sizes of PGIs and environmental influences aren't apples-to-apples because researchers usually study one particular environmental factor or many on a phenotype, whereas a PGI summarizes the predictive power of SNPs across the genome. As discussed further in FAQ 3.3, for social and behavioral phenotypes, the sum of all environmental (i.e., non-genetic) influences substantially outweigh the sum of all genetic influences that a PGI aims to capture.

As we discuss in FAQ 3.3, an individual's PGIs (even for height) do *not* very accurately predict that *individual's* phenotypes. However, PGIs are useful for *scientific studies* (including social science, health research, etc.) that are concerned with aggregate population trends and *averages* rather than with individual phenotypes. For example, for a PGI that predicts 1% of the variation across individuals, studies of its association with other variables can be well powered in sample sizes as small as 785 individuals; 19 out of the 20 datasets participating in the Repository have sample sizes larger than that.

A major goal of the Polygenic Index Repository is to enable other research that is valuable to social scientists and health researchers. Such studies are already being conducted with some PGIs (see FAQ 1.10). For some phenotypes, the PGIs in the Repository are more predictive than those that were previously possible to construct; examples include having asthma/eczema/rhinitis, number of cigarettes smoked per day, having migraines, nearsightedness, self-reported physical activity, self-rated overall health, and subjective well-being (i.e., self-reported happiness or life satisfaction). For other phenotypes, no PGI of any predictive strength were available prior to the Repository because there had been no large, publicly available, published GWASs for those phenotypes; examples include childhood reading, self-rated math ability, and self-reported narcissism, and several allergies including to pollen.

#### 2.4. How does the Repository enable "causal polygenic index studies"?

One goal of the paper accompanying the release of version 2 of the Repository (Alemu et al., 2025) is to promote causal polygenic index (PGI) studies. It does so in two ways. First, in datasets that do not have genetic data on parents but do have genetic data on other family members, such as siblings, we imputed parental genetic data—meaning that we used the laws of genetic inheritance to make optimal statistical guesses about parents' genotypes (Young et al., 2022) and then constructed parental polygenic indexes (PGIs). Previous research has shown that these parental PGIs constructed from imputed parental genetic data can be used in causal PGI studies as if the parental PGIs were constructed from actual parental genetic data, and the statistical results remain valid. Version 2 of the Repository makes these parental PGIs available to researchers so they can conduct causal PGI studies. Second, the paper aims to help clarify the interpretation of causal PGI studies, and it provides a formula for assessing how much such a study will underestimate causal effects of genetic variants.

#### 2.5. What is the "measurement-error-corrected estimator"? How will it and the Repository improve comparability of results across future studies?

The measurement-error-corrected estimator was introduced in the paper accompanying version 1 of the Polygenic Index Repository (Becker et al., 2021). To understand this tool, it is helpful to imagine the theoretically ideal PGI that could result from an infinitely large GWAS. In the paper, we call the predictor that would result from this ideal GWAS the "additive SNP factor." The actual PGIs that exist in the world are "noisy" measures of, and therefore only proxies for, this additive SNP factor. The signal-to-noise ratio of a PGI—i.e., the extent to which it reflects the additive SNP factor—is determined by the sample size of the GWAS from which the PGI is constructed (a larger GWAS leads to less noise and therefore a higher signal-to-noise ratio). The fact that the PGI is noisy distorts the results of most analyses that use the PGI (relative to what the results would be with the ideal predictor). These distortions can lead researchers to reach incorrect conclusions. For example, in an analysis of how genes and environments interact in influencing some phenotype, the noise in the PGI will usually cause a researcher to underestimate how strongly genes and environments interact.

Moreover, as discussed in FAQ 1.9, there are many reasons why two PGIs for the same phenotype could differ from each other, including differences in the GWAS that the PGI is based on and different methods for constructing the PGI. Many of these differences among GWASs produce differences in the signal-to-noise ratios of their resulting PGIs. Two studies using PGIs with different signal-to-noise ratios will, in turn, have results that are distorted to differing degrees, reducing comparability of results across studies that use the PGIs.

The "measurement-error-corrected estimator" we derive in the paper enables researchers to conduct analyses *without* the distortion that comes from the noise. It works because we (often) have a good estimate of how much noise a given PGI has. We can use that information to calculate what the results of an analysis would have been if the PGI had no noise. The estimator improves comparability of results across papers because it avoids the distortions in results that arise from having a noisy PGI. Rather than being distorted to different degrees, two studies using PGIs with different signal-to-noise ratios that use our estimator will both have undistorted results. We have made available the software for this estimator. We will maintain and provide user support for this software.

Moreover, across all the PGIs and across all the datasets participating in the Repository, we constructed the PGIs in a uniform way. To the extent that future studies use the PGIs from the Repository, their results will therefore be more comparable.

#### 2.6. What is in the User Guide that accompanies the Repository?

Along with the polygenic indexes (PGIs), we have distributed to the participating datasets a User Guide. Data providers distribute this User Guide to researchers as part of the Repository. The User Guide contains technical details about the construction of the PGIs, as well as details about data and software availability. It also describes a set of key interpretational considerations that researchers should keep in mind when analyzing PGIs. These include when to use a single-trait versus multi-trait PGI (see FAQ 2.1), reasons why associations between a PGI and an phenotype generally cannot be interpreted as causal unless parental PGIs are controlled for (see FAQs 1.6 & 1.8), and in what sense they can be interpreted as causal when parental PGIs *are* controlled for (see FAQ 1.8). Finally, the User Guide contains a discussion of six "interpretational considerations" that we urge researchers who use PGIs to consider as part of the responsible conduct and communication of their research (see FAQ 3.6).

#### 2.7. Who can access the Repository polygenic indexes, and how?

Researchers can access the Repository polygenic indexes (PGIs) through the data access procedures for each of the datasets participating in the Repository. These are summarized in the Supplementary Note of each paper and kept up to date on the SSGAC's webpage (https://www.thessgac.org/data-access-procedures). Typically, data providers require researchers to submit a brief a description of the planned research and to sign a Data Use Agreement. The Data Use Agreement usually requires researchers to agree to protect the confidentiality of individuals in the dataset and, to that end, to analyze the data on computers that satisfy certain security protocols.

We provided the PGIs we created to the 20 datasets participating in the Repository, so that the data providers can distribute them to users of the datasets. We designed the Repository this way for three reasons (corresponding to problems #1, #2, and #3 in FAQ 1.10; problem #4 is addressed by using a consistent methodology for constructing the PGIs). First, because we are making available the PGIs (rather than the GWAS results from which they are constructed), researchers do not need to spend time constructing the PGIs from GWAS results. Second, for many phenotypes, the PGIs we construct are based on more data than are in the largest previously published GWAS. Because the Repository PGIs for those phenotypes are based on more data, they are more accurate (i.e., predictive) than PGIs that could be constructed based only on publicly available GWAS results. Third, we tailored the PGIs we constructed to each of the 20 datasets. Specifically, we ensured that for a given dataset, its PGIs were *not* based on GWAS results that included that dataset (which would have led to "sample overlap" that would make it problematic to use the PGI with that dataset).

#### 2.8. How will the Repository be updated?

We plan to continue to update the Repository regularly as new GWASs are published or new data become available with which we can conduct our own GWAS. The updates will increase the predictive power of PGIs already in the Repository, as well as expand the set of phenotypes for which PGIs are available. We also expect to include additional datasets whose stewards want to participate in the Repository and make their data broadly available to the research community.

#### 3. Ethical and social implications of the study

##### 3.1. Do GWAS or the polygenic indexes they produce identify the gene—or genes—“for” a particular phenotype?

No. GWAS of complex phenotypes identify *many* SNPs that are associated with a phenotype like height or educational attainment. Although it was once believed that scientists would discover numerous strong one-to-one associations between specific genes and phenotypes, we have known for a number of years that the vast majority of human phenotypes are complex and are influenced by thousands of genes, each of which alone tends to have a small influence on the relevant phenotype.

Furthermore, many complex phenotypes are also influenced by parts of the genome that are not genes at all but instead serve to regulate genes (e.g., influencing when a gene is turned on or off). Genes typically contain many SNPs (often dozens or hundreds, in some cases thousands), and there are even more SNPs outside of genes than inside genes. Complex phenotypes are often influenced by hundreds of thousands of SNPs.

Although the GWASs that produced the polygenic indexes included in the Repository did find several SNPs that are associated with particular phenotypes, we believe that characterizing these as “genes for X”—or, more accurately—“SNPs for X” (e.g., educational attainment, height) is still likely to mislead, for many reasons, and we urge researchers and reporters to avoid this usage.

As an example, consider the phenotype of educational attainment. First, most of the variation in people’s educational attainment is accounted for by social and other environmental factors, not by additive genetic effects (See FAQ 3.3). “Genes for educational attainment” might be read to imply, incorrectly, that genes are the strongest predictor of variation in educational attainment.

Second, the SNPs that are associated with educational attainment are also associated with many other things. These SNPs are no more “for” educational attainment than they are “for” the other phenotypes with which they are associated.

Third, the “predictive” power (see FAQ 1.7) of each individual SNP that we identify is very small. Our previous work (Okbay et al., 2022) has shown that genetic associations with educational attainment are comprised of thousands, or even millions, of SNPs, each of which has a tiny effect size. Each SNP is therefore weakly associated with, rather than a strong influence on, educational attainment. Science writing that refers to “genes for educational attainment” can be misunderstood to imply that there exists a strong predictive relationship of each gene (or, more accurately, SNP).

Fourth, environmental factors can increase or decrease the impact of specific SNPs (see FAQ 3.3). Put differently, even if a SNP is associated with higher or lower levels of educational attainment *on average*, it may have a much larger or smaller effect depending on environmental conditions. Indeed, in one of our GWAS of educational attainment (J. J. Lee et al., 2018) and elsewhere, we report exploratory analyses that provide evidence of such gene-environment interactions. Educational attainment couldn’t even exist as a meaningful object of measurement if we didn't have schools, and having schools introduces societal mechanisms that influence who goes to them. Accordingly, genetic associations with educational attainment necessarily will be mediated by societal systems and therefore genetic variation should be expected to interact with environmental factors when it influences social phenomena, such as educational attainment. "Genes for educational attainment" suggests a stability in the relationship between these genes and the phenotype of educational attainment that does not exist.

Finally, SNPs do not affect educational attainment directly. As described in our previous work (J. J. Lee et al., 2018), the genes identified as associated with educational attainment tend to be especially active in the brain and involved in neural development and neuron-to-neuron communication. The "predictive" power (see FAQ 1.7) of SNPs on educational attainment may therefore be the result of a long process starting with brain development, followed by the emergence of particular psychological traits (e.g., cognitive abilities and personality). These traits may then lead to behavioral tendencies as well as experiences and treatment by parents, peers, and teachers. All of these factors may additionally interact with the environment in which a person lives. Eventually these traits, behaviors, and experiences may influence (but not completely determine) educational attainment.

#### 3.2. Do polygenic indexes show that these phenotypes are determined, or fixed, at conception?

Absolutely not. Social and other environmental factors account for most variation in most of the phenotypes for which the Repository contains polygenic indexes. But even *if* it were true that genetic factors accounted for *all* of the differences among individuals in a phenotype, it would *still* not follow that an individual's phenotype is "determined" at conception. There are at least three reasons for this.

First, some genetic effects may operate through environmental channels (Jencks, 1980). Again, consider educational attainment as an example. Suppose—hypothetically— that some of the SNPs in the index help students to memorize and, as a result, to become better at taking tests that rely on memorization. In this example, changes to the intermediate environmental channels—the type of tests administered in schools—could have large effects on individuals' educational attainment, even though individuals' genome would not have changed. Certain SNPs may not be associated with educational attainment *at all* if schools did not use tests that rely on memorization. More generally, the PGI for educational attainment in the Repository might be less predictive if the education system were organized differently than it is at present (see also FAQ 3.3).

Second, even if the genetic associations with educational attainment operated entirely through non-environmental mechanisms that are difficult to modify (such as direct influences on the formation of neurons in the brain and the biochemical interactions among them), there could still exist powerful environmental interventions that could change the genetic relationships. In a famous example suggested by the economist Arthur Goldberger, even if all variation in unaided eyesight were due to genes, there would still be enormous benefits from introducing eyeglasses (Goldberger, 1979). Similarly, policies such as a required minimum number of years of education and dedicated resources for individuals with learning disabilities can increase educational attainment in the entire population and/or reduce differences among individuals.

Third, even if the genetic effects on a phenotype were not influenced by changes in the environment, those environmental changes themselves could still have a major impact on the phenotype in the population as a whole. For example, if young children were given more nutritious diets, then everyone's school performance might improve, and college graduation rates might increase. Or consider the phenotype of height: 80%-90% of the variation across individuals in height is due to genetic factors. Yet the current generation of people is much taller than past generations due to changes in the environment such as improved nutrition.

##### 3.3. Can the polygenic indexes from the Repository be used to accurately predict a particular person's phenotypes?

No. While the "predictive" power (see FAQ 1.7) of our polygenic indexes (PGIs) makes most of them useful in research for some purposes (see FAQ 2.3), these PGIs *fail to predict* the majority of variation across individuals. Even the height PGI—the one with the greatest predictive power—fails to predict 60% of the variation in height among people.

Indeed, an important message of a number of our earlier papers is that DNA does *not* "determine" an individual's behavioral and social phenotypes, for at least four reasons: First, in the environments in which the phenotypes have been measured, other studies have estimated that the additive effects of SNPs will only ever account (even with arbitrarily large samples used to construct PGIs) for a minority of the variation across individuals in the phenotypes we study. For example, we estimate that the theoretical upper bound for additive effects of SNPs would account for 46% of the variation in height, 24% in body mass index, 20% in age at first menses, and less than 10% for most of the social/behavioral phenotypes we study. So even a hypothetical PGI that perfectly reflects the additive SNP factor (see FAQ 2.5) is likely to only explain a small fraction of the variation across individuals. Second, *today's* PGIs are *not* perfect; they are only able to predict a fraction of that already small fraction of hypothetical cross-sectional predictive power. Third, since SNPs matter more or less depending on environmental context (see FAQ 3.2), a PGI might be less (or more) predictive for individuals in some environments than for individuals in others. Finally, and similarly, polygenic predictions only hold for as long as the environment in which they were developed remains substantially the same.

To illustrate these final two reasons, consider the example of educational attainment (for which we have included a PGI in the Repository and on which we have done previous research): if the pedagogy underlying the educational system in which the GWAS that produced the PGI was conducted is substantially different than the pedagogy of the *different population* to which that PGI is being applied, the PGI may be less (or, conceivably, more) predictive in this second population (for an example, see FAQ 3.2). The same is true if the PGI is applied to the same population, but at a *later time* when the pedagogy has changed substantially. Just as eyeglasses allow those genetically predisposed to poor vision to have nearly perfect vision, innovations in education (say, an innovation that makes education highly engaging, thus mitigating the risk to those with SNPs associated with lower ability to pay attention or maintain self-control) might result in those with lower PGIs now achieving just as much education, on average, as those with higher PGIs.

As sample sizes for GWAS continue to grow, it will likely be possible to construct PGIs for many phenotypes whose predictive power comes closer to the total amount of variation that is theoretically predictable from additive effects of common SNPs for those phenotypes (the upper bounds given above). Even these levels of predictive power would pale in comparison to some other scientific predictors. For example, professional weather forecasts correctly predict (one day ahead) about 95% of the variation in day-to-day temperatures. Weather forecasters are therefore vastly more accurate forecasters than social science geneticists will ever be.

PGIs created by GWASs are increasingly used by commercial and research direct-to-consumer platforms to predict individual phenotypes. We recognize that returning individual genomic "results" can be a fun way to engage people in research and other projects and has at least the theoretical potential to stoke their interest in, and educate them about, genomics and how genes and environments interact. But it is important that participants/users understand that, at present, most of these individual results, including all social and behavioral phenotypes, are not *meaningful* predictions (in the sense that they generally have very little predictive power at the individual level). Failure to make this point clear risks sowing confusion and undermining trust in genetics research.

#### 3.4. Can the polygenic indexes accurately be used for research studies in non-European-genetic-ancestry populations?

No. We constructed polygenic indexes (PGIs) only for individuals classified as "European genetic ancestries." (The precise definition of "European genetic ancestry" differs in different datasets, but it usually means that a person's pattern of genetic variation across the genome is statistically close to the average pattern from a "reference sample" for some European country. The reference samples used by geneticists are based on samples of people who live in the European country today and whose recent ancestors also lived in that country.) Therefore, the Polygenic Index Repository only includes PGIs for these individuals.

The main reason we only constructed PGIs for these individuals is that the PGIs are likely to be much less predictive—and hence much less useful—in a sample of people of non-European genetic ancestries. That is because our original GWAS data was obtained from samples of people with European genetic ancestries, and GWAS results have been found to have only limited portability across genetic ancestries (Belsky et al., 2013; Domingue et al., 2015, 2017; Martin et al., 2017; Vassos et al., 2017). There are a number of reasons for the limited portability. For one thing, the set of SNPs that are associated with a phenotype in people of European genetic ancestries is unlikely to overlap closely with the set of SNPs associated with the phenotype in people of non-European genetic ancestries. And even if a given SNP is associated in both ancestry groups, the effect size—in other words, the strength of the association—will almost surely differ. This is primarily because linkage disequilibrium (LD) patterns (i.e., the correlation structure of the genome) vary by genetic ancestry. This means that some SNP may be associated with the phenotype because the SNP is in LD (i.e., correlated) with a SNP elsewhere in the genome that causally affects the phenotype (see FAQ 1.6). If the strength of the correlation is greater in one ancestry group than in another, then the size of the association will be larger in that ancestry group. Moreover, even if LD patterns were similar in each ancestry group, the association may differ in different groups because environmental conditions differ (see FAQ 1.7). The fact that there are differences across ancestry groups in the set of associated SNPs and their effect sizes means that the weights for constructing PGIs in individuals with European genetic ancestries (FAQ 1.4) would be the "wrong" weights for non-European-genetic-ancestry individuals. For a more extensive, excellent discussion of these and related issues, see Graham Coop's blog post, "Polygenic scores and tea drinking."

Unfortunately, this attenuation of predictive power means that for non-European-genetic-ancestry populations, many of the benefits of having a PGI available will have to wait until large GWAS studies are conducted using samples from these populations. (Currently, most large genotyped samples are of European genetic ancestries.) Future versions of the Polygenic Index Repository will include PGIs for non-European-genetic-ancestry populations, once it becomes possible to produce PGIs with adequate predictive power.

##### 3.5. Could research on polygenic indexes lead to discrimination against, or stigmatization of, people with higher or lower polygenic indexes for certain phenotypes? If so, why facilitate the spread of polygenic indexes?

Unfortunately, like a great deal of research—including, for instance, research identifying genomic variation associated with increased cancer risk or research identifying environmental predictors of socially disfavored phenotypes—the results can be misunderstood and misapplied. This includes being used to discriminate against those with higher or lower polygenic indexes (PGIs) for certain phenotypes (e.g., in insurance markets). Nevertheless, for a variety of reasons, in this instance, we do not think that the best response to the possibility that useful knowledge could be misused is to refrain from producing the knowledge. Moreover, many researchers already have access to and use PGIs; against this background, the Repository helps ensure that a much wider array of researchers have the same opportunity to access and probe these research tools, and also that the PGIs themselves will be more accurate. Here, we briefly discuss some of the broad potential benefits of this research. We then describe what we see as our ethical duty as researchers conducting this work.

First, one benefit of conducting social-science genetics research in ever larger samples is that doing so allows us to correct the scientific record. An important theme in our earlier work has been to point out that most existing studies in social-science genetics that report genetic associations with behavioral phenotypes have serious methodological limitations, fail to replicate, and are likely to be false-positive findings (Benjamin et al., 2012; Chabris et al., 2012, 2015). This same point was made in an editorial in *Behavior Genetics* (the leading journal for the genetics of behavioral phenotypes), which stated that "it now seems likely that many of the published [behavior genetics] findings of the last decade are wrong or misleading and have not contributed to real advances in knowledge" (Hewitt, 2012). One of the most important reasons why earlier work has generated unreliable results is that the sample sizes were far too small, given that the true effects of individual SNPs on behavioral phenotypes are tiny. Pre-existing claims of genetic associations with complex social-science phenotypes have reported widely varying effect sizes, many of them purporting to "predict" as much of the variation across individuals as do the PGIs we construct in this paper that aggregate the effects of millions of SNPs.

Second, behavioral genetics research also has the potential to correct the *social* record and thereby to help *combat* discrimination and stigmatization. For instance, overestimating the role of genetics can be damaging, and the present work can help debunk the myth of genetic determinism. By quantifying how various phenotypes are predicted by genetic data, we show that for all of the phenotypes we study, the genetic data can explain a very small fraction of the variation across individuals (see FAQ 2.3). By clarifying the *limits* of deterministic views of complex phenotypes, recent behavioral genetics research—if communicated responsibly—could make appeals to genetic justifications for discrimination and stigmatization *less* persuasive to the public in the future.

Third, behavioral genetics research has the potential to yield many other benefits, especially as sample sizes continue to increase—as briefly summarized in FAQ 1.10. Foregoing this research necessarily entails foregoing these and any other possible benefits, some of which will likely be the result of serendipity. Indeed, very few of the uses of PGIs were anticipated when they were first proposed (Wray et al., 2007).

In sum, we agree with the U.K. Nuffield Council on Bioethics, which concluded in a report (Nuffield Council on Bioethics, 2002, p. 114) that "research in behavioural genetics has the potential to advance our understanding of human behaviour and that the research can therefore be justified," but that "researchers and those who report research have a duty to communicate findings in a responsible manner" (see FAQ 3.6).

#### 3.6. What have you done to mitigate the risks of research using Repository polygenic indexes?

In our view, the responsible behavioral genetics research called for by the Nuffield Council on Bioethics (see FAQ 3.5) includes sound methodology and analysis of data (e.g., only conducting analyses that are adequately powered and, when feasible, preregistering power calculations and planned analyses); a commitment to publish all results, including any negative results; and transparent, complete reporting of methodology and findings in publications, presentations, and communications with the media and the public. A critical aspect of the latter is particular vigilance regarding what research results do—and do not—show, and how polygenic indexes (PGIs) can—and cannot—be appropriately used. In an effort to reduce the risk that its results might be misinterpreted by readers, misreported by the media, or misused, the SSGAC has developed and publicly posted FAQs like this document with every major paper it has published since its first paper in 2013.

Some of us have also written about the problems with using PGIs for certain purposes (Meyer, Appelbaum, et al., 2023; Meyer et al., 2024; Meyer, Tan, et al., 2023; Turley et al., 2021) and the importance of responsibly communicating the results of social science genomics research (Meyer, Appelbaum, et al., 2023).

In addition, the SSGAC requires researchers who download the SNP weights for constructing PGIs to agree to Terms of Service meant to promote responsible research and communication. For example, we ask researchers to acknowledge "I understand that comparisons of genetically predicted phenotype levels across ancestral groups are usually scientifically confounded due to the effects of linkage disequilibrium, gene-environment correlation, gene-environment interactions, and other methodological problems" (see FAQ 3.5).

These Terms of Service stem from the observation that SNP associations are not necessarily causal (see FAQ 1.6) and depend on the environment of the individuals included in the GWAS (see FAQ 1.7). Different ancestry groups arise in the population because they became partially separated from each other many generations ago, for example, due to geographic factors or social forces. When two groups are geographically or socially separated, they also face different environments, which not only may have direct effects on certain phenotypes (such as disease risk) but may also change the strength of the association between the phenotypes and certain SNPs. Therefore, when individuals from two ancestry groups have different average phenotypes, it is extremely difficult to identify whether the difference is due to average genetic differences between the groups or to the different environments faced by the groups. For this reason, it is scientifically invalid to make general statements about ancestry group differences based on SNP associations identified in a GWAS. (Also see FAQ 3.2.) The Terms of Service also require users to securely store the data and to immediately report any breach of the Terms.

Finally, we have developed and provided to participating data providers a User Guide to be distributed to researchers who use Repository PGIs (see FAQ 2.6). We will also provide the User Guide to researchers who download the SNP weights. One section of the User Guide discusses six "interpretational considerations" that are likely to arise when conducting research with PGIs and which we urge researchers to seriously consider as a critical part of responsibly conducting and communicating their research. One recurring ethical concern about genetic research is the tendency for its predictive power to become exaggerated in the media and in the public's minds, at the expense of a more nuanced understanding of how genes and environment interact, the importance of environmental influences, and the ability of interventions to improve outcomes. Many of the interpretational considerations we discuss in the User Guide involve how to anticipate and address potential confounds and how to navigate complex questions about causality and ensure responsible communication of causality.

For instance, the User Guide cautions researchers to appreciate and communicate that associations between a PGI and a phenotype may operate through *environmental* (rather than biological) mechanisms (see FAQs 3.2 and 3.3).
