## Supplementary Methods and User Guide for "An Updated Polygenic Index Repository: Expanded Phenotypes, New Cohorts, and Improved Causal Inference"

August 3, 2025

### 1 Theoretical Framework

Here we expand on and provide proofs of the claims in the paper's theoretical framework. To keep the supplementary section self-contained, we repeat the setup in the paper while also providing additional details.

Consider a phenotype $y_i^*$. The allele count for individual $i$ and his/her parents at SNP $j$ is denoted by $x_{ij}^* \in \{0,1,2\}$ and $x_{p,ij}^* \in \{0,1,2,3,4\}$ respectively. Without loss of generality, we use mean-centred transformations of the phenotype and allele counts, such that $y_i = y_i^* - \mathbb{E}(y_i^*)$, $x_{ij} = x_{ij}^* - \mathbb{E}(x_{ij}^*)$ and $x_{p,ij} = x_{p,ij}^* - \mathbb{E}(x_{p,ij}^*)$, for each SNP $j$. Note that $x_{p,ij} = x_{ij} + x_{n,ij}$, where $x_{n,ij}$ denotes non-transmitted alleles. Denote a vector of mean-centered allele counts of $i$ and his/her parents across $J$ SNPs by $\mathbf{x}_i = (x_{i1}, x_{i2}, ..., x_{iJ})$ and $\mathbf{x}_{p,i} = (x_{p,i1}, x_{p,i2}, ..., x_{p,iJ})$, respectively. Similarly, a vector of non-transmitted alleles is denoted by $\mathbf{x}_{n,i} = (x_{n,i1}, x_{n,i2}, ..., x_{n,iJ})$.

For simplicity, here we assume that the causal effects of the genetic variants are homogeneous across individuals. In a more realistic scenario where causal effects vary across individuals, the causal parameters below should be interpreted as weighted averages. Individuals with heterozygous parents receive more weight because the variance in their genotype is greater (see refs.[1]and[2]). In this model, "causal effect of a genetic variant" is a convenient shorthand for such a weighted average.

Suppose that the phenotype $\mathbf{y}$ is determined by

$$y_i = \mathbf{x}_i\boldsymbol{\gamma} + \mathbf{x}_{p,i}\boldsymbol{\theta} + u_i. \tag{1}$$

The coefficient on the parental genotype vector, $\boldsymbol{\theta}$, captures (a linear approximation to) causal effects of parental genotypes on $y_i$ that operate through parental phenotypes that affect individual $i$'s environment - which we call *parental genetic effects* (the more common term is "parental indirect genetic effects") - but it also captures confounding from gene-environment correlation, population stratification, and effects of genetic variants not included in $\mathbf{x}_i$ but that are correlated with $\mathbf{x}_i$ due to non-random mating (assortative mating and population structure)[3]. In contrast, because $\mathbf{x}_i$ is randomly assigned conditional on $\mathbf{x}_{p,i}$, the coefficient vector $\boldsymbol{\gamma}$ is free from confounding from gene-environment correlation: it is the best linear approximation to the causal genetic component, given the set of SNPs included in the analysis[2]. It captures causal effects of SNPs included in $\mathbf{x}_i$, and it includes causal effects of genetic variants not included in $\mathbf{x}_i$ to the extent that they are correlated with included SNPs. Note that if controls were included in (1), $\boldsymbol{\theta}$ could change but $\boldsymbol{\gamma}$ would remain unchanged as long as those controls are causally prior to $\mathbf{x}_i$ (i.e., the controls are not themselves causally affected by $\mathbf{x}_i$).

We define the the *causal additive SNP factor* as

$$g_i^c = \frac{\mathbf{x}_i\boldsymbol{\gamma}}{sd(\mathbf{x}_i\boldsymbol{\gamma})}, \tag{2}$$

The causal additive SNP factor maximizes the variance explained in $y_i$ conditional on the parental genotypes, and in that sense, it represents the overall causal effects of genetic variants as faithfully as possible, conditional on the SNPs included in the analysis and on using a linear approximation. We refer to the variance explained by the causal additive SNP factor, denoted $h_\gamma^2 \equiv Var(\mathbf{x}_i\boldsymbol{\gamma})/Var(y_i)$, as the *causal SNP heritability.* (If $\mathbf{x}_i$ and $\mathbf{x}_{p,i}$ contained all genetic variants in the genome, then $g_i^c$ would be the *causal additive genetic factor*, and the phenotypic variance explained by $\mathbf{x}_i\boldsymbol{\gamma}$ would be the *narrow-sense heritability.*)

Researchers cannot use the causal additive SNP factor as a PGI because $\boldsymbol{\gamma}$ is unknown. Although $\boldsymbol{\gamma}$ can be estimated from summary statistics of a family-based GWAS[4,5], the estimates are noisy, mostly because sample sizes are relatively small. Indeed, even though genetic effects are estimated with bias in a standard GWAS, at present, the mean squared error of the estimates from standard GWAS currently tend to be far smaller than the unbiased estimates from family-based GWAS due to the smaller estimation error[6]. Consequently, PGIs based on standard GWAS have much greater predictive power. For that reason, PGIs used in practice, including those in the Repository, are constructed from standard (non-family-based) GWAS. To formalize these PGIs, consider the population regression of $y_i$ on $\mathbf{x}_i$:

$$y_i = \mathbf{x}_i\boldsymbol{\mu} + \xi_i, \tag{3}$$

where $\boldsymbol{\mu}$ captures both the causal effect and the parental association since $y_i = \mathbf{x}_i\boldsymbol{\gamma} + (\mathbf{x}_i + \mathbf{x}_{n,i})\boldsymbol{\mu} + u_i = \mathbf{x}_i(\boldsymbol{\gamma} + \boldsymbol{\theta}) + \mathbf{x}_{n,i}\boldsymbol{\theta} + u_i = \mathbf{x}_i\boldsymbol{\mu} + \mathbf{x}_{n,i}\boldsymbol{\theta} + u_i = \mathbf{x}_i\boldsymbol{\mu} + \xi_i$. The coefficient vector $\boldsymbol{\mu}$ maximizes the variance explained in $y_i$, but it does not have a causal interpretation because $\boldsymbol{\mu}$ will partially capture confounding, including gene-environment correlation (due to its $\boldsymbol{\theta}$ component). This is consistent with Trejo et al.[7] who show that the causal genetic effect is overestimated if parental genetic effect is not controlled for.

Standard methods for constructing SNP weights from GWAS summary statistics, such as SBayesR[8] (which we use) and LDpred[9], generate a SNP weight vector $\mathbf{w} = \hat{\boldsymbol{\mu}}$ that is a consistent estimate of $\boldsymbol{\mu}$. We assume that $\hat{\boldsymbol{\mu}}$ is an unbiased estimate of $\boldsymbol{\mu}$, which we showed in our earlier paper[10] is a good approximation when the GWAS sample size is large. We refer to $\mathbf{x}_i\boldsymbol{\mu}$ as the *associational additive SNP factor*, and the variance explained by it, denoted $h^2_\mu \equiv Var(\mathbf{x}_i\boldsymbol{\mu})/Var(y_i)$ , as the *associational SNP heritability.*

We can write the standardized PGI constructed with the weight vector $\mathbf{w} = \hat{\mu}$ as

$$\hat{g}_i \equiv \frac{\mathbf{x}_i\hat{\boldsymbol{\mu}}}{sd(\mathbf{x}_i\hat{\boldsymbol{\mu}})}. \tag{4}$$

The same weights are used to construct the parental PGI:

$$\hat{g}_{p,i} \equiv \frac{\mathbf{x}_{p,i}\hat{\boldsymbol{\mu}}}{sd(\mathbf{x}_{p,i}\hat{\boldsymbol{\mu}})} \tag{5}$$

We can write the standardized child's and parental PGIs as:

$$\hat{g}_i = \frac{g_i^c + q_i + \varepsilon_i}{sd(g_i^c + q_i + \varepsilon_i)} = \frac{g_i^c + q_i + \varepsilon_i}{\alpha} = \frac{g_i + \varepsilon_i}{\alpha} \tag{6}$$

$$\hat{g}_{p,i} = \frac{g_{p,i}^c + q_{p,i} + \varepsilon_{p,i}}{sd(g_{p,i}^c + q_{p,i} + \varepsilon_{p,i})} = \frac{g_{p,i}^c + q_{p,i} + \varepsilon_{p,i}}{\alpha} = \frac{g_{p,i} + \varepsilon_{p,i}}{\alpha}, \tag{7}$$

where $g_i = g_i^c + q_i$ and $g_{p,i} = g_{p,i}^c + q_{p,i}$ such that

$$\varepsilon_i = \frac{\mathbf{x}_i(\hat{\boldsymbol{\mu}} - \boldsymbol{\mu})}{sd(\mathbf{x}_i\boldsymbol{\gamma})} \tag{8}$$

$$\varepsilon_{p,i} = \frac{\mathbf{x}_{p,i}(\hat{\boldsymbol{\mu}} - \boldsymbol{\mu})}{sd(\mathbf{x}_{p,i}\boldsymbol{\gamma})} \tag{9}$$

$$q_i = \frac{\mathbf{x}_i(\boldsymbol{\mu} - \boldsymbol{\gamma})}{sd(\mathbf{x}_i\boldsymbol{\gamma})} = \frac{\mathbf{x}_i\boldsymbol{\theta}}{sd(\mathbf{x}_i\boldsymbol{\gamma})} \tag{10}$$

$$q_{p,i} = \frac{\mathbf{x}_{p,i}(\boldsymbol{\mu} - \boldsymbol{\gamma})}{sd(\mathbf{x}_{p,i}\boldsymbol{\gamma})} = \frac{\mathbf{x}_{p,i}\boldsymbol{\theta}}{sd(\mathbf{x}_{p,i}\boldsymbol{\gamma})} \tag{11}$$

$$g_{p,i}^c = \frac{\mathbf{x}_{p,i}\boldsymbol{\gamma}}{sd(\mathbf{x}_{p,i}\boldsymbol{\gamma})} \tag{12}$$

$$\alpha = sd(g_i^c + q_i + \varepsilon_i) = \frac{sd(\mathbf{x}_i\hat{\boldsymbol{\mu}})}{sd(\mathbf{x}_i\boldsymbol{\gamma})} = sd(g_{p,i}^c + q_{p,i} + \varepsilon_i) = \frac{sd(\mathbf{x}_{p,i}\hat{\boldsymbol{\mu}})}{sd(\mathbf{x}_{p,i}\boldsymbol{\gamma})}. \tag{13}$$

So, $\varepsilon_i$ and $\varepsilon_{p,i}$ are the estimation errors, which we assume to be uncorrelated with $g_i^c$, $g_{p,i}^c$, $q_i$, $q_{p,i}$, $u_i$, and $y_i$. On the other hand, $q_i$ and $q_{p,i}$ are non-classical measurement errors that arise because $\boldsymbol{\mu}$ is estimated in a population GWAS and captures not only the causal genetic effect $\boldsymbol{\gamma}$ but also the parental association $\boldsymbol{\theta}$.

#### 1.1 Special Case

Suppose that the parental associations and causal genetic effects are proportional, so that $\boldsymbol{\theta} = \lambda\boldsymbol{\gamma}$, where $\lambda \geq 0$ is a scaling constant (possibly $\lambda = 0$ if there are no parental genetic effects and no confounding from gene-environment correlation in standard GWAS estimates). In addition, assume that the SNP weights are estimated in a very large GWAS so that there is no estimation error in the SNP weights and therefore no classical measurement error in the PGIs: $\varepsilon_{p,i} = \varepsilon_i = 0$ for all $i$. Then, it is straightforward that

$q_i = \lambda g_i^c$, and $\alpha = \sqrt{Var(q_i + g_i^c)} = \sqrt{Var\Big((1+\lambda)g_i^c\Big)} = (1+\lambda)sd(g_i^c) = (1+\lambda)$. This implies that $\hat{g}_i = \frac{g_i^c + q_i}{\alpha} = \frac{(1+\lambda)g_i^c}{1+\lambda} = g_i^c$.

This implies that using $\hat{g}_i$ and $\hat{g}_{p,i}$ instead of $g_i^c$ and $g_{p,i}^c$ will not introduce any bias in the estimation. Note that this also will imply that $Corr(\hat{g}_i, g_i^c) = 1$, which is unlikely to be the case for most phenotypes[11].

#### 1.2 Bias Derivation

Consider the following model:

$$\phi_i = \beta_c g_i^c + \beta_{p,c} g_{p,i}^c + \mathbf{z}_i \boldsymbol{\zeta}_c + \boldsymbol{\varpi}_i^c \boldsymbol{\delta}_c + \boldsymbol{\varpi}_{p,i}^c \boldsymbol{\delta}_{p,c} + \upsilon_{i,c}, \tag{14}$$

where $\mathbf{z}_i$ is a vector of size $z$ of mean-zero covariates normalized to have $sd(\mathbf{z}_i) = 1$. The model also includes interaction between $\boldsymbol{\varpi}_i^c = g_i^c \mathbf{z}_{int,i}$ and $\boldsymbol{\varpi}_{p,i}^c = g_{p,i}^c \mathbf{z}_{int,i}$, where $\mathbf{z}_{int,i} \subseteq \{z_{i,j} | j = 1, 2, \ldots, z\}$ and it has size $z_{int}$.

We first derive the coefficients from the correct model defined by equation (14).

Denote $\boldsymbol{\beta}_{G^c} = (\beta_c, \boldsymbol{\delta}_c, \beta_{p,c}, \boldsymbol{\delta}_{p,c}, \boldsymbol{\zeta}_c)'$. Then the coefficient vector $\boldsymbol{\beta}_{G^c}$ is equal to

$$\boldsymbol{\beta}_{G^c} = \begin{pmatrix} Var(\boldsymbol{G}_i^c) & Cov(\boldsymbol{G}_i^c, \mathbf{z}_i) \\ Cov(\mathbf{z}_i, \boldsymbol{G}_i^c) & Var(\mathbf{z}_i) \end{pmatrix}^{-1} \begin{pmatrix} Cov(\boldsymbol{G}_i^c, \phi_i) \\ Cov(\mathbf{z}_i, \phi_i) \end{pmatrix} = V_{g^c}^{-1} \begin{pmatrix} Cov(\boldsymbol{G}_i^c, \phi_i) \\ Cov(\mathbf{z}_i, \phi_i) \end{pmatrix}, \tag{15}$$

where $\boldsymbol{G}_i^c = (g_i^c, \boldsymbol{\varpi}_i^c, g_{p,i}^c, \boldsymbol{\varpi}_{p,i}^c)'$ and we denote its size by $\mathbb{G}$, and $Var(\boldsymbol{G}_i^c)$ is the variance-covariance matrix of $\boldsymbol{G}_i^c$.

Now suppose that we only observe $\hat{g}_i$ and $\hat{g}_{p,i}$. So, we estimate the model

$$\phi_i = \beta \hat{g}_i + \beta_p \hat{g}_{p,i} + \mathbf{z}_i \boldsymbol{\zeta} + \hat{\boldsymbol{\varpi}}_i \boldsymbol{\delta} + \hat{\boldsymbol{\varpi}}_{p,i} \boldsymbol{\delta}_p + \nu_i. \tag{16}$$

Denote $\boldsymbol{\beta}_{\hat{G}} = (\beta, \boldsymbol{\delta}, \beta_p, \boldsymbol{\delta}_p, \boldsymbol{\zeta})'$ and $\hat{\boldsymbol{G}}_i = (\hat{g}_i, \hat{\boldsymbol{\varpi}}_i, \hat{g}_{p,i}, \hat{\boldsymbol{\varpi}}_{p,i})$, where $\hat{\boldsymbol{\varpi}}_i = \hat{g}_i \mathbf{z}_{int,i}$ and $\hat{\boldsymbol{\varpi}}_{p,i} = \hat{g}_{p,i} \mathbf{z}_{int,i}$.

Then the coefficient vector $\boldsymbol{\beta}_{\hat{G}}$ is

$$\beta_{\hat{G}} = \begin{pmatrix} Var(\hat{\boldsymbol{G}}_i) & Cov(\hat{\boldsymbol{G}}_i, \mathbf{z}_i) \\ Cov(\hat{\boldsymbol{G}}_i, \mathbf{z}_i) & Var(\mathbf{z}_i) \end{pmatrix}^{-1} \begin{pmatrix} Cov(\hat{\boldsymbol{G}}_i, \phi_i) \\ Cov(\mathbf{z}_i, \phi_i) \end{pmatrix} = \mathbf{V}_{\hat{g}}^{-1} \begin{pmatrix} Cov(\hat{\boldsymbol{G}}_i, \phi_i) \\ Cov(\mathbf{z}_i, \phi_i) \end{pmatrix} \tag{17}$$

Note that

$$\begin{aligned} \begin{pmatrix} Cov(\hat{\boldsymbol{G}}_i, \phi_i) \\ Cov(\mathbf{z}_i, \phi_i) \end{pmatrix} &= \begin{pmatrix} Cov(\hat{g}_i, \phi_i) \\ Cov(\hat{\boldsymbol{\varpi}}_i, \phi_i) \\ Cov(\hat{g}_{p,i}, \phi_i) \\ Cov(\hat{\boldsymbol{\varpi}}_{p,i}, \phi_i) \\ Cov(\mathbf{z}_i, \phi_i) \end{pmatrix} = \begin{pmatrix} Cov(\frac{g_i^c + q_i + \varepsilon_i}{\alpha}, \phi_i) \\ Cov\Big(\frac{(g_i^c + q_i + \varepsilon_i)\mathbf{z}_{int,i}}{\alpha}, \phi_i\Big) \\ Cov(\frac{g_{p,i}^c + q_{p,i} + \varepsilon_{p,i}}{\alpha}, \phi_i) \\ Cov\Big(\frac{(g_{p,i}^c + q_{p,i} + \varepsilon_{p,i})\mathbf{z}_{int,i}}{\alpha}, \phi_i\Big) \\ Cov(\mathbf{z}_i, \phi_i) \end{pmatrix} \\ &= \mathbf{P}^{-1}\Big( \begin{pmatrix} Cov(\boldsymbol{G}_i^c, \phi_i) \\ Cov(\mathbf{z}_i, \phi_i) \end{pmatrix} + \begin{pmatrix} Cov(\boldsymbol{Q}_i, \phi_i) \\ 0_{|z \times \mathbb{G}|} \end{pmatrix} \Big), \end{aligned} \tag{18}$$

where $\mathbf{P} = \begin{pmatrix} diag(\alpha) & 0_{|\mathbb{G} \times z|} \\ 0_{|z \times \mathbb{G}|} & \mathbf{I}_{|z \times z|} \end{pmatrix}$, $diag(\alpha)$ is a diagonal matrix of size $\mathbb{G} \times \mathbb{G}$ with $\alpha$ on its main diagonal, $\mathbf{I}_{|z \times z|}$ is an identity matrix with $z$ rows, $0_{|z \times \mathbb{G}|}$ is a matrix of zeros of size $z \times \mathbb{G}$, and $\boldsymbol{Q}_i = (q_i, q_i \mathbf{z}_{int,i}, q_{p,i}, q_{p,i} \mathbf{z}_{int,i})$.

Importantly, equation (18) is based on the assumption that $\varepsilon_i$ and $\varepsilon_{p,i}$ are uncorrelated with $\mathbf{z}_{int,i}$ and $\phi_i$, which implies that $Cov(\varepsilon_i, \phi_i) = Cov(\varepsilon_{p,i}, \phi_i) = Cov(\varepsilon_i \mathbf{z}_{int,i}, \phi_i) = Cov(\varepsilon_{p,i} \mathbf{z}_{int,i}, \phi_i) = 0$.

Equations (17) and (18) imply that

$$\boldsymbol{\beta}_{\hat{G}} = \mathbf{V}_{\hat{g}}^{-1} \mathbf{P}^{-1} \Big( \begin{pmatrix} Cov(\boldsymbol{G}_i^c, \phi_i) \\ Cov(\mathbf{z}_i, \phi_i) \end{pmatrix} + \begin{pmatrix} Cov(\boldsymbol{Q}_i, \phi_i) \\ 0_{|z \times \mathbb{G}|} \end{pmatrix} \Big) = \mathbf{V}_{\hat{g}}^{-1} \mathbf{P}^{-1} \Big( \mathbf{V}_{g^c} \boldsymbol{\beta}_{G^c} + \begin{pmatrix} Cov(\boldsymbol{Q}_i, \phi_i) \\ 0_{|z \times \mathbb{G}|} \end{pmatrix} \Big). \tag{19}$$

Next, it can be shown that

$$\begin{pmatrix} Cov(\boldsymbol{Q}_i, \phi_i) \\ 0_{|z\times 1|} \end{pmatrix} = \begin{pmatrix} Cov(\boldsymbol{Q}_i, \boldsymbol{G}_i^c) & Cov(\boldsymbol{Q}_i, \mathbf{z}_i) \\ 0_{|z\times \mathbb{G}|} & 0_{|z\times z|} \end{pmatrix} \boldsymbol{\beta}_{G^c} = \mathbb{M}_{q,g^c} \boldsymbol{\beta}_{G^c}. \tag{20}$$

This transforms (19) into

$$\boldsymbol{\beta}_{\hat{G}} = \mathbf{V}_{\hat{g}}^{-1} \mathbf{P}^{-1} \Big( \mathbf{V}_{g^c} + \mathbb{M}_{q,g^c} \Big) \boldsymbol{\beta}_{G^c}. \tag{21}$$

Recall that $\mathbf{V}_{\hat{g}}^{-1} = \begin{pmatrix} Var(\hat{\boldsymbol{G}}_i) & Cov(\hat{\boldsymbol{G}}_i, \mathbf{z}_i) \\ Cov(\hat{\boldsymbol{G}}_i, \mathbf{z}_i) & Var(\mathbf{z}_i) \end{pmatrix}$. It can be shown that

$$\mathbf{V}_{\hat{g}}^{-1} = \mathbf{P}^{-1} \Big( \mathbf{V}_{g^c} + \mathbf{V}_q + \mathbf{V}_E + \mathbb{M}_{q,g^c}^{'} + \mathbb{M}_{q,g^c} \Big) \mathbf{P}^{-1} \ . \tag{22}$$

Hence,

$$\mathbf{V}_{g^c} = \mathbf{P} \mathbf{V}_{\hat{g}} \mathbf{P} - \mathbf{V}_q - \mathbf{V}_E - \mathbb{M}_{q,g^c}^{'} - \mathbb{M}_{q,g^c}, \tag{23}$$

where $\mathbf{V}_q$ is a partitioned matrix of the same size as $\mathbf{V}_{g^c}$ with the first block being the variance-covariance matrix of $\boldsymbol{Q}_i$ such that $\mathbf{V}_q = \begin{pmatrix} Var(\boldsymbol{Q}_i) & 0_{|\mathbb{G}\times z|} \\ 0_{|z\times \mathbb{G}|} & 0_{|z|} \end{pmatrix}$; and $\mathbf{V}_E$ is a partitioned matrix of the same size as $\mathbf{V}_{g^c}$ with the first block being $Var(\mathbb{E}_i)$, the variance-covariance matrix of $\mathbb{E}_i = (\varepsilon_i, \varepsilon_i \mathbf{z}_{int,i}, \varepsilon_{p,i}, \varepsilon_{p,i} \mathbf{z}_{int,i})$, such that $\mathbf{V}_E = \begin{pmatrix} Var(\mathbb{E}_i) & 0_{|\mathbb{G}\times z|} \\ 0_{|z\times \mathbb{G}|} & 0_{|z\times z|} \end{pmatrix}$.

Let us define

$$\Psi = \mathbf{V}_q + \mathbb{M}_{q,g^c} = \begin{pmatrix} Var(\boldsymbol{Q}_i) + Cov(\boldsymbol{G}_i^c, \boldsymbol{Q}_i) & 0_{|\mathbb{G}\times z|} \\ Cov(\mathbf{z}_i, \boldsymbol{Q}_i) & 0_{|z\times z|} \end{pmatrix}. \tag{24}$$

This transforms (21) into

$$\boldsymbol{\beta}_{\hat{G}} = \mathbf{V}_{\hat{g}}^{-1} \mathbf{P}^{-1} \Big( \mathbf{P} \mathbf{V}_{\hat{g}} \mathbf{P} - \mathbf{V}_E - \Psi \Big) \boldsymbol{\beta}_{G^c}. \tag{25}$$

Below we define each component of equation (25).

We start from the component of $\mathbf{V}_E$ from 24, $Var(\mathbb{E}_i)$. Note that we assume that the error terms $\varepsilon_i$ and $\varepsilon_{p,i}$ are independent from all the control variables and the genetic variables. This implies that

$$Var(\mathbb{E}_i) = \begin{pmatrix} Var(\varepsilon_i) & 0_{|1\times z_{int}|} & Cov(\varepsilon_i, \varepsilon_{p,i}) & 0_{|1\times z_{int}|} \\ 0_{|z_{int}\times 1|} & Var(\varepsilon_i) Var(\mathbf{z}_{int,i}) & 0_{|z_{int}\times 1|} & Cov(\varepsilon_i, \varepsilon_{p,i}) Var(\mathbf{z}_{int,i}) \\ Cov(\varepsilon_i, \varepsilon_{p,i}) & 0_{|1\times z_{int}|} & Var(\varepsilon_{p,i}) & 0_{|1\times z_{int}|} \\ 0_{|z_{int}\times 1|} & Cov(\varepsilon_{p,i}, \varepsilon_i) Var(\mathbf{z}_{int,i}) & 0_{|z_{int}\times 1|} & Var(\varepsilon_{p,i}) Var(\mathbf{z}_{int,i}) \end{pmatrix}, \tag{26}$$
$$= Var(\varepsilon) \bigotimes \mathbf{V}_{z_{int}}$$

where $Var(\varepsilon) = \begin{pmatrix} Var(\varepsilon_i) & Cov(\varepsilon_i, \varepsilon_{p,i}) \\ Cov(\varepsilon_i, \varepsilon_{p,i}) & Var(\varepsilon_{p,i}) \end{pmatrix}$, $\mathbf{V}_{z_{int}} = \begin{pmatrix} 1 & 0_{|1\times z_{int}|} \\ 0_{|z_{int}\times 1|} & Var(z_{int,i}) \end{pmatrix}$, and $\bigotimes$ denotes Kronecker product.

Next we define the first component of $\Psi$ from 24. Note that since $g_i = g_i^c + q_i$,

$$Var(\boldsymbol{Q}_i) + Cov(\boldsymbol{G}_i^c, \boldsymbol{Q}_i) = \begin{pmatrix} Cov(g_i, q_i) & Cov(g_i, q_i \mathbf{z}_{int,i}) & Cov(g_i, q_{p,i}) & Cov(g_i, q_{p,i} \mathbf{z}_{int,i}) \\ Cov(g_i \mathbf{z}_{int,i}, q_i) & Cov(g_i \mathbf{z}_{int,i}, q_i \mathbf{z}_{int,i}) & Cov(g_i \mathbf{z}_{int,i}, q_{p,i}) & Cov(g_i \mathbf{z}_{int,i}, q_{p,i} \mathbf{z}_{int,i}) \\ Cov(g_{p,i}, q_i) & Cov(g_{p,i}, q_i \mathbf{z}_{int,i}) & Cov(g_{p,i}, q_{p,i}) & Cov(g_{p,i}, q_{p,i} \mathbf{z}_{int,i}) \\ Cov(g_{p,i} \mathbf{z}_{int,i}, q_i) & Cov(g_{p,i} \mathbf{z}_{int,i}, q_i \mathbf{z}_{int,i}) & Cov(g_{p,i} \mathbf{z}_{int,i}, q_{p,i}) & Cov(g_{p,i} \mathbf{z}_{int,i}, q_{p,i} \mathbf{z}_{int,i}) \end{pmatrix} \tag{27}$$

Next, we define $\alpha$ and $Var(\varepsilon)$.Begin by defining $\psi = \frac{sd(\mathbf{x}_i \boldsymbol{\gamma})}{sd(\mathbf{x}_i \boldsymbol{\mu})}$. Thus,

$$\alpha^2 = Var(g_i^c + q_i + \varepsilon_i) = Var\Big( \frac{\mathbf{x}_i \boldsymbol{\mu}}{sd(\mathbf{x}_i \boldsymbol{\gamma})} + \varepsilon_i \Big) = \frac{1}{\psi^2} + Var(\varepsilon_i). \tag{28}$$

Now note that

$$h_{SNP}^2 = \frac{Cov(y_i, g_i)^2}{Var(y_i)Var(g_i)} \tag{29}$$

$$R^2 = \frac{Cov(y_i, \hat{g}_i)^2}{Var(y_i)Var(\hat{g}_i)} = \frac{Cov(y_i, \frac{g_i+\varepsilon_i}{\alpha})^2}{Var(y_i)} = \frac{Cov(y_i, g_i)^2}{\alpha^2 Var(y_i)} = \frac{Var(g_i)h_{SNP}^2}{\alpha^2}. \tag{30}$$

This implies that

$$\alpha^2 = \frac{Var(g_i)h_{SNP}^2}{R^2} = \frac{Var(\mathbf{x}_i\boldsymbol{\mu})}{Var(\mathbf{x}_i\boldsymbol{\gamma})}\frac{h_{SNP}^2}{R^2} = \frac{\rho^2}{\psi^2} \geq 1, \tag{31}$$

where $\rho^2 = \frac{h_{SNP}^2}{R^2}$. Hence, $Var(\varepsilon_i) = Var(\varepsilon_{p,i}) = \frac{1}{\psi^2}(\rho^2 - 1)$.

Now let us define $Cov(\varepsilon_i, \varepsilon_{p,i})$. Note that

$$\begin{gathered} Cov(\hat{g}_i, \hat{g}_{p,i}) = Cov(\tfrac{g_i}{\alpha}, \tfrac{g_{p,i}}{\alpha}) + Cov(\tfrac{\varepsilon_i}{\alpha}, \tfrac{\varepsilon_{p,i}}{\alpha}) = \\ \tfrac{1}{\alpha^2}\Big(Cov(\tfrac{sd(\mathbf{x}_i\boldsymbol{\mu})\mathbf{x}_i\boldsymbol{\mu}}{sd(\mathbf{x}_i\boldsymbol{\gamma})sd(\mathbf{x}_i\boldsymbol{\mu})}, \tfrac{sd(\mathbf{x}_{p,i}\boldsymbol{\mu})\mathbf{x}_{p,i}\boldsymbol{\mu}}{sd(\mathbf{x}_{p,i}\boldsymbol{\gamma})sd(\mathbf{x}_{p,i}\boldsymbol{\mu})}) + Cov(\varepsilon_i, \varepsilon_{p,i})\Big) = \\ \tfrac{\psi^2}{\rho^2}\Big(\tfrac{\rho_{po}}{\psi^2} + Cov(\varepsilon_i, \varepsilon_{p,i})\Big). \end{gathered} \tag{32}$$

This implies that

$$\begin{gathered} Cov(\varepsilon_i, \varepsilon_{p,i}) = \tfrac{\rho^2}{\psi^2}Cov(\hat{g}_i, \hat{g}_{p,i}) - \tfrac{1}{\psi^2}\rho_{po} = \\ \tfrac{1}{\psi^2}\Big(Cov(\hat{g}_i, \hat{g}_{p,i})\rho^2 - \rho_{po}\Big), \end{gathered} \tag{33}$$

where $\rho_{po}$ is a parent-offspring genetic correlation. This implies that

$$Var(\varepsilon) = \frac{1}{\psi^2}\begin{pmatrix} \rho^2 - 1 & Cov(\hat{g}_i, \hat{g}_{p,i})\rho^2 - \rho_{po} \\ Cov(\hat{g}_i, \hat{g}_{p,i})\rho^2 - \rho_{po} & \rho^2 - 1 \end{pmatrix} = \frac{1}{\psi^2}\boldsymbol{\Omega}\ . \tag{34}$$

Note that under random mating and when parental genotypes are not imputed, $Cov(\hat{g}_i, \hat{g}_{p,i}) = \rho_{po} = \frac{1}{\sqrt{2}}$, implying that $Cov(\varepsilon_i, \varepsilon_{p,i}) = \frac{1}{\psi^2}\frac{1}{\sqrt{2}}(\rho^2 - 1)$.

Therefore, having information of $\Psi$, $\psi$, and $\rho$ allows us to correct the bias in estimates of $\boldsymbol{\beta}_{\hat{G}}$, where the vector of corrected estimates is

$$\hat{\boldsymbol{\beta}}_{corr} = \mathbf{A}^{-1}\hat{\boldsymbol{\beta}}_{\hat{G}}, \tag{35}$$

where $\mathbf{A} = \Big(\mathbf{V}_{\hat{g}}^{-1}\mathbf{P}^{-1}(\mathbf{P}\mathbf{V}_{\hat{g}}\mathbf{P} - \mathbf{V}_E - \Psi)\Big)$. And the standard errors can be obtained from

$$Var(\hat{\boldsymbol{\beta}}_{corr}) = \mathbf{A}^{-1}Var(\hat{\boldsymbol{\beta}}_{\hat{G}})(\mathbf{A}^{-1})^T. \tag{36}$$

However, in this general case, it is typically infeasible to implement the bias correction. That is mainly because $\Psi$ is unobserved in most cases because the non-classical measurement error may not be independent of the covariates $\mathbf{z}_i$, and the variance of $\mathbf{z}_{int,i}$ may depend on $g_i$ and $q_i$, making it difficult to infer $Cov(g_i\mathbf{z}_{int,i}, q_i\mathbf{z}_{int,i})$. Also note that in the special case $\boldsymbol{\gamma} = \boldsymbol{\mu}$, then $\mathbf{A} = \Big(\mathbf{V}_{\hat{g}}^{-1}\mathbf{P}^{-1}(\mathbf{P}\mathbf{V}_{\hat{g}}\mathbf{P} - \mathbf{V}_E)\Big)$, which is the correction equation from[12].

Hence, we make the following assumptions:

**Assumption** 1. Random mating.

**Assumption** 2. The control variables that are interacted with the PGI, $\mathbf{z}_{int,i}$, are independent of the individual and parental genotypes, $\mathbf{x}_i$ and $\mathbf{x}_{p,i}$.

#### 1.3 Bias derivation for two special cases

Here we specialize the bias formula derived above for two special cases: (1) the control variables $\mathbf{z}_i$ are uncorrelated with $\mathbf{x}_i$ and $\mathbf{x}_{p,i}$, and (2) the control variables $\mathbf{z}_i$ are not causally affected by the genotype vector $\mathbf{x}_i$. The first special case is simpler, and the second corresponds to the analysis described in the main text.

We begin with some more general observations. Under Assumption 1 (random mating), $Cov(\hat{g}_i, \hat{g}_{p,i}) = \rho_{po} = \frac{1}{\sqrt{2}}$. (Note that this might not be the case when parental genes are imputed even in the absence of assortative mating since imputation removes some of the parental genetic variation that is uncorrelated with the offspring genetic variation.)

Under both Assumptions 1 and 2 described above,

$$\mathbf{\Psi} = \mathbf{V}_q + \mathbb{M}'_{q,g^c} = \begin{pmatrix} Var(\boldsymbol{Q}_i) + Cov(\boldsymbol{G}_i^c, \boldsymbol{Q}_i) & 0_{|\mathbb{G}\times z|} \\ Cov(\mathbf{z}_i, \boldsymbol{Q}_i) & 0_{|z\times z|} \end{pmatrix} = \begin{pmatrix} \Big(Var(\mathbf{q}) + Cov(\mathbf{g}^c, \mathbf{q})\Big) \bigotimes \mathbf{V}_{z_{int}} & 0_{|\mathbb{G}\times z|} \\ Cov(\mathbf{z}_i, \boldsymbol{Q}_i) & 0_{|z\times z|} \end{pmatrix} \tag{37}$$

and $Var(\mathbf{q}) + Cov(\mathbf{g^c}, \mathbf{q}) = \Big(\frac{Var(\mathbf{x}_i\boldsymbol{\theta})}{Var(\mathbf{x}_i\boldsymbol{\gamma})} + \frac{Cov(\mathbf{x}_i\boldsymbol{\gamma}, \mathbf{x}_i\boldsymbol{\theta})}{Var(\mathbf{x}_i\boldsymbol{\gamma})}\Big)\mathbf{\Gamma}$ , where $\mathbf{\Gamma} = \begin{pmatrix} 1 & \frac{1}{\sqrt{2}} \\ \frac{1}{\sqrt{2}} & 1 \end{pmatrix}$.

Also note that

$$\frac{Cov(\mathbf{x}_i\boldsymbol{\gamma}, \mathbf{x}_i\boldsymbol{\theta})}{Var(\mathbf{x}_i\boldsymbol{\gamma})} + \frac{Var(\mathbf{x}_i\boldsymbol{\theta})}{Var(\mathbf{x}_i\boldsymbol{\gamma})} = \frac{Cov(\mathbf{x}_i\boldsymbol{\mu}, \mathbf{x}_i\boldsymbol{\mu} - \mathbf{x}_i\boldsymbol{\gamma})}{Var(\mathbf{x}_i\boldsymbol{\gamma})} = \\ \frac{Var(\mathbf{x}_i\boldsymbol{\mu})}{Var(\mathbf{x}_i\boldsymbol{\gamma})} - \frac{Cov(\mathbf{x}_i\boldsymbol{\gamma}, \mathbf{x}_i\boldsymbol{\mu})}{Var(\mathbf{x}_i\boldsymbol{\gamma})} = \frac{Var(\mathbf{x}_i\boldsymbol{\mu})}{Var(\mathbf{x}_i\boldsymbol{\gamma})} - \frac{Corr(\mathbf{x}_i\boldsymbol{\gamma}, \mathbf{x}_i\boldsymbol{\mu})sd(\mathbf{x}_i\boldsymbol{\mu})}{sd(\mathbf{x}_i\boldsymbol{\gamma})} = \frac{1 - \psi Corr(\mathbf{x}_i\boldsymbol{\gamma}, \mathbf{x}_i\boldsymbol{\mu})}{\psi^2} = \frac{1 - \psi r}{\psi^2}, \tag{38}$$

where $r = Corr(\mathbf{x}_i\boldsymbol{\gamma}, \mathbf{x}_i\boldsymbol{\mu})$, the correlation between the associative and causal genetic effect.

Note that under the assumption of random mating, $Var(\epsilon) = \frac{1}{\psi^2}\mathbf{\Omega} = \frac{1}{\psi^2}(\rho^2 - 1)\mathbf{\Gamma}$.

**Case 1**: Covariates are uncorrelated with $\mathbf{x}_i$ and $\mathbf{x}_{p,i}$.
Note this implies that $Cov(\mathbf{z}_i, \boldsymbol{Q}_i) = 0_{|z\times Q|}$. Thus,

$$Var(\hat{\boldsymbol{G}}_i) = \begin{pmatrix} \mathbf{\Gamma} \bigotimes \mathbf{V}_{z_{int}} & 0_{|\mathbb{G}\times z|} \\ 0_{|z\times\mathbb{G}|} & Var(z_i) \end{pmatrix},$$

$$V_E = \frac{1}{\psi^2}(\rho^2 - 1) \begin{pmatrix} \mathbf{\Gamma} \bigotimes \mathbf{V}_{z_{int}} & 0_{|\mathbb{G}\times z|} \\ 0_{|z\times\mathbb{G}|} & 0_{|z|}, \end{pmatrix},$$

$$\Psi = \frac{1}{\psi^2}\Big(1 - \psi r\Big) \begin{pmatrix} \mathbf{\Gamma} \bigotimes \mathbf{V}_{z_{int}} & 0_{|\mathbb{G}\times z|} \\ 0_{|z\times\mathbb{G}|} & 0_{|z|} \end{pmatrix}.$$

Substituting this into (35), we obtain:

$$\boldsymbol{\beta}_{\hat{G}} = \begin{pmatrix} diag(\frac{r}{\rho}) & 0_{|\mathbb{G}\times z|} \\ 0_{|z\times\mathbb{G}|} & I_{|z\times z|} \end{pmatrix} \boldsymbol{\beta}_{G^c}, \tag{39}$$

where $diag(\frac{r}{\rho})$ is a diagonal matrix of size $\mathbb{G} \times \mathbb{G}$ with $\frac{r}{\rho}$ on its main diagonal. Hence, when the controls are uncorrelated with $\mathbf{x}_i$ and $\mathbf{x}_{p,i}$, $\boldsymbol{\beta}_{\hat{G}} = \frac{r}{\rho}\boldsymbol{\beta}_{G^c}$ for both PGIs and their interactions, and $\boldsymbol{\beta}_{\hat{G}} = \boldsymbol{\beta}_{G^c}$ for the control variables.

Therefore, we can compute the corrected estimate $\hat{\boldsymbol{\beta}}_{corr} = \frac{\rho}{r}\hat{\boldsymbol{\beta}}_{\hat{G}}$ for both PGIs and their interactions and $\hat{\boldsymbol{\beta}}_{corr} = \hat{\boldsymbol{\beta}}_{\hat{G}}$ for the control variables. Note that we can compute $SE(\hat{\boldsymbol{\beta}}_{corr})$ by taking the square root of the diagonal elements of matrix $Var(\hat{\boldsymbol{\beta}}_{corr})$ computed according to equation 36, with $\mathbf{A} = \begin{pmatrix} diag(\frac{r}{\rho}) & 0_{|\mathbb{G}\times z|} \\ 0_{|z\times\mathbb{G}|} & I_{|z\times z|} \end{pmatrix}$.

Given that $\mathbf{A}$ is diagonal, it is straightforward that $SE(\hat{\boldsymbol{\beta}}_{corr}) = \frac{\rho}{r}SE(\hat{\boldsymbol{\beta}}_{\hat{G}})$ for both PGIs and their interactions, and $SE(\hat{\boldsymbol{\beta}}_{corr}) = SE(\hat{\boldsymbol{\beta}}_{\hat{G}})$ for the control variables.

**Case 2**: The control variables $\mathbf{z}_i$ are not causally affected by the genotype vector $\mathbf{x}_i$.

In this case, the correlation between the control variables $\mathbf{z}_i$ and the transmitted alleles is equal to the correlation between $\mathbf{z}_i$ and the non-transmitted alleles. This implies that: $Cov(\mathbf{z}_i, q_{p,i}) = \sqrt{2}Cov(\mathbf{z}_i, q_i)$ and $Cov(\mathbf{z}_i, \hat{g}_{p,i}) = \sqrt{2}Cov(\mathbf{z}_i, \hat{g}_i)$.

First, suppose that interactions are not included in the model. It can be shown that in this case, the estimated child's PGI effect satisfies

$$\beta = \frac{r}{\rho}\beta_c,$$

while the bias in the parental coefficient $\beta_p$ is more complex and depends on the correlation with the control variables. Notably, this involves computing $Var(\hat{\boldsymbol{G}}_i)^{-1}$.

Since we are demonstrating only the magnitude of the bias in the child's PGI effect $\beta_c$, it suffices to compute the first row of $Var(\hat{\boldsymbol{G}}_i)^{-1}$. To compute the inverse, we use the adjugate method:

$$Var(\hat{\mathbf{G}})^{-1}_{i,j} = (-1)^{i+j}\frac{M_{i,j}}{\det(Var(\hat{\mathbf{G}}))},$$

where $Var(\hat{\mathbf{G}})^{-1}_{i,j}$ is the $(i,j)$ entry of $Var(\hat{\mathbf{G}})^{-1}$, and $M_{i,j}$ is the minor of the $(i,j)$ entry of $Var(\hat{\mathbf{G}})$.

Now, the key observation is that if we delete the first row of $Var(\hat{\mathbf{G}})$ to compute the minors of the elements in the first row, columns 1 and 2 are identical up to a constant factor (column 2 is $\sqrt{2}\times$ column 1 under the imposed assumptions). This implies that

$$M_{1,z_1} = \cdots = M_{1,z_N} = 0,$$

and consequently, all elements in the last $z$ columns of row 1 are equal to zero.

Next, we determine the elements $Var(\hat{\mathbf{G}})^{-1}_{1,1}$ and $Var(\hat{\mathbf{G}})^{-1}_{1,2}$. Since all but the first two elements of the first row are zero, the Laplace expansion yields

$$Var(\hat{\mathbf{G}})_{1,1}Var(\hat{\mathbf{G}})^{-1}_{1,1} + Var(\hat{\mathbf{G}})_{1,2}Var(\hat{\mathbf{G}})^{-1}_{1,2} = Var(\hat{\mathbf{G}})^{-1}_{1,1} + \frac{1}{\sqrt{2}}Var(\hat{\mathbf{G}})^{-1}_{1,2} = 1.$$

Additionally, since deleting the first row of $Var(\hat{\mathbf{G}})$ results in column 2 being $\sqrt{2}\times$ column 1, we have

$$M_{1,2} = \sqrt{2}M_{1,1},$$

which implies that

$$Var(\hat{\mathbf{G}})^{-1}_{1,2} = -\sqrt{2}Var(\hat{\mathbf{G}})^{-1}_{1,1}.$$

Solving this system for $Var(\hat{\mathbf{G}})^{-1}_{1,1}$ and $Var(\hat{\mathbf{G}})^{-1}_{1,2}$, we obtain

$$Var(\hat{\mathbf{G}})^{-1}_{1,1} = 2, \quad Var(\hat{\mathbf{G}})^{-1}_{1,2} = -\sqrt{2}.$$

Thus, the first row of $Var(\hat{\mathbf{G}})^{-1}$ is

$$[2, -\sqrt{2}, 0, \ldots, 0].$$

The remainder of the proof straightforward and shows that the first row of the correction matrix $\mathbf{A}$ is

$$\left[\frac{r}{\rho}, 0, \ldots, 0\right],$$

which implies that

$$\beta = \frac{r}{\rho}\beta_c.$$

In a similar manner, we can show that under our assumptions, when the interactions between $\hat{g}_i$ and $\hat{g}_{p,i}$ with $\mathbf{z}_{int,i}$ are included, the first $1+z_{int}$ rows (corresponding to $\hat{g}_i$ and $\hat{g}_i\mathbf{z}_{int,i}$) of $Var(\hat{\mathbf{G}})^{-1}$ are $[2, 0, -\sqrt{2}, 0, .., 0]$

and $[0, 2, 0, -\sqrt{2}, 0, .., 0]$. This implies that the corresponding rows of the correction matrix $\mathbf{A}$ are $[\frac{r}{\rho}, 0, .., 0]$ and $[0, \frac{r}{\rho}, 0, .., 0]$. This, in turn, implies that

$$\beta = \frac{r}{\rho}\beta_c$$

and

$$\boldsymbol{\delta} = \frac{r}{\rho}\boldsymbol{\delta}_c.$$

Hence both the child's PGI effect and its interactions with $\mathbf{z}_{int,i}$ will be attenuated by a factor of $\frac{r}{\rho}$. This is the result reported in the main text.

Therefore, we can compute the vector of corrected estimates $\hat{\boldsymbol{\beta}}_{corr} = \frac{\rho}{r}\hat{\boldsymbol{\beta}}_{\hat{G}}$ for the individuals own PGI effect and its interactions with $\mathbf{z}_{int}$. We can compute $SE(\hat{\boldsymbol{\beta}}_{corr})$ by taking the square root of the diagonal elements of matrix $Var(\hat{\boldsymbol{\beta}}_{corr})$ computed according to equation 36. Note that because the first $1 + z_{int}$ rows of the correction matrix $\mathbf{A}$ have $\frac{r}{\rho}$ in the main diagonal and the rest of elements in these rows are equal to zero, the first $1 + z_{int}$ rows of $\mathbf{A}^{-1}$ have $\frac{\rho}{r}$ in the main diagonal and the rest of elements in these rows are equal to zero. This can be shown using the Laplace expansion and the adjugate method to compute the elements of $\mathbf{A}^{-1}$. Hence, equation 36 implies that the first $1 + z_{int}$ diagonal elements of $Var(\hat{\boldsymbol{\beta}}_{corr})$ are the first $1 + z_{int}$ diagonal elements of $Var(\hat{\boldsymbol{\beta}}_{\hat{G}})$ multiplied by $\frac{\rho^2}{r^2}$, which implies that $SE(\hat{\boldsymbol{\beta}}_{corr}) = \frac{\rho}{r}SE(\hat{\boldsymbol{\beta}}_{\hat{G}})$ for child's own genetic effect and its interactions with the independent controls $\mathbf{z}_{int,i}$. Note that the correction of parental genetic effect and its interactions with $\mathbf{z}_{int,i}$ is more complex and requires additional information.

#### 1.4 Assortative mating

A GWAS coefficient is estimated by regressing $y_i$ on each genetic variant $x_{ij}$. Thus, the GWAS coefficient for variant $j$ is $\hat{\boldsymbol{\mu}}_j = \boldsymbol{\gamma}_j + \boldsymbol{\theta}_j(1 + Cov(x_{ij}, x_{n,ij})) + \varepsilon_j$. This suggests that when there is assortative mating, PGIs cannot be expressed as described in equation (6).

To analyse how the bias changes when there is assortative mating and how this bias is comparable to our theoretically derived bias, we conduct a simple simulation under random and assortative mating.

We start by generating genomes of 2,000 biallelic, independent SNPs for two samples of size $M \in \{15000, 24286\}$ individuals in the initial generation, assuming half are male and half are female. For the random mating simulation, we randomly match each male with a female and simulate two offspring per pair under the laws of Mendelian segregation assuming each SNP is inherited independently. We then generate a phenotype with variance one according to Equation (1), where the effect sizes are drawn from mean-zero normal distribution with variance such that the narrow-sense heritability is 0.2, the contribution of the parental component to variance is 0.1, and the correlation of these two vectors of coefficients (which we refer to as the "Child-parental effect correlation") varies between zero and one.

For the assortive mating simulation, we add standard normal noise to the phenotype and match the males and females according to their rank of the noisy phenotype such that the correlation of the mates for the original phenotype is 0.5. Then, we simulate two offspring for each pair and calculate their phenotypes using the same simulated self and and parental effect sizes as in the randomly mating generation. This is repeated 100 times such that population is in equilibrium. In the final generation, we do not add noise to the phenotype.

Next, for both simulations, we conduct a GWAS (not controlling for the parental genotypes) in a sample of either 5,000 individulals (high $\rho$) or 14,286 indivduals (low $\rho$). These sample sizes were chosen because, under the data generation procedure describe above, $N = 5,000$ corresponds to $\rho = 1.7$ and $N = 14,286$ corresponds to $\rho = 3$ in a randomly mating population where the child-parental effect correlation is zero. We use the GWAS coefficients as SNP weights to build PGIs for the parents and the offspring in the residual sample of 10,000 individuals. We then regress the PGI onto the phenotype in the prediction sample and report the coefficient from this regression.

Figure 1 presents the results from 10 replications of this simulation. The black markers represent the true (i.e., observed) mean coefficient from the PGI regression for the Low $\rho$ and High $\rho$ setting as well as the 95% confidence intervals. The gray markers represent the theoretical expected attenuation derived above under a model of random mating. If the GWAS sample size were infinite and there were no bias from the parental effects, the estimated effect would be one. However, because sample sizes are finite, introducing sampling error, and there is confounding from parental effects, the estimates are attenuated. In Panel (a), we see that using data from our random-mating simulation, the theoretical attenuation is contained within the confidence interval of the observed attenuation. However, in our assortative-mating simulation in Panel (b), the attenuation is larger than predicted by our theoretical model, although the difference is small.

Figure 1: Attenuation factor of the individual's PGI effect. Simulation under random and assortative mating

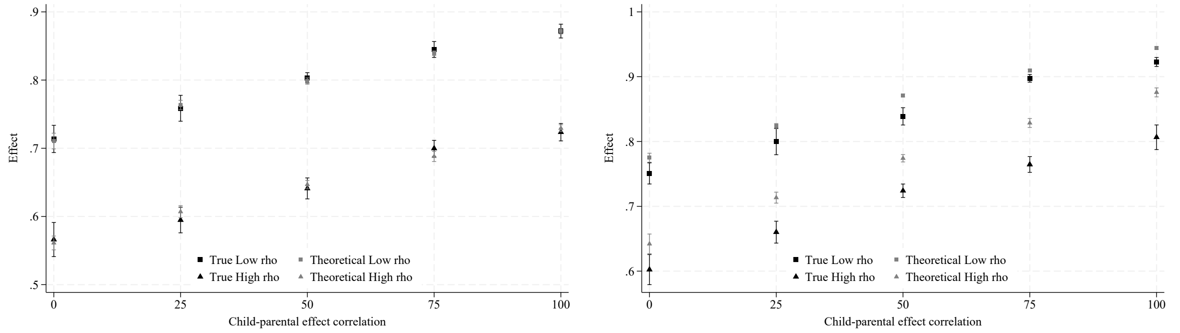

(a) Random Mating

(b) Assortative Mating (50%)

The average values of the observed and theoretical attenuation factors, along with the 95% confidence intervals from 10 simulations for each set of parameters, are reported. The theoretical attenuation factor is calculated as $Corr(assoc., causal)/\rho$.

### 2 Polygenic Index Repository User Guide

In this guide, we summarize the key information regarding the construction of the Repository PGIs, lay out some of the interpretational issues that are likely to arise as researchers begin to use PGIs from the Repository, and outline how we suggest thinking through those issues. This version of the User Guide is up to date as of the publication of ref.[13]; the most up-to-date version is available on the SSGAC website: `https://www.thessgac.org/pgi-repository`.

#### 2.1 Summary information about Repository PGIs

Here, we provide a brief summary of how the PGIs were constructed (please see Methods for a more detailed description). We refer the reader to the relevant tables where more information can be found.

##### 2.1.1 Phenotype definitions and GWAS

The PGIs are based on meta-analyses of summary statistics from up to three sources: GWAS conducted in 23andMe, Inc. and UKB (some of which are novel), and published GWAS. Supplementary Table 2 lists the phenotype measures used in the new or updated UKB GWAS that we conducted ourselves, including information on how repeated measures were handled and the sample size in each of the three UKB partitions. Supplementary Table 7 lists the phenotype definitions and describes the association models for all novel or published 23andMe GWAS, and for published GWAS, it cites the relevant publications. For phenotypes included in the first release of the Repository whose UKB or 23andMe GWAS were not updated in the current release, the corresponding information can be found in Supplementary Tables 5 and 6 in Becker et al.[10].

In order to avoid sample overlap between the GWAS and Repository datasets, we conducted multiple versions of the GWAS meta-analysis for each phenotype (so as to have, for each dataset, a version of the meta-analysis that excludes that dataset). Supplementary Table 11 lists all GWAS meta-analyses used as inputs for the

PGIs. The "Repository datasets meta-analysis is used to make PGIs for" column shows which meta-analysis the SNP weights come from for each Repository dataset.

###### 2.1.2 PGI construction

The PGIs were made using SBayesR[8] applied to the overlapping variants between each input GWAS meta-analysis and 2,865,810 pruned common variants from the full UKB European-genetic-ancestry dataset for which LD estimates were made available by Lloyd-Jones et al.[8]. The inclusion criterion was that the "expected" out-of-sample predictive power of a PGI be greater than 1%. The expected predictive power was calculated from the results of the largest GWAS meta-analysis available for that phenotype[15]. The expected predictive power of each PGI (including the ones not included in the Repository because they did not pass the cutoff of 1%) are shown in Supplementary Table 11. Notably, even though the *expected* predictive power of each PGI is greater than 1%, in many instances, the *actual* predictive power of the PGI in a particular dataset may be less than 1%.

###### 2.1.3 PC construction

As part of the Repository, we also release 20 principal components (PCs) based on the genome-wide data in each of the participating cohorts. The primary purpose of the release is to make them available for users who wish to use them as controls for population stratification. In order to make the PCs, we first restricted the samples to individuals of European genetic ancestries and removed markers with imputation accuracy less than 70% or minor allele frequency less than 1%, as well as markers in long-range LD blocks (provided by the plinkQC R package[16]). We then pruned all SNPs that survived these filters using a 1Mb rolling window (incremented in steps of 5 variants) and an $r^2$ threshold of 0.1. Next, we calculated the pairwise relatedness between all individuals in our full sample and generated a sample of conventionally unrelated individuals by dropping one individual from each pair of individuals with an estimated relatedness greater than 0.05 as calculated by in Plink1.9[17]. We then estimated the first 20 PC loadings in this sample of approximately unrelated individuals. Finally, we projected all individuals in the sample— including both members of related pairs—onto these loadings to compute their corresponding PCs.
In HRS, we re-labeled the PCs in sets of five in order to address identifiability concerns. Therefore, it is only possible to infer from the variable name of a PC if it is one of the first five PCs (PC 1-5), one of the next five PC (PCs 6-10), etc.

###### 2.1.4 Genotyping, imputation, and phenotype definitions in Repository datasets

Details on genotyping and imputation of the Repository datasets are listed in Supplementary Table 6. Supplementary Table 14 lists the phenotype definitions for the subset of these datasets that we used to validate our PGIs, excluding UKB. The phenotype definitions for UKB can be found in Supplementary Table 2.

###### 2.1.5 Predictive power of Repository PGIs in validation datasets

Supplementary Table 3 shows the observed predictive power of the Repository PGIs in our three validation datasets, together with 95% confidence intervals obtained using a bootstrap with 1000 repetitions. For phenotypes that were included in the first release of the Repository, the table also shows the predictive power of the first release PGIs for comparison.

##### 2.2 Interpretational considerations

In this section, we lay out some of the interpretational issues that are likely to arise as researchers begin to use PGIs from the Repository, and we outline how we suggest thinking through those issues. The executive summary is as follows:

1. The methodologies used to conduct the GWAS and to construct the SNP weights jointly determine the additive SNP factor that is proxied for by the PGI.

2. These methodologies, together with the PGI phenotype, determine the relative importance of various potential confounds to a causal interpretation of PGI associations. In most applications, researchers should control for PCs (which are available from the datasets, along with the PGIs, as part of the Repository).

3. Whether and which confounds should be highlighted (or can be safely ignored) depends on the application.

4. Currently, the best way to cleanly identify causal effects is to conduct a family-based PGI study (where the analysis controls for the parental PGI, constructed from either measured or imputed parental genotypes). While the results of such a study have a causal interpretation, the correct interpretation is subtle, and the results will generally underestimate the causal effects of genetic variants. In the absence of clean identification of causal effects, researchers should highlight the potential confounds to a causal interpretation.

5. In interpreting PGI associations (whether causal or not), it is important to keep in mind that genetic effects can operate through environmental mechanisms, and these mechanisms may be modifiable. For this reason, researchers should be cautious about using terminology such as "genetic endowment" that can connote genetic determinism. Researchers should remind readers of the potential role of environmental mechanisms in explaining PGI associations.

The following subsections, numbered 1 through 5, provide more detail on the points above. In addition to attending to these interpretational issues, we urge users of the Repository to conduct power calculations prior to undertaking analyses; to pursue analyses only if they are adequately powered; and, when feasible, to preregister planned analyses (along with the power calculations).

We note that the GWAS from which the Repository PGIs are constructed were conducted in samples of European genetic ancestries (where "European genetic ancestry" is operationalized differently depending on the study but almost always involves sample restrictions based on the genetic PCs; e.g., for our UKB GWAS, see the "UKB GWAS" subsection of Section II in Methods). Due to the limited portability of such GWAS results to other ancestries, for the PGIs released to participating datasets, the current version of the Repository is restricted to individuals of European genetic ancestries, as defined by how their genetic PCs cluster together with those classified as having EUR-genetic-ancestry in the 1000 Genomes Project (see the "Subject-level QC" subsection of Section I in Methods).

##### 2.2.1 GWAS and SNP-Weight Methodologies and the Additive SNP Factor

In the Supplementary Methods section 6 of Becker et al.[10], we showed how the set of control variables used in a GWAS affects the additive SNP factor proxied for by a PGI. The choice of controls, however, is just one of many dimensions of GWAS methodology. A change to any of these dimensions is likely to result in a different additive SNP factor (with a different interpretation). For example, it is increasingly common for researchers to conduct association analyses using mixed-linear models rather than OLS[18,19]. Since mixed-linear models often produce estimates that are more robust to stratification, the additive SNP factor will be akin to that generated by an OLS-based GWAS with some additional controls for stratification. Knowledge of the methodology of the GWAS underlying a particular PGI is therefore often a necessary first step for understanding what additive SNP factor a specific PGI is proxying for. For example, the methodologies underlying the GWASs we conducted in UKB for the PGIs in the Repository are described in the "UKB GWAS" subsection of Section II in Methods. Information about the association models in the 23andMe GWASs can be found in Supplementary Table 6 of Becker et al. (2021)[10] and Supplementary Table 7 of the current paper.

The SNP-weight methodology can matter, as well. For example, our Repository PGI SNP-weights are calculated from the GWAS results using $\sim 2.9$ million pruned common variants from the full UKB European-genetic-ancestry ($N \approx 450{,}000$) data set from Lloyd-Jones et al.[8], which primarily capture common genetic variation. If SNP weights were instead calculated based on results from SNPs that capture a different mix of common and rare genetic variation, then the additive SNP factor corresponding to that PGI would have a different interpretation: it would be the best linear predictor based on that set of SNPs.

##### 2.2.2 Potential Confounds to a Causal Interpretation

It is increasingly understood that standard GWAS approaches with a limited set of controls – for example, sex, age, and up to 10 PCs, as in most of the GWASs underlying the Repository PGIs – generate PGIs that can be subject to a number of confounds to a causal interpretation[20–23]. For example, PGIs for educational attainment derive a substantial share of their overall predictive power from their positive association with rearing environment. In behavior-genetic parlance, this positive correlation arises due to the vertical transmission of the parental phenotypes (parents' phenotypes impact their children's phenotypes). In recent molecular-genetic research, this source of positive gene-environment correlation has been labeled "genetic nurture"[21] or "parental indirect effects"; we refer to them as "parental genetic effects." These effects can be further exacerbated by assortative mating at the genetic level.

As another example, when the PCs are estimated in a small sample, they are often not very accurate proxies for ancestry. Failure to adequately control for genetic ancestry gives rise to "population stratification"[24]: because the PGI is correlated with ancestry, which in turn is correlated with ethnicity and regional background, it picks up cultural or environmental factors that are correlated with these factors. In many empirical applications, the goal is to estimate an association that is net of any such cultural and environmental confounds. In such cases, it may be possible to mitigate concerns that the underlying GWAS may have relied on inaccurate ancestry controls by including a richer-than-usual set of environmental controls in the analysis of the PGI.

Indeed, in most applications (that cannot exploit family data to control for parental PGIs), researchers should include PCs in the set of controls. When estimating PGI-by-environment interactions, researchers should additionally control for interactions between PCs and the "environment" variable[25]. For these purposes, dataset-specific PCs are made available as part of the Repository. However, it is important to recognize and acknowledge that the PCs are not fully accurate measures of ancestry, so even after controlling for PCs, residual confounding almost surely remains.

The relevance of potential confounds could vary across phenotypes[20,22,23]. For example, parental genetic effects are much smaller for height than educational attainment. Although the noisiness of PCs as measures of ancestry in a given sample is the same across phenotypes, the noisiness is likely to be substantially more problematic for educational attainment than for height because finer ancestral distinctions (which require more PCs to capture) probably matter for the social and environmental factors that influence educational attainment. More generally, it seems likely that potential confounds to a causal interpretation matter more for PGIs for social and behavioral phenotypes than for PGIs for more biologically proximal phenotypes.

##### 2.2.3 Importance of Confounds Depends On the Application

The degree to which potential confounds to a causal interpretation matter depends on how the PGI is used. For example, if a PGI is used as a control variable to increase precision for a randomized treatment evaluation[26,27], then the goal is simply to use controls that absorb as much residual variance as possible (and avoid controlling for any variables realized after the randomized intervention). Since the PGI is simply being used as a predictive variable, its interpretation is irrelevant in that case. As a contrasting example, consider the illustrative application in Becker et al.[10] that tests how much parental education mediates the predictive power of the PGI for educational attainment. There, the PGI should be understood as capturing some of the parental genetic effects and ancestry associations with education. In most applications, the potential confounds do matter and should be highlighted.

##### 2.2.4 Identifying Causal Effects of Genetic Variants Using a Family-Based PGI Study

The cleanest way to identify the causal effects of a PGI is to control for the parental PGI (which may be constructed from either parental genotypes that are directly measured or imputed from other genotyped family data, such as sibling data). This empirical strategy exploits a natural experiment: conditional on a pair of biological parents, genetic inheritance is random. A robustly estimated non-zero estimate in a family-based PGI study from a large and attrition-free sample would provide strong evidence of causal effects of genetic variants. However, the causal interpretation of this estimate is nuanced because there is no single, well-defined thought experiment that corresponds to changing the value of the PGI, since changing different genotypes that have different effects could generate the same change in the PGI. The correct interpretation

of the coefficient estimate is a weighted average of treatment effects from hypothetical experiments that randomly modify, at conception, the genotypes of the causal SNPs responsible for the predictive power of the PGI[1,2].
The additive SNP factors corresponding to the PGIs in the Repository are not the best linear predictors conditional on a pair of biological parents (because the GWAS underlying the SNP weights do not control for the biological parents' genotypes). The PGIs proxying for additive SNP factors that would be the best linear predictors for such a family-based analysis would be PGIs constructed from GWAS that control for parental genotypes (or from GWAS in sibling samples that control for family fixed effects, although these GWAS estimates would be biased if siblings' genotypes have causal effects on an individual's phenotype). Unfortunately, to date genotyped family-based samples have been too small to produce reliable "within-family PGIs." The Repository does not yet contain any such PGIs. Ultimately, however, when genotyped family-based samples become sufficiently large, the resulting within-family PGIs will be more predictive for family-based analyses than PGIs constructed from currently-standard (between-family) GWAS.

##### 2.2.5 Genetic Effects Can Operate Through Environmental Mechanisms

We urge researchers who use PGIs in their research to be mindful of three important issues of interpretation for the causal effects of a PGI. First, a PGI could exert its effects through the environment[28]. Consider a PGI for BMI[26]. Suppose a family-based association analysis yields unambiguous evidence of a within-family association between the PGI and BMI. Even though the family-based design provides strong support for a causal interpretation, this does *not* imply that the SNPs in the PGI must be influencing BMI through some narrowly physiological mechanism. In principle, the sibling differences in BMI could arise because of sibling differences in genes that influence the proneness to eat sweets, exercise habits, or myriad other behaviors with downstream effects on BMI. PGIs for seemingly "biological" phenotypes can thus have a substantial behavioral component. A PGI for lung health may similarly derive predictive power from SNPs that influence lung health very indirectly, through smoking habits[29,30].
Second and relatedly, it is therefore a fallacy to assume that any genetic sources of heterogeneity captured by a PGI are immutable—or even at least harder to modify than environmental sources of heterogeneity. Indeed, the possibility of identifying modifiable mechanisms through which PGIs exert some of their effects motivates some of the research using PGIs[31,32]. To continue the BMI example, the widespread replacement of sugar by low-calorie sweeteners or better behavioral tools for avoiding temptation could eliminate or reduce the effect of the PGI on BMI. Because of these issues, we urge researchers to avoid describing PGIs as "genetic endowments" or other terms that may, however inadvertently, promote the common misunderstanding that genes are a resource that is easily separable from choices made in light of that resource.
Third, because the additive genetic factor is defined conditional on the GWAS phenotype, population, and environment, the same PGI may have different predictive power in different samples if there are differences in the phenotype measure, population sampled, the sampling methodology, or the environmental context. For example, the research participants from the UKB were recruited through the mail and had a 5.5% response rate. Those that responded to the recruitment mailers were more healthy and more educated than the UK population as a whole[33,34]. Because UKB participants make up a large fraction of the discovery sample for many phenotypes, it may be that the PGI from this Repository does not correspond to a PGI that would be produced from a representative sample or a sample of individuals not from the UK.
